# In-silico development of a method for the selection of optimal enzymes using L-asparaginase II against Acute Lymphoblastic Leukemia as an example

**DOI:** 10.1101/2020.10.13.337097

**Authors:** Adesh Baral, Ritesh Gorkhali, Amit Basnet, Shubham Koirala, Hitesh K. Bhattarai

**Affiliations:** Department of Biotechnology, Kathmandu University, Dhulikhel, Nepal

**Keywords:** L-Asparaginase II, Acute Lymphoblastic Leukemia, Enzyme kinetics, Binding affinity, Homology modelling, Docking

## Abstract

L-Asparaginase II (asnB), a periplasmic protein, commercially extracted from *E. coli* and *Erwinia*, is often used to treat Acute Lymphoblastic Leukemia. L-Asparaginase is an enzyme that converts L-asparagine to aspartic acid and ammonia. Cancer cells are dependent on asparagine from other sources for growth and when these cells are deprived of asparagine by the action of the enzyme the cancer cells selectively die. Questions remain as to whether asnB from *E. coli* and *Erwinia* is the best asparaginase as they have many side-effects. asnB with the lowest Michaelis constant (Km) (most potent), and with the lowest immunogenicity is considered the most optimal enzyme. In this paper asnB sequence of *E. coli* was used to search for homologous proteins in different bacterial and archaeal phyla and a maximum likelihood phylogenetic tree was constructed. The sequences that are most distant from *E. coli* and *Erwinia* were considered best candidates in terms of immunogenicity and were chosen for further processing. The structures of these proteins were built by homology modeling and asparagine was docked with these proteins to calculate the binding energy. asnBs from *Streptomyces griseus*, *Streptomyces venezuelae* and *Streptomyces collinus* were found to have the highest binding energy i.e. −5.3 kcal/mol, −5.2 kcal/mol, and −5.3 kcal/mol respectively (Higher than the *E.coli* and *Erwinia* asnBs) and were predicted to have the lowest Kms as we found that there is an inverse relationship between binding energy and Km. Besides predicting the most optimal asparaginase, this technique can also be used to predict the most optimal enzymes where the substrate is known and the structure of one of the homologs is solved.

## 1. INTRODUCTION

Acute Lymphoblastic Leukemia (ALL) is a malignant cancer of the white blood cells characterized by uncontrolled overproduction and accumulation of lymphoid progenitor cells(“Childhood Acute Lymphoblastic Leukemia Treatment (PDQ^®^)–Patient Version - National Cancer Institute,” n.d.). It is most common among children that compromise 80% of the world wide ALL occurrences, although some cases in adults are also seen. It is equally life-threatening to in both cases. In the US, ALL is estimated to have a frequency of 1.7 cases per 100,000 people(Baljevic et al., 2016). In 2015 alone 111,000 deaths were reported out of 876,000 cases globally(Vos et al., 2016). Thus, a substantial potential market exists for new and improved therapies to ALL.

L asparaginase is a non-human enzyme of often bacterial origin. It belongs to an amidase group that hydrolyses the amide bond in L-asparagine to L-aspartic acid and ammonia(Kumar and Verma, 2012). It is an effective anti-neoplastic agent and is often used in conjugation with chemotherapy for ALL treatment. Experiments with guinea pig serum, which is a rich source of L-asparaginase(Oliver, 2013) show that it can inhibit the growth of transplantable lymphoblastic tumors in mice and rats along with radiation-induced leukemia in mice(John G. Kidd, 1953).

Normal cells require L-asparagine as an amino acid for the synthesis of proteins. A natural diet like vegetables is one of the sources of L-asparagine for the body. It is not classified as an essential amino acid as it is naturally synthesized by the body through a pathway involving the enzyme L-asparagine synthase which coverts aspartic acid and glutamic acid into L-asparagine(Savitri et al., 2003). Neoplastic cells like ALL cells lack this enzyme and therefore are not able to produce L-asparagine on their own(Keating et al., 1993). This leaves them dependent on L-asparagine from outside sources like the serum where it is pooled from diet and from normal cells. This provides the basis for the use of L-asparaginase as a therapeutic agent against ALL, the intent being to deplete the local circulating pools of L-asparagine in the blood serum thus starving the cancer cells of the amino acid and causing cell death.

L-asparaginase is produced by a wide variety of organisms and can be classified into several families. The ones of therapeutic interest can consist of two enzymes called L-asparaginase of two closely related families named L-asparaginase I and L-asparaginase II. L-asparaginase I, referred also as asnA, is a low-affinity enzyme found in the cytoplasm and is constitutively produced by the organism. L-asparaginase II, referred to as asnB, on the other hand, is a high-affinity periplasmic enzyme expressed during anaerobiosis. Its expression is dependent on aeration, carbon source, and amino acid availability(Jennings and Beacham, 1990).

Extracellular L-asparaginase accumulates in the culture broth and thus is most favorable for extraction and downstream processing for commercial production(Amena et al., 2010). The most commercial form of therapeutic L-asparaginase is extracted from *E. coli* and *Erwinia* species. They secrete the enzyme into the periplasmic space between the plasma membrane and the cell envelope(Cedar and Schwartz, 1968). The enzyme is extracted by lysis of the cells which brings the enzyme along with inner cell contents into the culture medium. It is usually purified using fractionation with ammonia sulfate.

However, the commercially available L-asparaginase has several drawbacks. L-asparaginase from *E. coli* and *Erwinia* is known to show immunogenic and allergic reactions. Most therapeutic use of L-asparaginase has shown toxicity(Duval et al., 2002). Toxicity of L-asparaginase can be attributed to lower activity of the enzyme to L-asparagine and higher activity to glutamine. Thus, the decrease in glutamine levels in the normal cells causes an allergic reaction(Campbell, H. A., Mashburn, L. T., Boyse, E. A. and Old, 2014). Another problem with the currently available L-asparaginase is the immunological response. The body recognizes the enzyme as being foreign and thus mounts an immune response against the enzyme which can range from a mild allergic reaction to anaphylactic shock(Moola et al., 1994).

The Michaelis constant (Km) is a value for the substrate concentration at which the reaction rate is half of the maximum reaction rate. A lower Km suggests that the enzyme can reach half the maximum reaction rate at lower substrate concentrations. One can interpret this to mean that enzymes with lower Km have greater activity towards that substrate. An enzyme with greater activity towards L-asparagine can be expected to show fewer undesirable effects as it will have a lower activity to unintended substrates(Johnson and Goody, 2011). Another useful metric for the measurement of enzyme activity is k_cat_ or the turn over number. It gives the number of substrates converted to a product by a single molecule of enzyme per unit time. The turn over number signifies the rate at which a substrate is catalyzed by the enzyme(Johnson, 2013).

Catalysis is based on binding energy that lowers the activation energy and to overcome the unfavorable entropic requirements needed for the correct orientation of the catalyst and reactants brought together for reaction(Hansen and Raines, 1990). Binding energy is the energy released when a substrate forms weak bonds with the enzyme active site. Binding energy is measured as the free energy (Delta G). Gibbs free energy is defined as “a thermodynamic potential that measures the capacity of a thermodynamic system to do maximum or reversible work at a constant temperature and pressure (isothermal, isobaric), is one of the most important thermodynamic quantities for the characterization of the driving forces”(Gibbs, 1873).

Experimental calculation of this energy is difficult and cumbersome. Thus, experimental screening techniques for a lead compound for drug candidates are still very expensive and slow despite several advances in automation and parallelization of the process. A more efficient method would be to screen a large library of small molecules *in silico* before shortlisting a small group for experimental verification. The availability of large volumes of experimental data on the three-dimensional structure of the enzymes and their substrates allows us to analyses their interaction. Docking is one of these *in silico* methods where rigid body interaction of contact surfaces of the ligand/small molecules and the target protein is determined using computational methods. Combinatorial methods are used to account for the ligand conformational flexibility and various energy functions are used to calculate energetics of the interaction. Docking is typically used to screen for potential lead compound candidates from a large library of small molecules based on their binding energy and other parameters to the target protein. Those compounds with greater binding energy to the protein are seen as potential inhibitors and thus considered to lead for developing drugs of therapeutic value(Lengauer and Rarey, 1996). However, in L-asparaginase based therapy of ALL, the enzyme itself is used as a therapeutic agent while the substrate, L-asparagine, is the target compound. Our goal in this research is to find a better enzyme candidate with more favorable interaction with our target compound. Thus, our use of docking in this research is different from the standard use of the docking method. We used docking to screen a collection of L-asparaginase enzyme from different organisms and select a suitable enzyme based on its binding energy to L-asparagine.

The *E. coli* L-asparaginase II has a functional form in a homo-tetramer having the molecular mass from 140 to 160 kDa. The monomers are 330 amino acid long and have two distinct domains. One is the larger N-terminal domain and the other is the smaller C-terminal domain. The two domains are connected by a 20-residue linker. The functional form of the enzyme is thought to contains five active sites(Khushoo et al., 2004).

Homology modelling is a technique used to generate a model from an amino acid sequence-based on a template of a three-dimensional structure of a closely related protein obtained via experimental data. It uses comparative protein structure modeling where the template and the query sequences are aligned, and the query’s structure is predicted. According to Narayanan and Co. in Comparative Protein Structure Modeling Using Modeller(Eswar et al., 2016), it has the following four major steps: fold assignment, which identifies similarity between the target and at least one known template structure; alignment of the target sequence and the template(s); building a model based on the alignment with the chosen template(s), and predicting model errors. We have used MODELLER 9.22 to model L-asparaginase sequence from the organisms that were selected, using the *E. coli* L-asparaginase II (PDB Id.: 1nns) as a template for generating all of them.

*E. coli* and *Erwinia* L-asparaginases, the two commercially available forms of the therapeutic enzymes, suffer from deficiencies in the above-mentioned parameters. Thus, they show unsatisfactory results and side effects. In this research, we hope to find a better L-asparaginase from a different host organism for the commercial production of this therapeutic enzyme. We hypothesize that a host whose L-asparaginase amino acid sequence is distinct from that of the currently used organisms can be assumed to have markedly different properties. We can screen such a family/genus of host organisms and hope to find L-asparaginase that displays kinetic and binding properties that decrease the chances of immunogenic and allergic reactions making it more favorable for therapeutic use. We have used a phylogenetic tree-based approach to find such host organisms. A phylogenetic tree is an important bioinformatics tool that allows us to analyze the sequences of proteins/DNA/RNA to find the historical and evolutionary relationship between the sequences. The nodes of a tree can be given values as support values for its reliability. These are called bootstrap values which give the expectation of that particular node in the many alternate tree generated by re-runs of the same sequence data set(Rokas, 2011). Many algorithms for tree construction exist. Here we have used the Maximum Likelihood (ML) algorithm in the MEGA bioinformatics tool to construct, bootstrap, and analyze our tree. The tree was used to look for hosts with evolutionarily distant L-asparaginase sequences which can be screened for desired properties using docking tools.

## 2. Methods

### 2.1 Phylogenetic Tree Construction

In order to construct a phylogenetic tree, we retrieved the L-asparaginase B (asnB) protein sequence of *Escherichia coli k12* strain from the Uniprot (https://www.uniprot.org/uniprot/P00805) (UniProtKB - P00805 (ASPG2_ECOLI). Microorganisms which are capable of producing the asnB based on the previous literature (Peterson and Ciegler, 1969),(Cachumba et al., 2016),(Jha et al., 2012),(El-Naggar et al., 2014), were searched by doing blastp in NCBI (https://blast.ncbi.nlm.nih.gov/Blast.cgi?PAGE=Proteins). The organisms with percentage identity greater or equal to 30% were selected. The genomes of two types of organisms were searched for the presence of asnB. The first group of organisms were already characterized for the production of asnB protein. The other group of organisms included bacteria and archaea from various phyla(Petitjean et al., 2014) that represented the entire tree of life. Hundred and one sequences were retrieved after searching for asnB sequence in organisms given by literature. Organisms having more than one asnB sequences were also retrieved and labeled as Genus species 1, 2, or 3. Then the phylogenetic tree was constructed in Mega-X software (https://www.megasoftware.net/), in which the alignment was done by Muscle. The following criteria were used to run a tree Statistical method: Maximum likelihood, Test of Phylogeny: Bootstrap method, Substitution Type: Amino acid, Model/method: WAG model, Rates among sites: Gamma Distributed With Invariant Sites (G+ I), No of Discrete Gamma Categories: 5, Gaps/Missing Data Treatment: Partial deletion, Site Coverage Cutoff(%):95% ML Heuristic Method: Nearest-Neighbor-Interchange, Initial Tree for ML: Make initial tree automatically(Default-NJ/BioNJ), Branch Swap Filter: None, Number of thread: 3(Hall, 2013).

### 2.2 Homology Modelling

The organisms, which were distantly placed in the phylogenetic tree with respect to *E.coli* and *Erwinia*, were chosen and organisms whose enzymes were characterized in literature were also chosen. To carry out homology modelling MODELLER 9.22 was used. The selected organism’s asnB sequence was used as the query while *E.coli k12* asnB (“1nns”)(https://www.rcsb.org/structure/1NNS) with a resolution of 1.95 Å was used as the reference template. For each organism, the structure with the lowest DOPE or SOAP assessment score and with the highest GA341 assessment score was selected(Šali, 2013). Each proteins models were then checked for protein structure stereochemistry including Ramachandran plot and Psi/Phi angles using PROCHECK. Further verification was done using WHATCHECK and ProSA-web (https://prosa.services.came.sbg.ac.at/prosa.php).

### 2.3 Active site prediction

After the validation of the model, active sites for each protein were determined using PyMol software (https://pymol.org/2/). The models built were superimposed to the 1nns structure and then by aligning both model and 1nns sequences, the active site with reference to 1nns active site was predicted. The active site of 1nns for L-asparagine is T(12), S(58), Q(59), T(89), D(90)(Sanches et al., 2003).

### 2.4 Molecular Docking studies

Docking of ligands, L-asparagine (derived from PubChem website) with enzymes L-asparaginases (distant proteins from *E. coli* and *Erwinia* and enzymes with measured Km value) was performed by using auto dock vina (http://vina.scripps.edu/) conjugated with PyRx software (https://pyrx.sourceforge.io/). The auto dock tool’s graphic interface is used for the preparation of all the proteins (enzymes). Proteins were prepared by removing water, adding polar hydrogen, merging non-polar, and adding Kollman charge. In the case of ligand, L-asparagine was retrieved from the PubChem(Compound CID: 6267), Molecular formula: C4H8N2O3, Molecular weight: 132.12 g/mol(https://pubchem.ncbi.nlm.nih.gov/compound/6267). Energy minimization was done by the Universal force field (UFF) using Open babel software (http://openbabel.org/wiki/Main_Page) conjugated with PyRx. GPF (grid parameter file) and DPF (docking parameter file) were set and the grid points for auto grid calculations were set as 25 × 25 × 25 Å with the active site residues in the middle of the grid box. The algorithm used in the overall process was the Lamarckian genetic algorithm which was used to calculate protein-fixed, ligand-flexible calculations(Sippl, 1993).

Distant organisms’ asnBs with the best binding energies were selected. The docked protein and ligand file was run on ligPlot+ software (https://www.ebi.ac.uk/thornton-srv/software/LIGPLOT/) for viewing the interacting atoms between ligands and proteins.

## 3. Results

### 3.1 Deductions from the phylogenetic tree

A list of asparaginase producing organisms were compiled from the literature. Asparaginase II (asnB) homologs of these organisms were searched by protein blasting asnB from *E.coli* against the non-redundant protein database of these organisms in NCBI. The organisms whose genomes are not sequenced are not used in this study. Additionally, the protein database of a wide variety of bacteria and archaea from different phylum were searched for the presence of asnB. The two lists were compiled to make up our list of a wide range of asnBs. Maximum Likelihood (ML) phylogenetic tree was drawn for these proteins using Mega X software using the parameters described in the methods section. The resulting tree is shown in Figure 1. The phylum of bacteria, archaea, and fungi to which the proteins belong to is labeled on the right. Unlike most other proteins for which similar trees were drawn, there were minimal proteins from the same phylum that lay next to each other in the tree. When a similar tree was drawn for Ku protein in bacteria and beta clamp for bacteria, proteins from the same phylum tended to cluster together in the tree (unpublished data). Although some clustering is found for asnB tree, proteins from the same phylum are distributed throughout the tree, indicating extensive horizontal gene transfer. Among the list of asnBs that we have collected, the largest number of proteins comes from proteobacteria (alpha, beta, gamma, delta, and epsilon).

**Figure 1.**
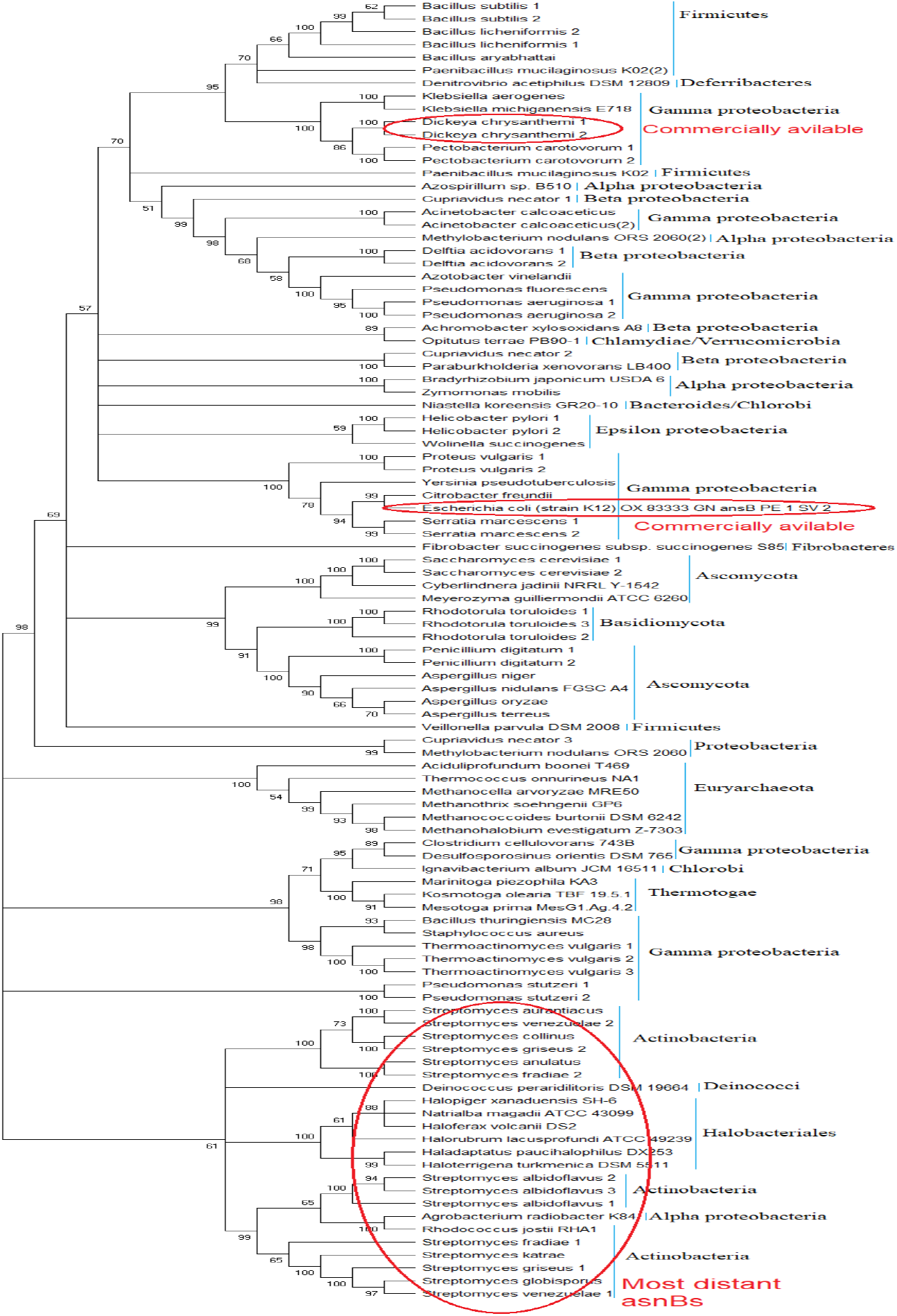
Phylogenetic tree of the total hundred and one sequences of asnBs using the Maximum likelihood method. Top and the middle portion of the tree under the red oval showing the organisms which are currently used for the commercial production of asnBs for the treatment of ALL. And the bottom portion of the tree showing the organisms which are most distant to the E.coli,(mostly Actinobacteria) and their enzyme activity yet to discover.

Besides predicting the origin and history of asparaginases, the tree is also useful in predicting which of the asnBs are closely related by evaluating which lie close together and which lie further apart. From the tree, the most common commercially used asnB from *E. coli* lies somewhere in the center. The other commercially used asnB from *Erwinia* (nowadays called *Dickeya chrysanthami*) lies at the bottom of the tree. The asnBs that are most distant from these two commercially available asparaginases, and hence least likely to give an immunogenic reaction when these two give an immunogenic reaction, lie at the top of the tree. These have been labeled in the figure (Fig.1). Most of them lie in the Streptomyces genus and some are from archaea.

### 3.2. Homology modeling and verification

For homology modeling MODELLER 9.22 software was used in which five models were built for each protein among which the model with the lowest discrete optimized protein energy (DOPE) was selected. This software uses an inbuilt DOPE function to access the quality of all the model which are made. The model which were selected according to the lowest DOPE scores were validated using Ramachandran Plot. Ramachandran Plot of the three best organisms which lie in distant to the *E.coli* and have a better binding affinity toward L-asparagine than *E.coli* as well as *Dickeya chrysanthami* are shown in figure 2. The plot shows that 94.5% of residues in most favored regions, 4.4% in additional allowed regions, 0.4% residues in generously allowed regions, and 0.7% residues in disallowed regions for *Streptomyces collinus*(Fig. 2a), 86% of residues in most favored regions, 10.5% in additional allowed regions, 2.3% residues in generously allowed regions, 0.7% residues in disallowed regions for *Streptomyces griseus 1*(Fig. 2b) and 90.7% of residues in most favored regions, 7.8% in additional allowed regions, 0.7% residues in generously allowed regions, 0.7% residues in disallowed regions for *Streptomyces venezuelae 2*(Fig. 2c). More than 99% residues are in allowed region given by Ramachandran plot indicates a very good model. Furthermore, the Ramachandran Z-score calculated by WHATCHECK −0.245, - 1.024, −0.830 for *S. collinus, S. griseus 1, S.venezuelae 2* respectively falls on the accepted region(Sousa et al., 2006) and allowed by the WHATCHECK. The structures were finally validated using ProSA-web server. This server gives the z-score which indicates the overall model quality and measures the deviation of the total energy of the structure with respect to an energy distribution derived from random conformations(Hooft et al., 1997).The z-score given by the server −9.44, −7.88, −9.07 for *S. collinus, S. griseus 1, S.venezuelae 2* falls inside the range of plot (black dot) that contains the z-score of all the experimentally determined protein in the PDB (X-ray, NMR) (Figs. 3a,4a,5a). In the energy plot (Figs. 3b,4b,5b) which indicates the local model quality by plotting energy as the function of the amino acid sequence. Generally, the portion in the positive region of the plot indicates the erroneous part of the structure. We can conclude from the plot that the structure is feasible or accepted as overall residue energies fall under the negative part of the plot. The colored 3D structure of the proteins (Figs. 3c,4c,5c) shows that the portion in red color is of high energy, and the portions with the blue color are of low energy(Wiederstein and Sippl, 2007). Validation of all other structures used in the experiment is in the supplement data. Most of the active site residues are conserved in every model made by the MODELLER 9.22 in reference to the 1nns structure also signifies the good models were made during the process and can proceed toward the docking (Table 1).

**Figure 2:**
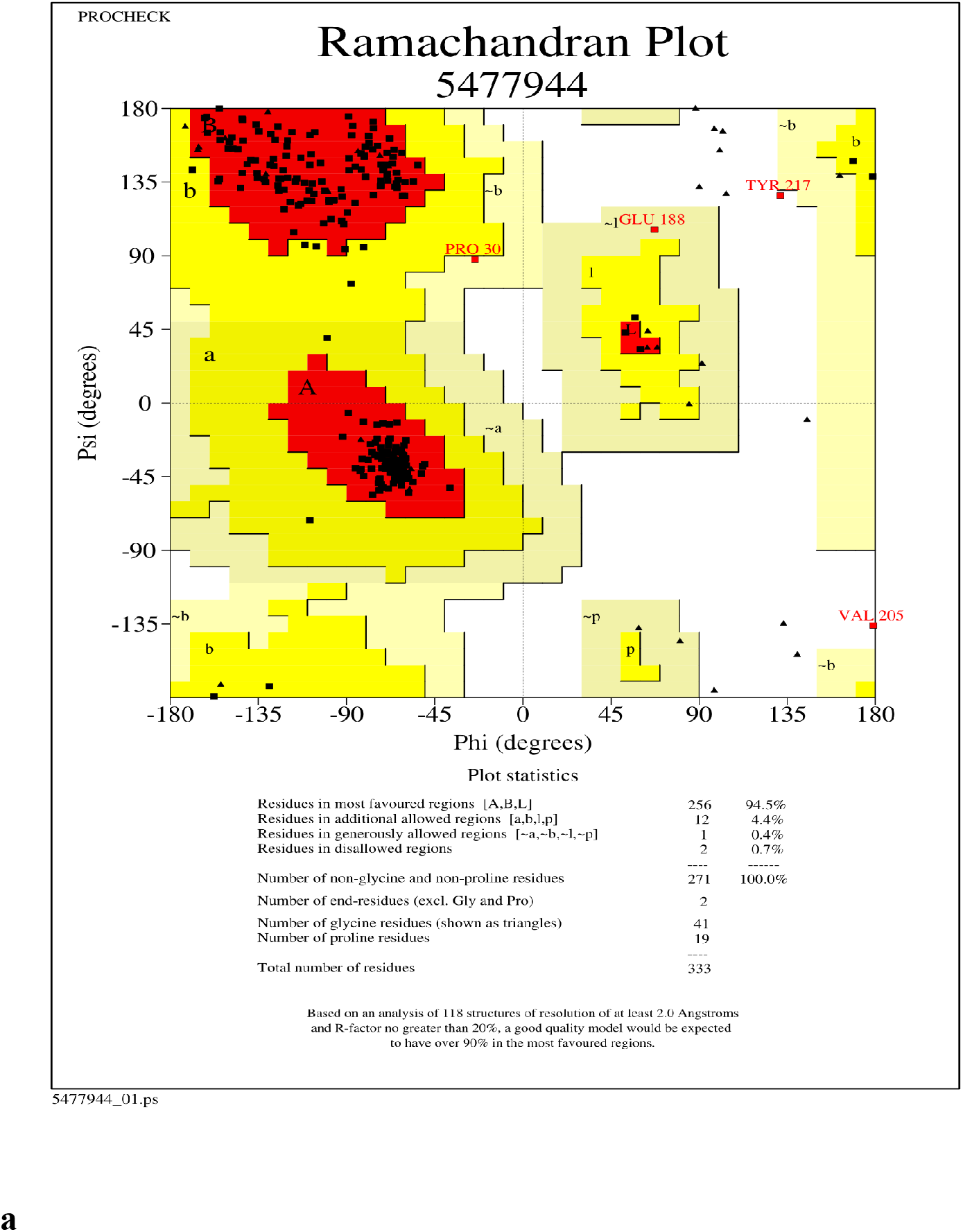
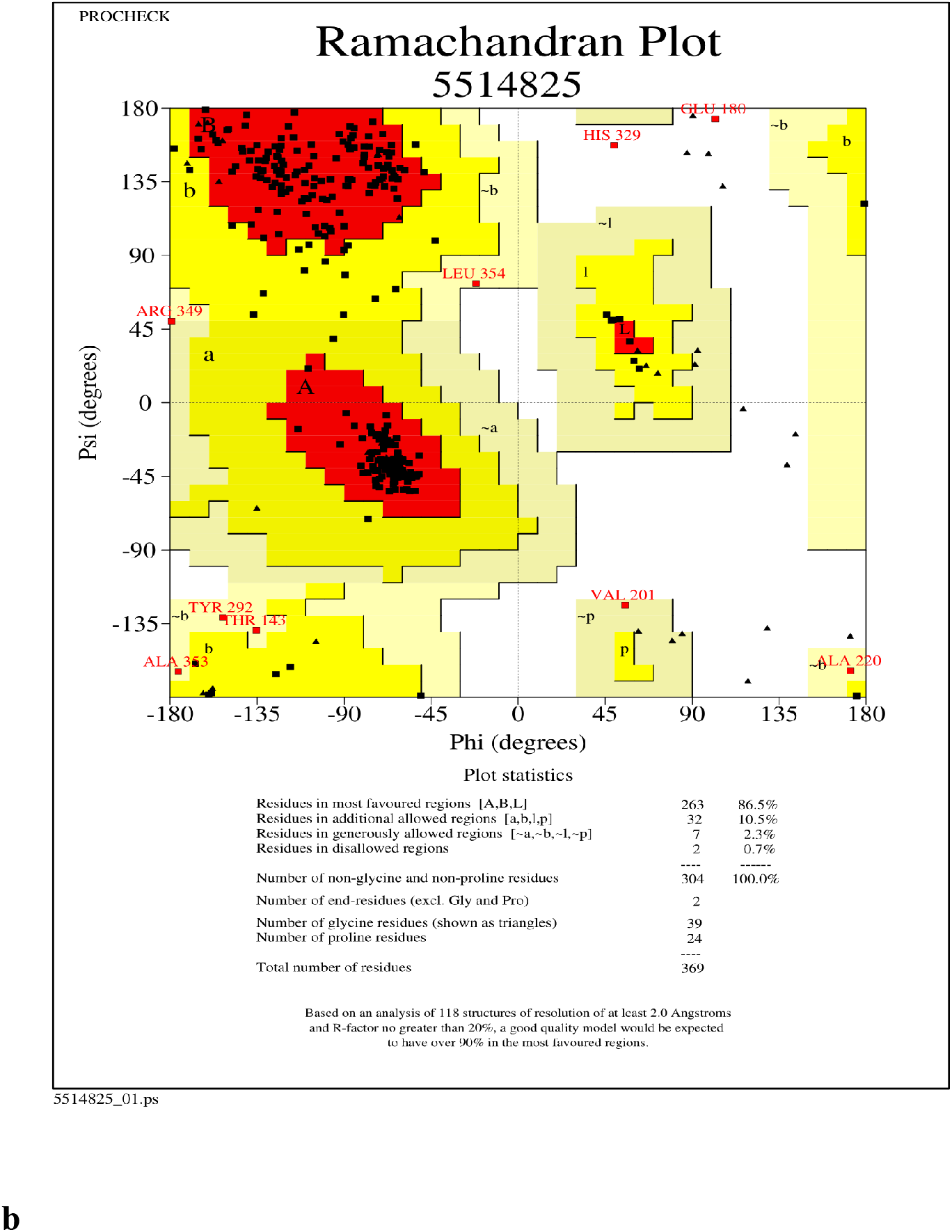
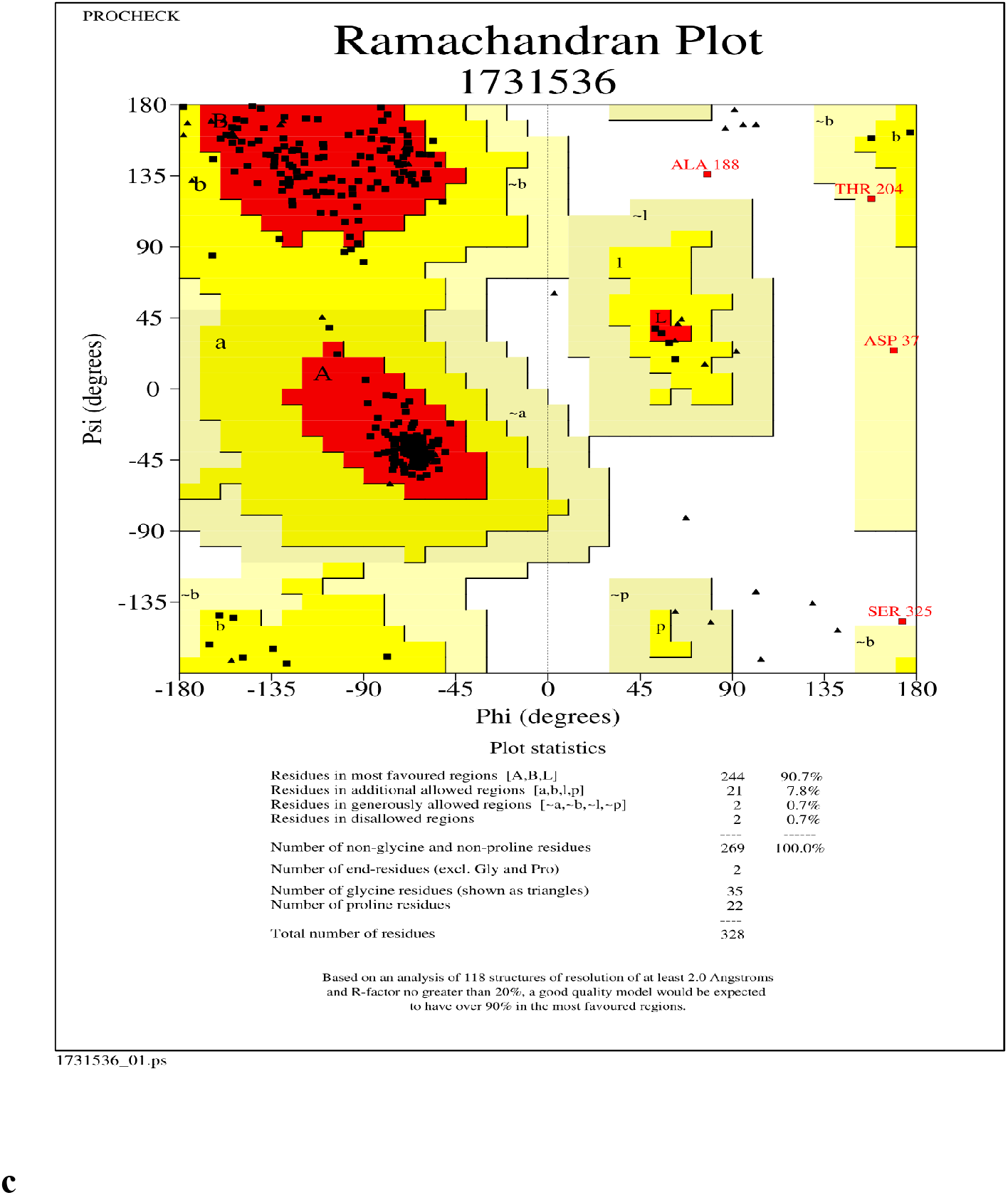
Ramachandran Plot of model protein asnB of **(a)***Streptomyces collinus*, **(b)***Streptomyces griseus 1* and **(c)***Streptomyces venezuelae 2.* The Ramachandran plot shows the phi-psi torsion angles for all residues (black cubic box) in the structure (except those at the chain termini). Glycine residues are separately identified by triangles as these are not restricted to the regions of the plot appropriate to the other sidechain types. The darkest red area indicates the “core” regions representing the most favorable combinations of phi-psi values. The regions are labeled as follows: **A**-Core alpha, **L**- Core left-handed alpha, **a**-Allowed alpha, **l** - Allowed left-handed alpha, **~a** - Generous alpha, **~l** - Generous left-handed alpha, **B** - Core beta,**p** - Allowed epsilon, **b**-Allowed beta,**~p**-Generous epsilon, **~b**-Generous beta.

**Figure 3.**
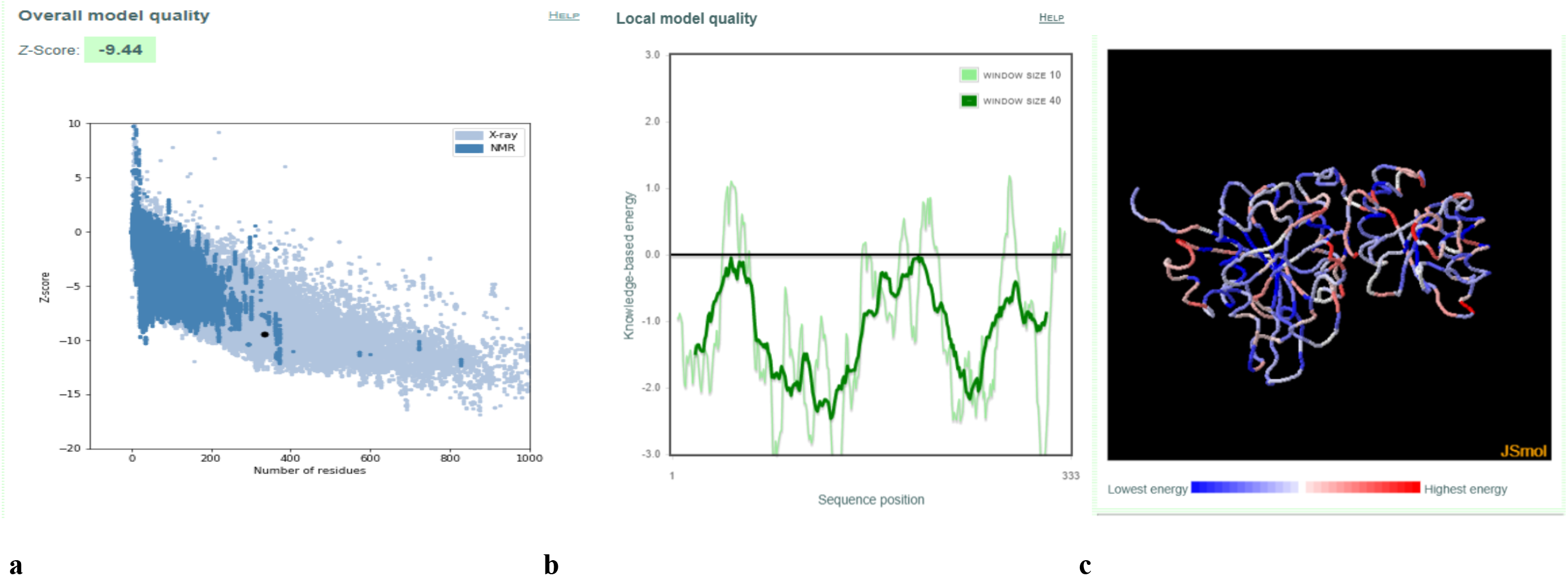
Validation of model protein structure of asnB of *Streptomyces collinus* using ProSA-web server **(a)**ProSA-web z-scores of all protein chains in PDB determined by X-ray crystallography (light blue) or NMR spectroscopy (dark blue) with respect to their length. The black dot in the plot indicates that the model protein structure falls inside the range of the plot that contains the z-score of all the experimentally determined protein in the PDB. The plot shows only chains with less than 1000 residues and a z-score 10. The z-scores of model proteins are highlighted as large dots. **(b)**Energy plot of model protein which indicates the local model quality by plotting energy as the function of the amino acid sequence. Generally, the portion in the positive region of the plot indicates the erroneous part of the structure **(c)**Residues are colored from blue to red in the order of increasing residue energy.

**Figure 4.**
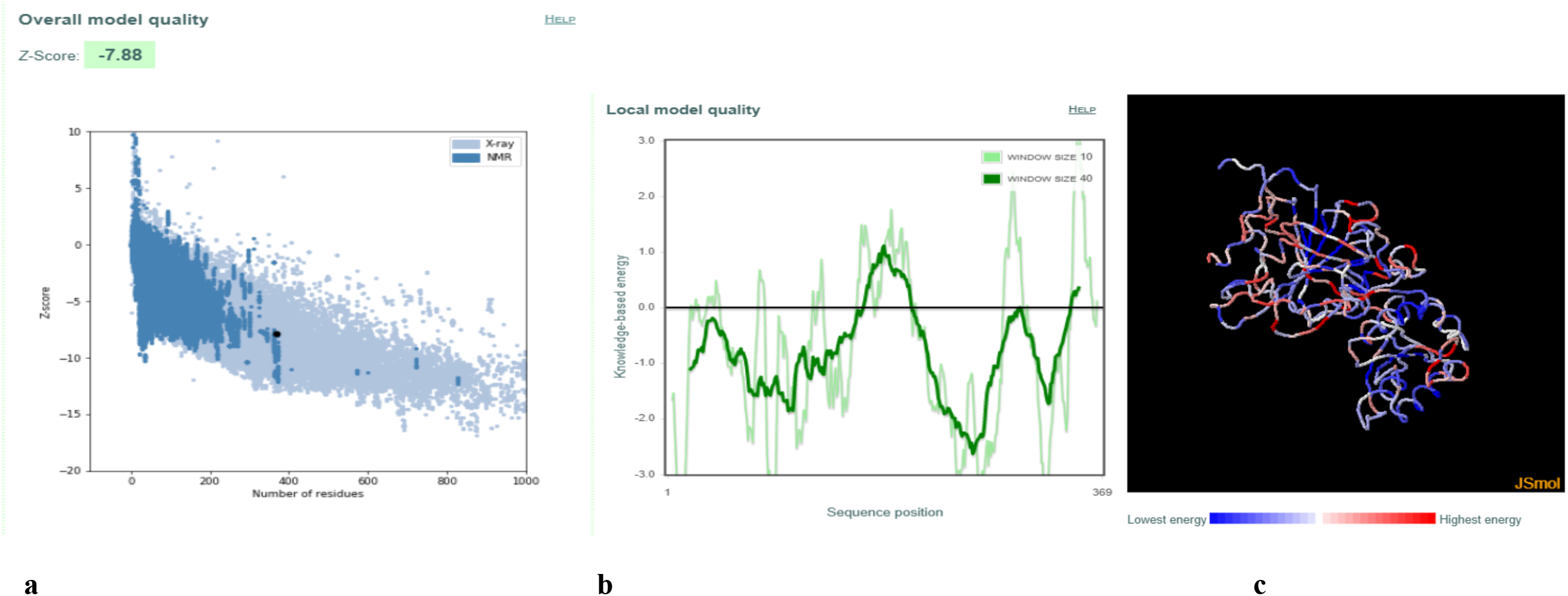
Validation of model protein structure of asnB of *Streptomyces griseus 1* **(a)**ProSA-web z-scores of all protein chains in PDB determined by X-ray crystallography (light blue) or NMR spectroscopy (dark blue) with respect to their length. The black dot in the plot indicates that the model protein structure falls inside the range of the plot that contains the z-score of all the experimentally determined protein in the PDB. The plot shows only chains with less than 1000 residues and a z-score 10. The z-scores of model proteins are highlighted as large dots. **(b)**Energy plot of model protein which indicates the local model quality by plotting energy as the function of the amino acid sequence. Generally, the portion in the positive region of the plot indicates the erroneous part of the structure **(c)**Residues are colored from blue to red in the order of increasing residue energy.

**Figure 5.**
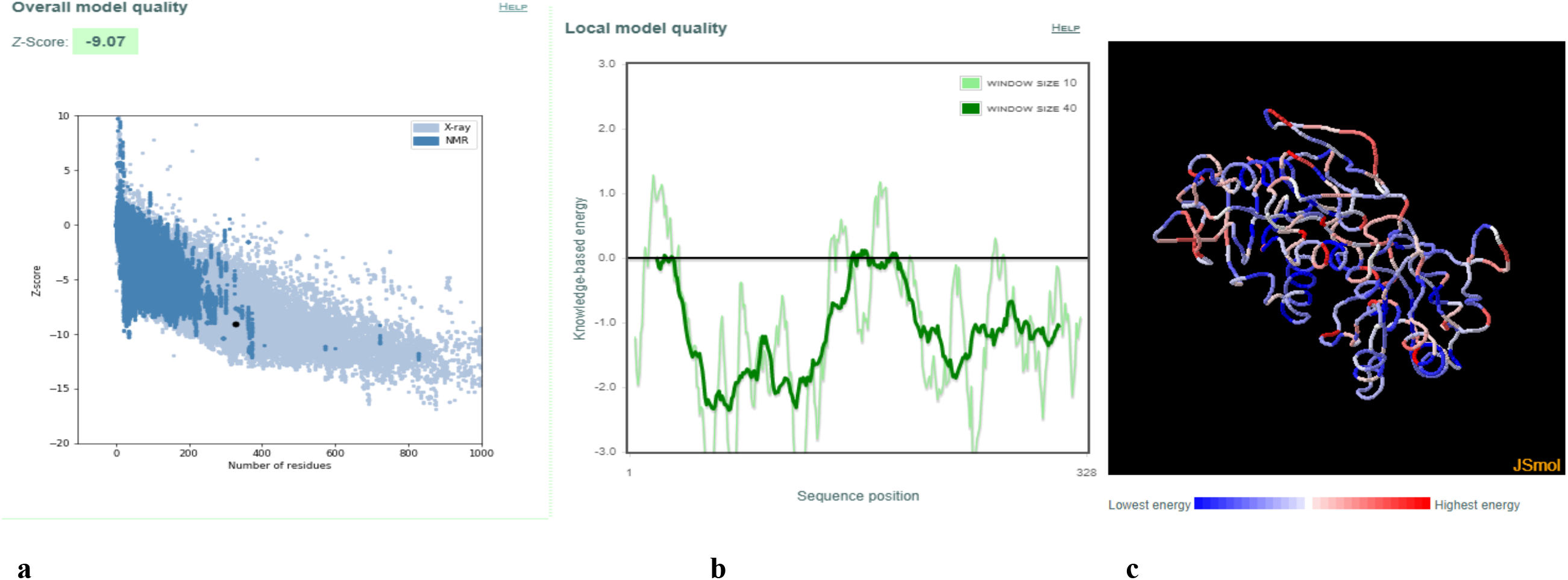
Validation of model protein structure of asnB of *Streptomyces venezuelae 2* using ProSA-web server **(a)**ProSA-web z-scores of all protein chains in PDB determined by X-ray crystallography (light blue) or NMR spectroscopy (dark blue) with respect to their length. The black dot in the plot indicates that the model protein structure falls inside the range of the plot that contains the z-score of all the experimentally determined protein in the PDB. The plot shows only chains with less than 1000 residues and a z-score 10. The z-scores of model proteins are highlighted as large dots. **(b)**Energy plot of model protein which indicates the local model quality by plotting energy as the function of the amino acid sequence. Generally, the portion in the positive region of the plot indicates the erroneous part of the structure **(c)**Residues are colored from blue to red in the order of increasing residue energy.

**TABLE 1:** Predicted active sites of proteins of organisms that were distant to the *E.coli* and organisms whose Km have has been determined experimentally mentioned in research paper elsewhere. Five amino acids are conserved which have been termed pentad in the paper. The letter represents the amino acid involved in the active site, the number in the bracket represents the position of the amino acid. When no amino acid homology was found, the site is left blank with a dash.

| Organisms | Predicted Active Site Residues |
| --- | --- |
| *Escherichia coli* | T(34), S(80), Q(81), T(111), D(112) |
| *Streptomyces globisporus* | I(12), S(61), S(62), T(94), D(95) |
| *Streptomyces venezuelae 1* | I(12), S(61), S(62), T(94), D(95) |
| *Streptomyces griseus 1* | T(20), S(61), S(62), T(94), D(95) |
| *Streptomyces katrae* | T(12), S(53), P(54), T(86), D(87) |
| *Streptomyces fradiae* | A(12), G(43), A(44), T(75), D(76) |
| *Streptomyces albidoflavus 1* | T(12), M(62), R(63), T(94), D(95) |
| *Streptomyces albidoflavus 2* | T(12), M(62), R(63), T(94), D(95) |
| *Streptomyces albidoflavus 3* | T(12), R(63), L(64), T(94), D(95) |
| *Streptomyces fradiae 2* | T(8), S(50), Y(51), T(83), D(84) |
| *Streptomyces collinus* | T(16), S(63), L(64), T(94), D(95) |
| *Streptomyces griseus 2* | T(16), P(60), G(61), T(94), D(95) |
| *Streptomyces aurontiacus* | T(13), S(54), L(55), T(83), D(84) |
| *Streptomyces venezuelae 2* | T(12), -, -, T(79), D(80) |
| *Pectobacterium carotovorum 1* | T(34), S(81), E(82), T(114), D(115) |
| *Dickeya chrysanthami (Erwinia) 1* | T(36), S(83), E(84), T(116), D(117) |
| *Bacillus aryabhattai* | T(55), S(102), Q(103), S(135), D(136) |
| *Bacillus Licheniformis 1* | T(62), S(109), Q(110), T(142), D(143) |
| *Bacillus subtilis 1* | T(61), S(108), T(109), T(141), D(142) |
| *Delftia acidovorans 1* | T(62), S(109), E(110), T(142), D(143) |
| *Azotobacter vinelandii* | T(45), S(92), E(93), T(125), D(126) |
| *Dickeya chrysanthami (Erwinia) 2* | T(36), S(83), E(84), T(116), D(117) |
| *Helicobacter pylori* 1 | T(34), S(80), Q(81), T(113), D(114) |
| *Pseudomonas stutzeri* 1 | - , S(80), D(81), T(113), D(114) |
| *Pseudomonas stutzeri* 2 | - , S(80), D(81), T(113), D(114) |
| *Bacillus subtilis* 2 | T(61), S(108), T(109), T(141), D(142) |
| *Bacillus licheniformis* 2 | T(63), S(110), T(111), T(143), D(144) |
| *Delftia acidovorans* 2 | T(62), S(109), E(110), T(142), D(143) |
| *Helicobacter pylori* 2 | T(34), S(80), Q(81), T(113), D(114) |
| *Pectobacterium carotovorum* 2 | T(34), S(81), E(82), T(114), D(115) |

### 3.3. Active site of asnBs

Along with the 1nns structure of E. coli asnB comes the description of active site amino acid residues. Using aspartate as a surrogate for asparagine, the active sites have been predicted. For the full-length protein, the active site contains 5 amino acid residues: T(34), S(80), Q(81), T(111), D(112). These 5 residues can be called a pentad. A table with these pentad residues has been constructed for asnBs of other organisms (Table 1). Four of the five residues—T(34), S(80), T(111), and D(112)—are highly conserved across species (Table 1).

### 3.4. Km, k_cat_ and binding energies of asnBs

To further predict which of the list of asnBs would be most useful to treat ALL, binding energies were calculated using docking software. Firstly using 1nns structure of E. coli asnB, structures of unsolved asnBs were predicted using homology modeling. These structures were docked to asparagine to calculate binding energy. To evaluate if the binding energy could predict the relative efficacy of the enzymes, Km, and kcat values from the literature were tabulated alongside binding energy (Table 2). A total of 10 Km values were obtained from the literature for asnBs of different species. For the species with only one Km value—*Escherichia coli*, *Azobacter vinelandi*, *Bacillus aryabhattai*—comparison between the relationship of Km and binding energy was easy. When Km value increased, binding energy decreased. Species with the highest binding energy, *E. coli*, also had the lowest Km value. Species with the lowest binding energy, *Bacillus aryabhattai*, had the highest Km value.

**TABLE 2:** Km value, k_cat_ value (retrieved from the literature), and Binding energy (calculated by auto dock vina) of the enzyme, asnB, toward L-asparagine were tabulated against each other. For six species marked with * corresponding Km values and binding energies are known (i.e. sequence of protein used to calculate the Km experimentally and sequence of protein used to calculate the binding energy were the same). For four other species, Km value that best fits the binding energy value were randomly assigned. The six Km values are perfectly inversely correlated to binding energies. k_cat_ values demonstrate no relationship to the binding energy.

| Organism | Michaelis constant value from literature (mM) | Measured $k_{cat}$ values from literature ($s^{-1}$) | Binding affinity (kcal/mol) calculated from docking |
| --- | --- | --- | --- |
| *Bacillus licheniformis* 1 | 0.014 (Mahajan et al., 2014) | $2.68 \times 10^3$ (Mahajan et al., 2014) | -4.8 |
| *Escherichia coli* * | 0.015 (Derst et al., 2000) | $2.4 \times 10^1$ (Derst et al., 2000) | -5.1 |
| *Deftia acidovorans* 1 | 0.015 (Davidson et al., 1977) |  | -5.1 |
| *Dickeya chrysanthami* 2 * | 0.058 (Kotzia and Labrou, 2007) | $23.8 \times 10^3$ (Kotzia and Labrou, 2007) | -5.0 |
| *Azobacter vinelandi* * | 0.11 (Gaffar and Shethna, 1977) |  | -4.9 |
| *Pseudomonas stutzeri* 2 | 0.14 (Manna et al., 1995) |  | -4.9 |
| *Bacillus aryabhattai* * | 0.257 (Singh et al., 2013) |  | -4.8 |
| *Helicobacter pylori* 1* | 0.29 (Cappelletti et al., 2008) | 19.26+/-0.56 (Cappelletti et al., 2008) | -4.8 |
| *Bacillus subtilis* 1* | 0.43 (Jia et al., 2013) |  | -4.5 |
| *Pectobacterium carotovorum* 1 | 0.657 (Kumar et al., 2011) | 2.751*10<sup>3</sup> (Kumar et al., 2011) | -4.4 |
| *Dickeya chrysanthami* 1 |  |  | -4.4 |
| *Bacillus licheniformis* 2 |  |  | -4.6 |
| *Pseudomonas stutzeri* 1 |  |  | -5.1 |
| *Deftia acidovorans* 2 |  |  | -5.0 |
| *Bacillus subtilis* 2 |  |  | -5.0 |
| *Pectobacterium carotovorum* 2 |  |  | -4.7 |
| *Helicobacter pylori* 2 |  |  | -5.1 |

However, six species contained two asparaginases. From the literature, specific Km values could be assigned to specific asnBs (i.e. sequence of protein used to calculate the Km experimentally and sequence of protein used to calculate the binding energy were the same). Those asnBs are marked with * in the table. *Dickeya chrysanthami 2*, *Heliobacter pylori 1*, and *Bacillus subtilis 1* had known Km values that were assigned next to them on the table. Similarly, using docking, separate binding energies could be calculated for each asnB protein. In species where two asnBs are available the Km value measured for the species is assigned to asnB, which most closely forms an inverse relationship with the binding energy. For example, *Pseudomonas stutzeri* has two asnBs with binding energies of −5.1 Kcal/mol and −4.9 kcal/mol. Since its Km value is very high, the asnB with low binding energy was assigned this Km although this could not be verified experimentally. When all values were assigned a very clear inverse relationship between Km and binding energy emerged. The binding energies of asnB to asparagine ranged from −5.1 kcal/mol to −4.4 kcal/mol, which are relatively high values of binding in Auto dock Vina software. No relationship could be discerned for kcat value and binding energy. To be able to compare Km value to binding energies plots were drawn. A smooth curve was fitted (Fig 6).

**Figure 6:**
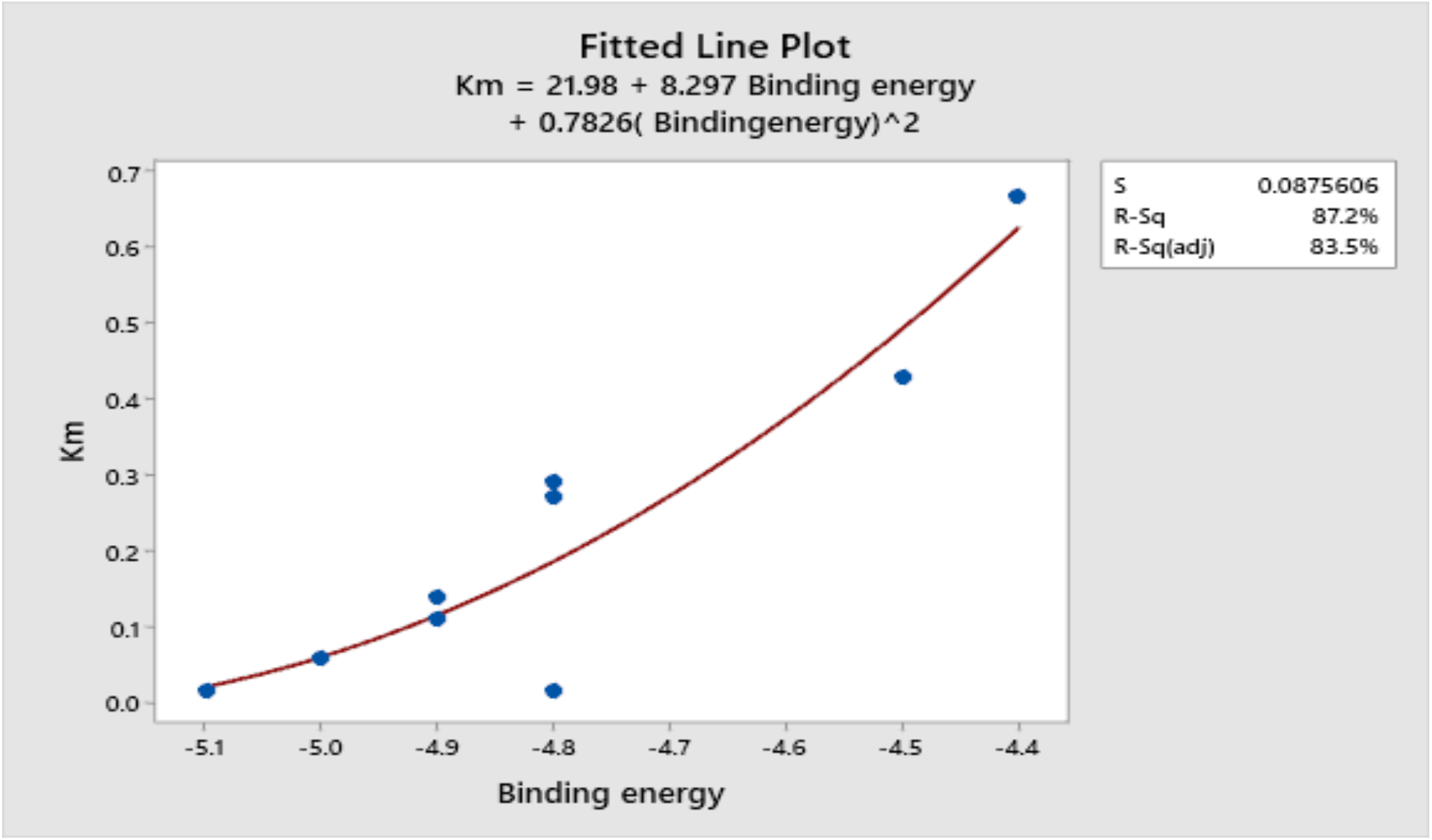
Relation between Km and Binding energy of enzyme towards L-asparagine. The fitted line plot shows that Km and binding energy are inversely proportional to each other. The more negative the binding energy less is the Km value. More negative binding energy, less Km signifies the greater affinity of enzyme toward the substrate. All the enzymes Km and Binding energy shows the inversely proportional to each other except one which is the enzyme from *Bacillus licheniformis* 1 (0.014mM Km at −4.8 kcal/mol). We were also unable to confirm that the sequence of the enzyme that was used to calculate the Km value(Mahajan et al., 2014) and the sequence of the enzyme used in this experiment were the same.

### 3.5. Finding an optimal asnBs

For 13 asnBs that are most distant from *E. coli* and *Erwinia* asparaginase, binding energies were calculated using docking (Table 3). The proteins for which binding energy were calculated are: *Streptomyces albidoflavus* 1, 2 and 3, *Streptomyces aurantiacus*, *Stereptomyces collinus*, *Streptomyces fladiae* 1 and 2, *Streptomyces globisporus*, *Streptomyces griseus* 1 and 2, *Streptomcyces katrae* and *Streptomyces venezuelae* 1 and 2. Out of these 13 proteins 3 asnBs, *Stereptomyces collinus*, Streptomyces griseus 1 and *Streptomyces venezualae* 2 showed biding energy of −5.3 kcal/mol, −5.3 kcal/mol, and 5.2 kcal/mol respectively higher than *E. coli* asnB. Docked structure are shown is figure 7. These asparaginases can be further cloned and tested for Km and k_cat_ values.

**Figure 7:**
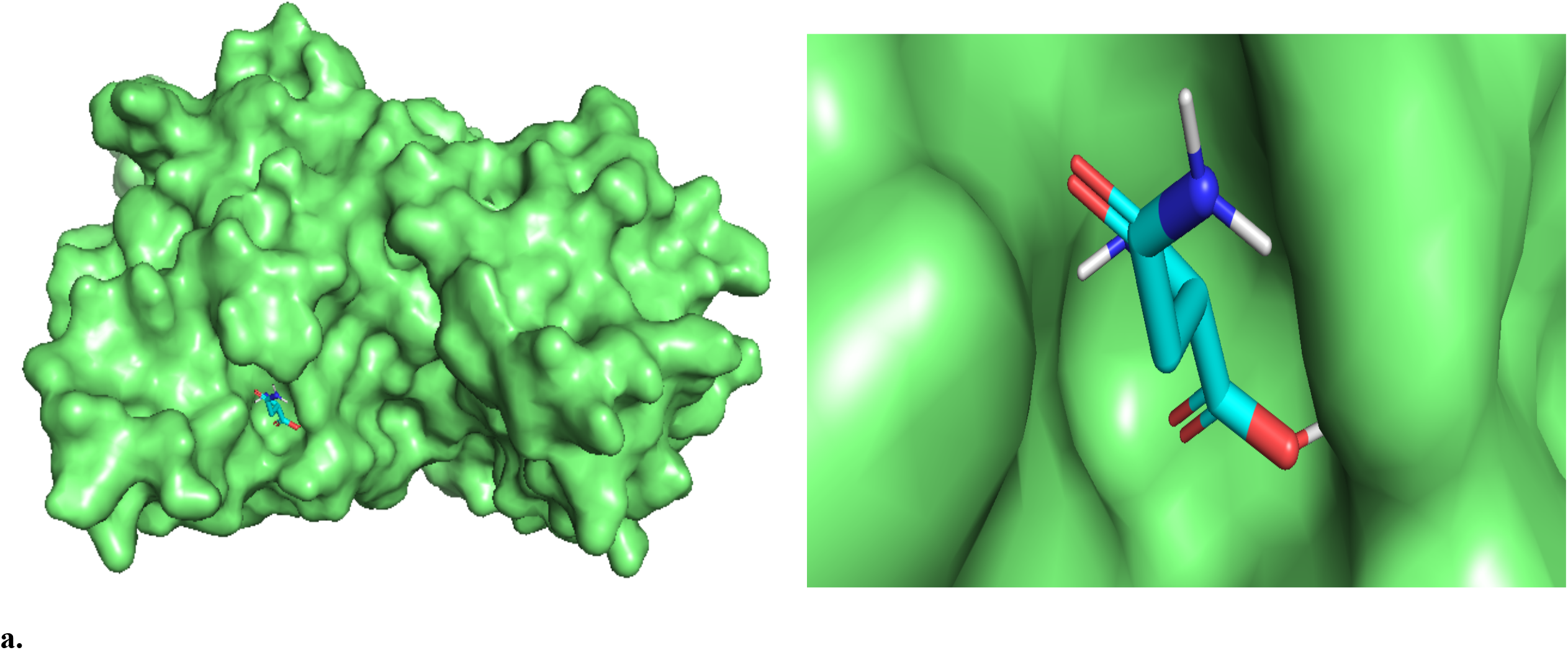
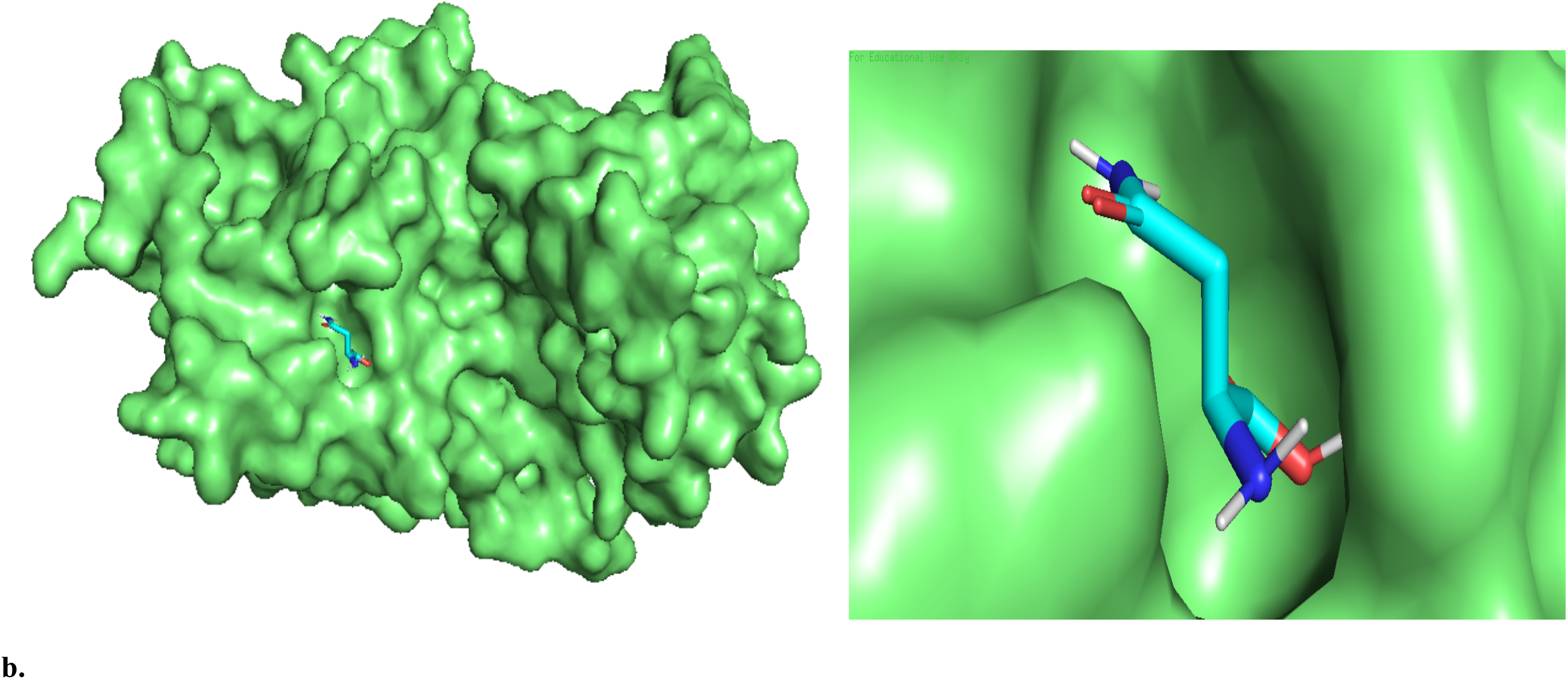
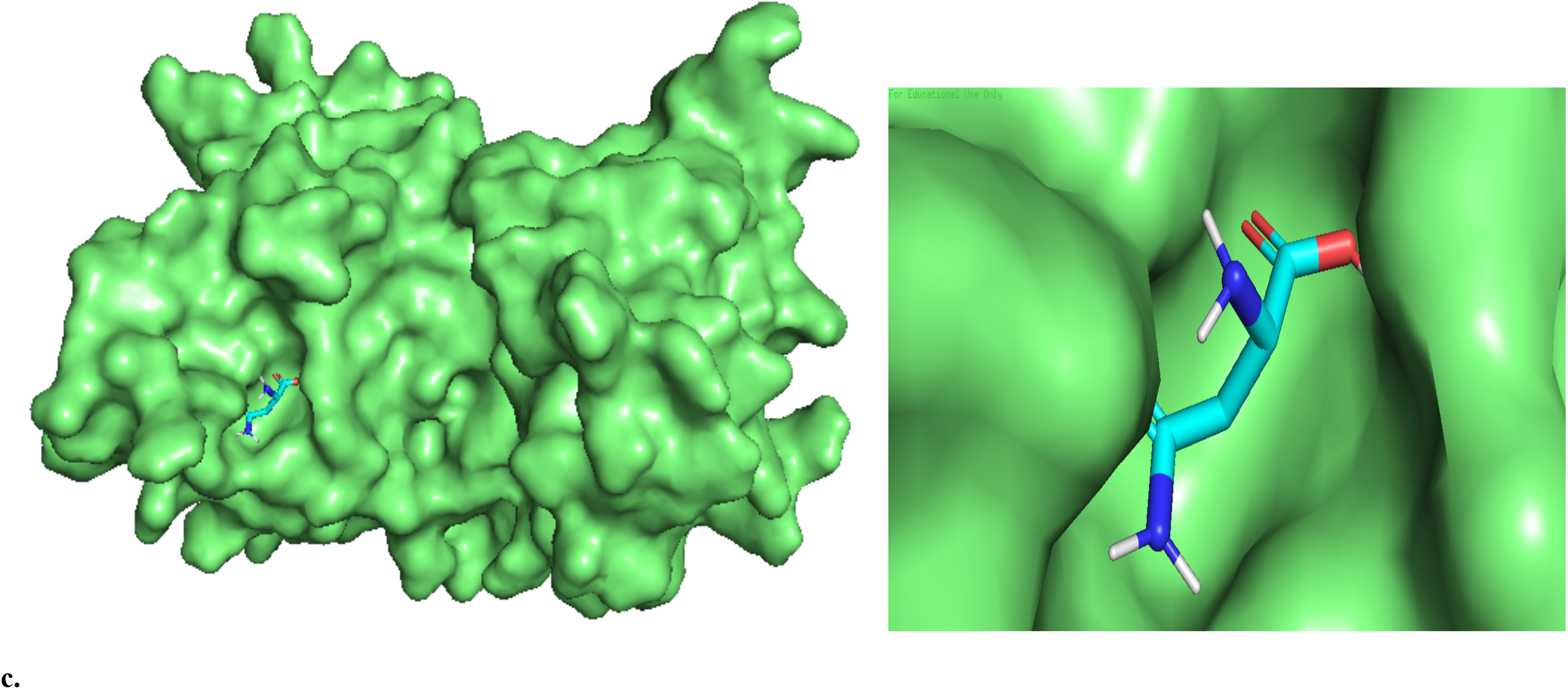
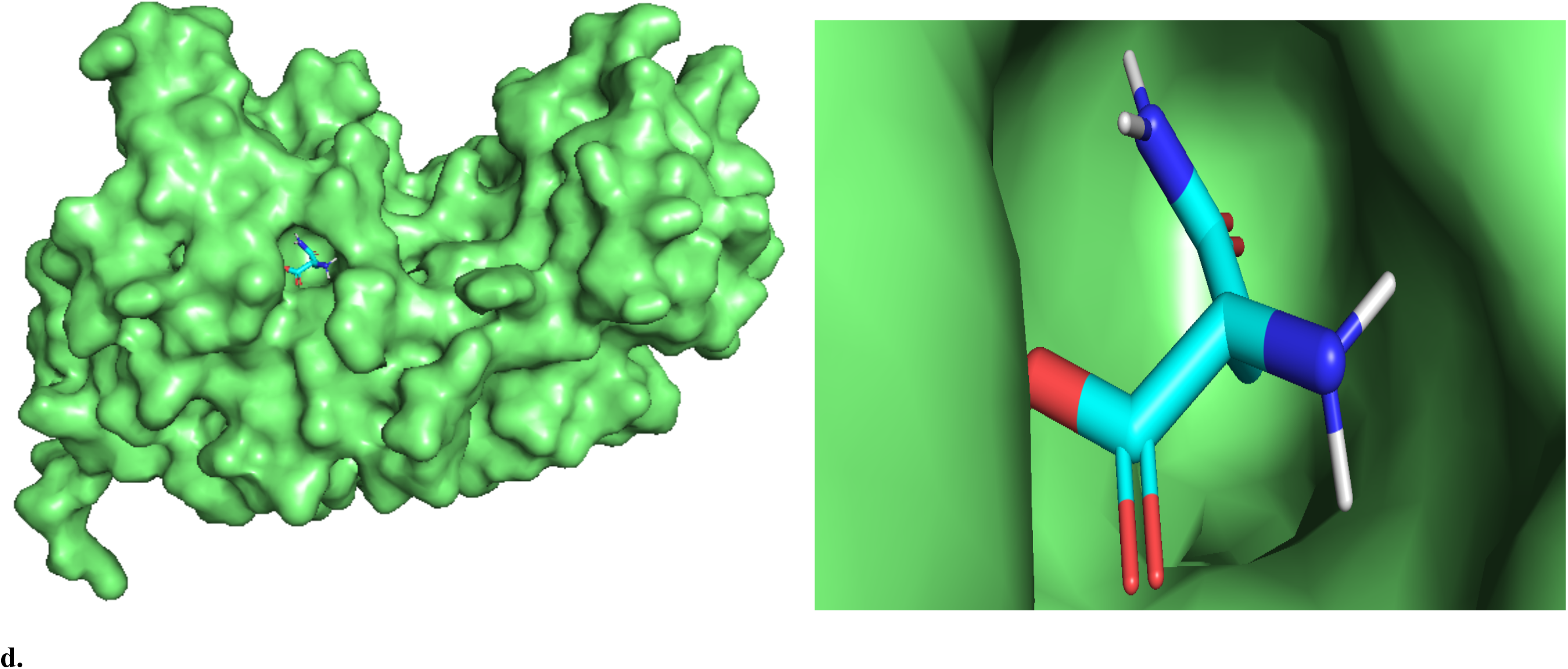
Docked structure of asnB and L-asparagine. (a) *Escherichia coli* asnB and L-asparagine (b) *Streptomyces griseus 1* asnB and L-asparagine (c) *Streptomyces venezuelae* 1 asnB and L-asparagine (d) *Streptomyces collinus* asnB and L-asparagine. Lasparagine is seen to be completely impended in catalytic pocket of the enzymes.

**TABLE 3:** Binding energy of distant organism’s asnB and L-asparagine. *Streptomyces collinus*, *Streptomyces griseus 1*, *Streptomyces venezuelae 2* asnBs have −5.3 kcal/mol, 5.3 kcal/mol, 5.2 kcal/mol binding energy which is greater than the *E.coli* and *Dickeya chrysanthami* −5.1 and −5.0 kcal/mol respectively which indicate that these organisms’ asnB have a greater affinity toward the L-asparagine.

| Organisms | Binding affinity (kcal/mol) calculated from docking |
| --- | --- |
| *Streptomyces albidoflavus* 1 | -4.8 |
| *Streptomyces albidoflavus* 2 | -4.8 |
| *Streptomyces albidoflavus* 3 | -4.5 |
| *Streptomyces aurantiacus* | -4.2 |
| *Streptomyces collinus* | -5.3 |
| *Streptomyces fladiae* 1 | -4.9 |
| *Streptomyces fladiae* 2 | -4.9 |
| *Streptomyces globisporus* | -4.2 |
| *Streptomyces griseus* 1 | -5.3 |
| *Streptomyces griseus 2* | -4.6 |
| *Streptomyces katrae* | -4.9 |
| *Streptomyces venezuelae 1* | -4.8 |
| *Streptomyces venezuelae 2* | -5.2 |

### 3.6. Interaction with active sites

LigPlot showing active site interactions of asnB and asparagine was constructed and shown in Figure 7. The active site of E. coli asnB contains all 5 active site residues. Four of those residues T(34), S(80), Q(81), T(111) form direct hydrogen bonding with asparagine. D(112), unlike in the 1nns active site predicted by pdb, does not form hydrogen bond, and only stays in the active site as a hydrophobic interactor in our LigPlot model(Fig 8a). As 1nns is the structure complexed with aspartic acid (D), a closer inspection of the active site interactions in the 1nns predicted in the pdb website and our LigPlot model show some similarities and some variations.

**Figure 8:**
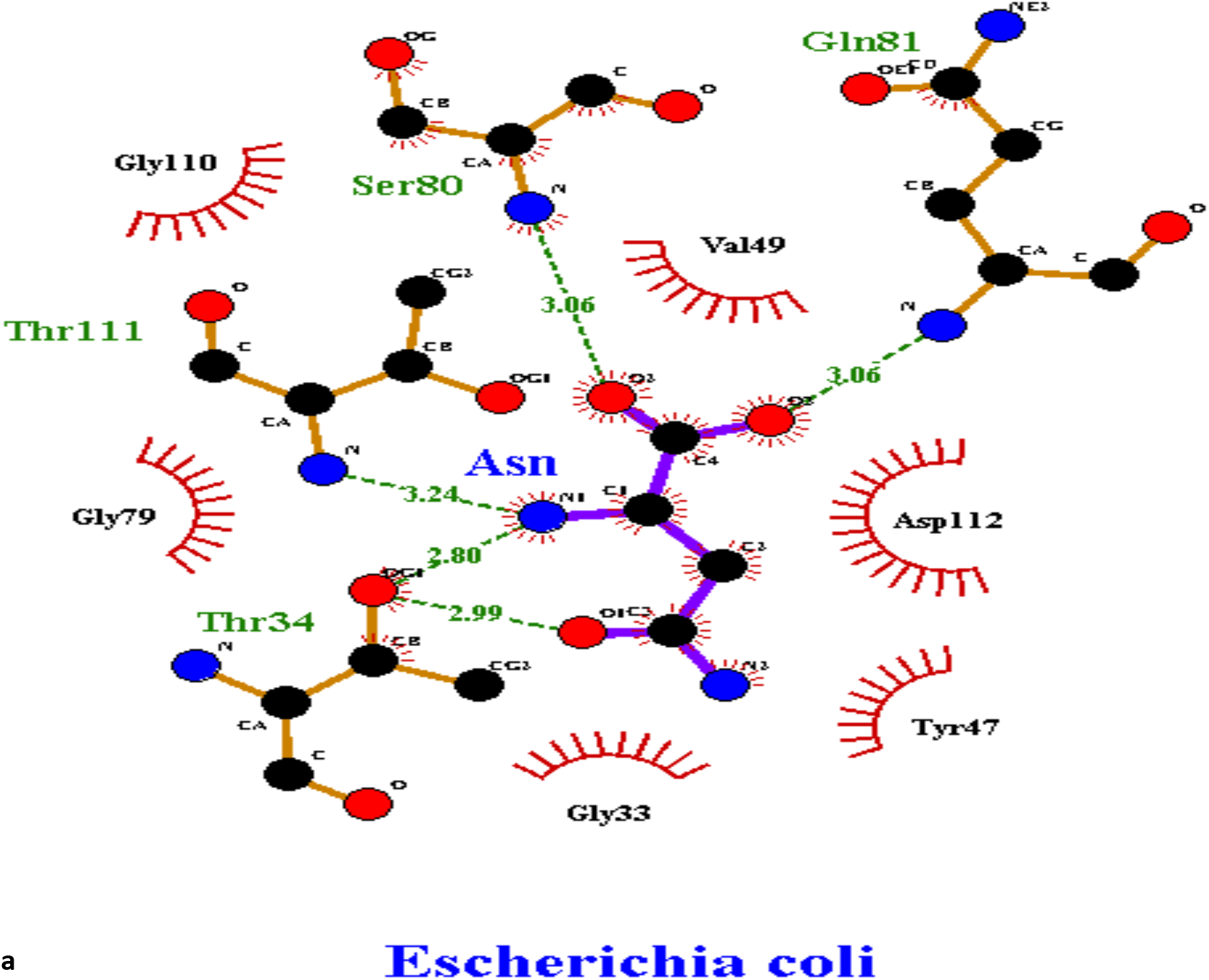
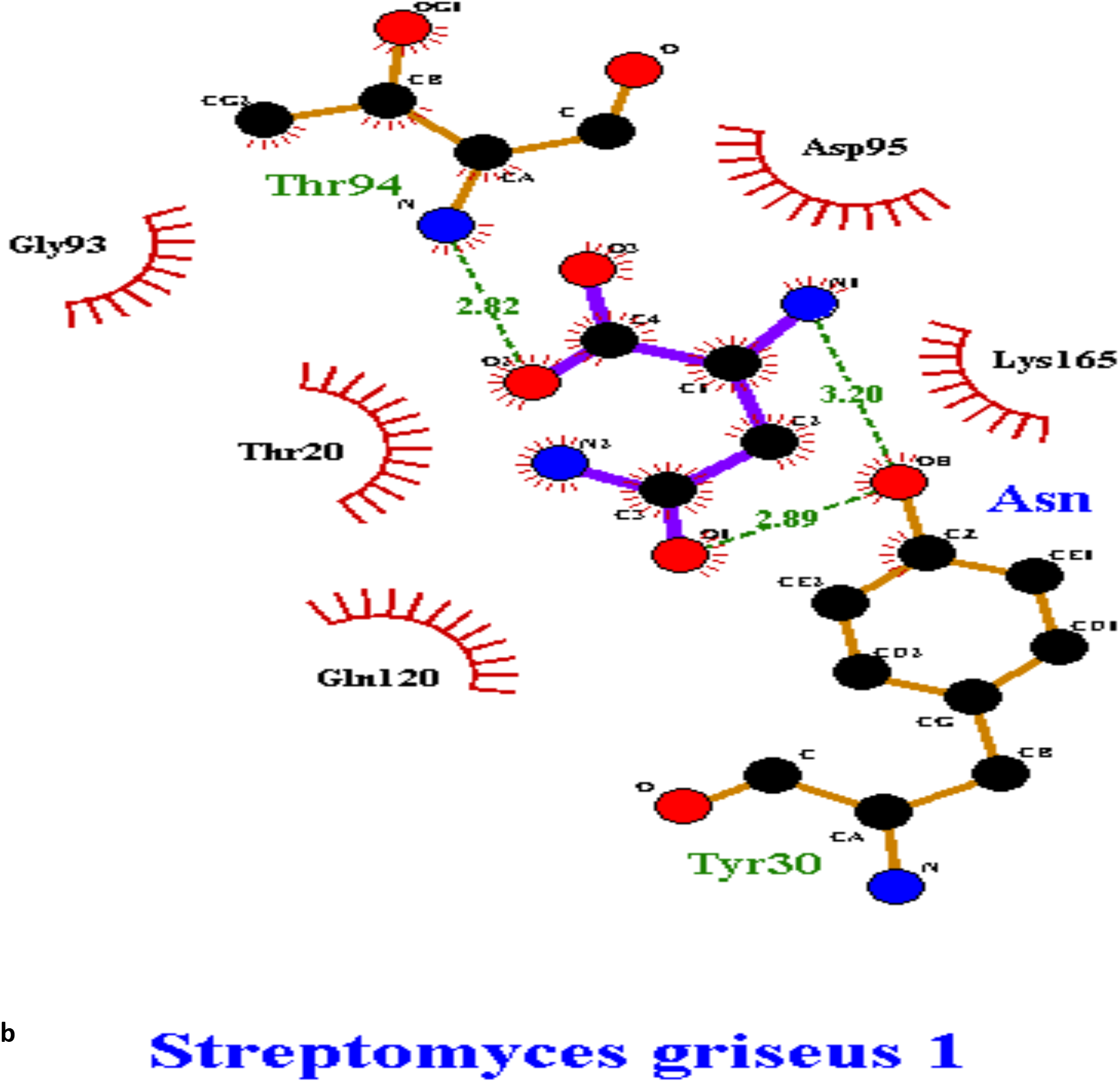
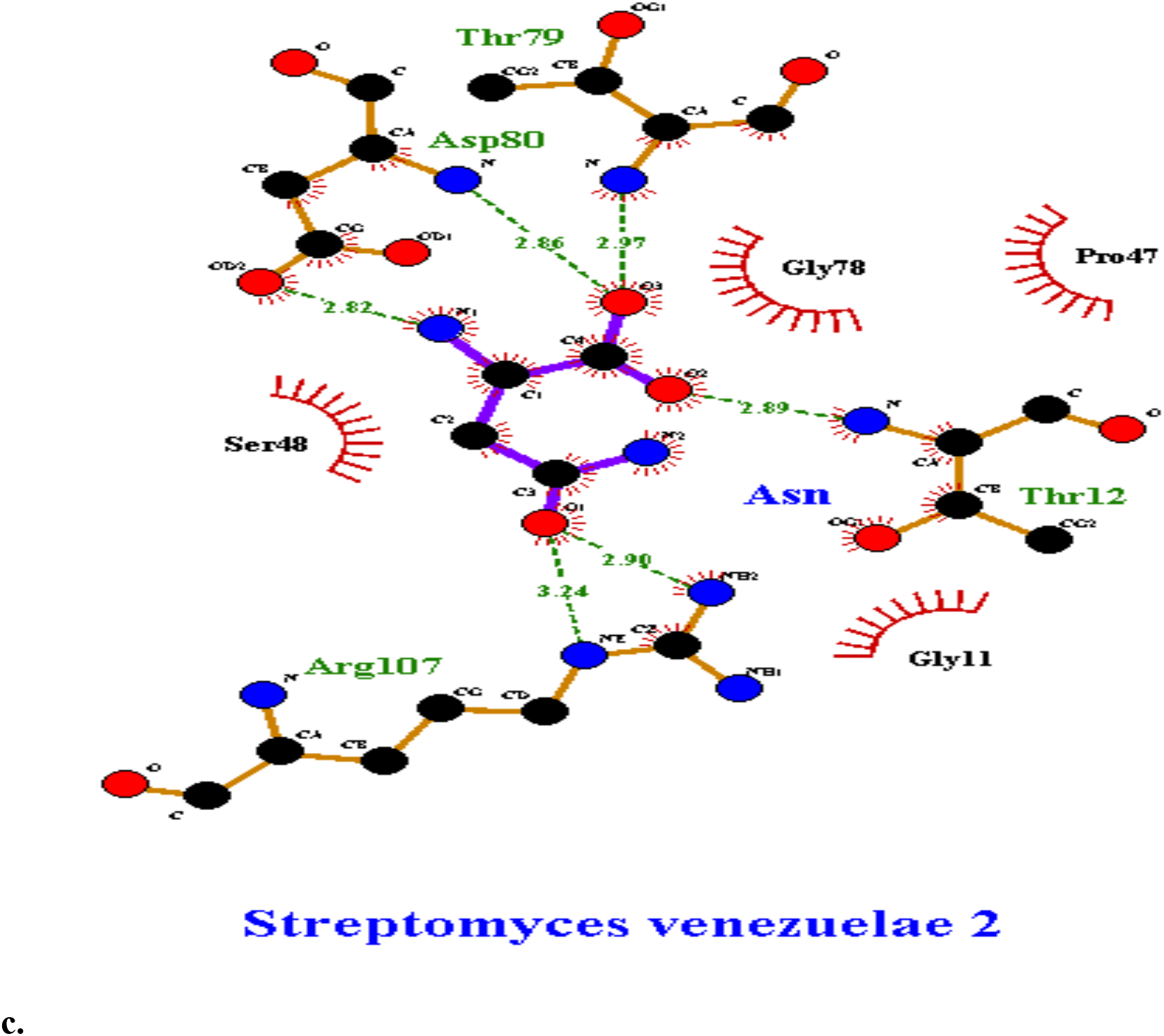
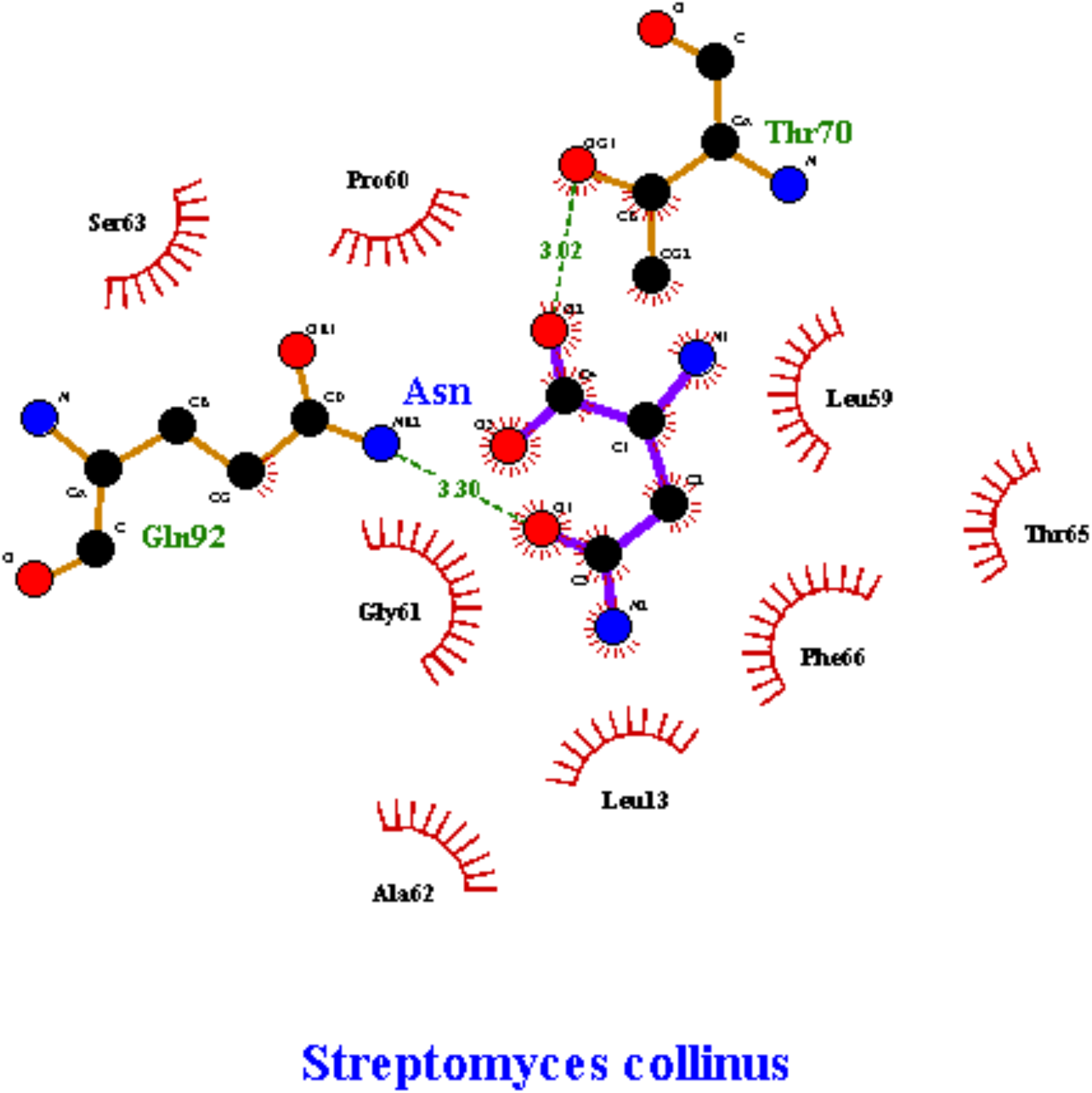
LigPlot of interacting atoms of E.coli and selected three organisms **(a**) *Escherichia coli* **(b**) *Streptomyces griseus 1* **(c)** *Streptomyces venezuelae* 1 **(d)***Streptomyces collinus* enzymes and L-asparagine (Asn).

LigPlot showing active site interactions of asnB and asparagine was constructed and shown in Figure 7. In Streptomyces griseus 1 asnB, 3 amino acid residues—T(20), T(94), and D(95)—of the pentad(out of five predicted residue) interacts with asparagine (Fig 8b). Out of three residues, only one residue T(94) is involved in the formation of hydrogen bond, whereas two other residues form hydrophobic interaction with asparagine. Y(30) forms another hydrogen bond with asparagine. Only 3 of the pentads were detected in *Streptomyces venezuelae* 2. All three amino acids form H-bond with asparagine. Additionally, R(107) forms a hydrogen bond with asparagine(Fig 8c).

As for *Streptomyces collinus* asnB, four of the catalytic pentad residues—T(16), L(64), T(94), and D(95)—are absent at the catalytic site interaction with asparagine. Only S(63) is present in the active site. When the ligand was docked to the *Streptomyces collinus* asnB predicted active site with the grid box size 25 × 25 × 25 Å, auto dock software automatically detected that there was another catalytical pocket present adjacent to the predicted one with almost the same interacting residues(Fig 8d) as predicted but with the different position that gives the binding energy of −5.3 kcal/mol, where T(70), Q(92) contributes on hydrogen bonding and other residues are involved in hydrophobic interaction. This binding site is shown in figure 8d, and is visibly almost the same but in a different position from all predicted active site residues.

The meaning of the items on the plot is as follows:

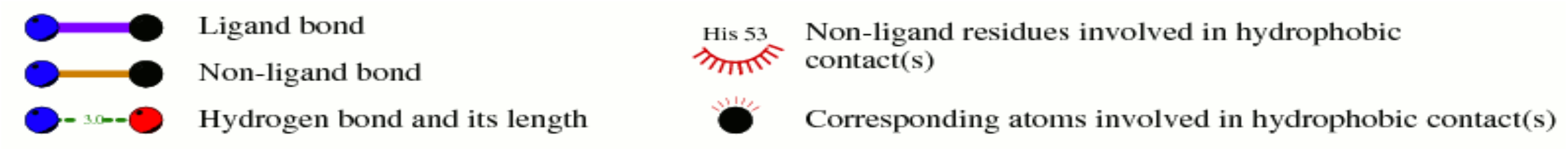

## 6. DISCUSSION

Rapid and cost-effective screening of enzymes is a common undertaking in enzymology. Industrially produced enzymes have a role in a wide range of functions in pharmaceutical, food, biofuel, and chemical industries. Such enzymes are often screened from novel organisms in the soil, water, or other resources. Many of the commercially useful enzymes have been discovered through such screens. The fungus that produces cellulase, Trichoderma reesei, was isolated from garments and canvas that was degraded in the Solomon Islands during the Second World War(Seidl et al., 2009). Similarly, most of the alpha amylases used in the industry find their source in Bacillus(de Souza and e Magalhães, 2010). Asparaginase that is used as an anticancer agent is derived from E. coli and Erwinia. Most of these microorganisms have been discovered from simple screens developed for certain enzymes. This does not necessarily mean that these enzymes have the most optimal sequences for activity. This is because the screen could have easily missed out on better sequences, which are not as well expressed in native cells. If these better sequences could be discovered, they would be easily cloned into amenable expression systems and expressed in high numbers and used for industrial purposes.

In this paper, we have developed a method to in silico screen for the sequence with the best enzymatic activity. Since asnB is one of the most widely screened and studied enzyme, we chose to in silico predict the optimal sequence for its production. The first task was to collect a list of sequences from which optimal sequences could be predicted. This task has been made easier in recent years by an explosion in the number of genomes of organisms sequenced. It has become easy to discover homologous proteins in different phyla and in different domains of life. We collected a total of 101 sequence homologs of asnB from different phyla in bacteria, archaea, and eukarya. Using these 101 sequences, an ML phylogenetic tree was constructed. The tree served two purposes. Firstly, it helped us predict the evolution and history of asnB protein. Since proteins from the same phylum tend to congregate very little in the tree, it can be predicted that there was a lot of horizontal gene transfer during the evolution of asnB. Less than half the species we searched had asnB sequences indicating the lack of universal presence of the enzyme in different organisms. Secondly, the tree helped us pick sequences that we most distant and hence least likely to cause immunogenicity when both E. coli and Erwinia asnBs show immunogenicity. E. coli, being one of the most studied model organisms, was the obvious first choice as a source of asnB. There is no clear indication in the literature as to why Erwinia was chosen as the second source of asnB. But the tree we have drawn confirms that Erwinia as a source was a wise choice since Erwinia asnB lies at one end of the tree distant to E.coli asnB that lies around the center of the tree. The organisms we have zeroed in on are distant compared to Erwinia and E. coli and mostly lie in the Streptomyces genus.

Additionally, we wanted to develop an in silico tool to predict the reaction kinetics of individual enzymes. To that end, we relied on molecular modeling and docking approaches. While reaction kinetics is defined by different parameters like Km, kcat, maximum velocity (Vmax), specificity constant (k_cat_/Km), Km is often the most widely measured quantity. This turned out to be the case for asnBs as well. From the literature, 10 Km values corresponding to asnBs from different species were discovered, while only four kcat values were discovered. We set out to discover if the sequence of asnB can predict Km value, without having to determine it experimentally. Through homology modelling we predicted the structures of asnBs with known Km. After that, asparagine (the substrate) was docked onto the predicted asnB structures and binding energy was calculated. This binding energy was compared to measured Km values to detect correlation. Out of 10 species for which Km is known, only in 6 species could Km be definitely assigned to a certain sequence. A clear inverse relationship between Km value and binding energy emerged. A higher Km value corresponded to lower binding energy.

This finding makes sense according to a definition of Km. The Michaelis Menten kinetics is derived using the following equation:

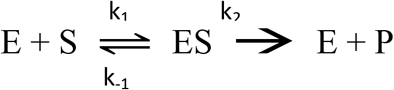

Where E is the enzyme, S is the substrate, ES is the enzyme-substrate complex, P is the product, k_1_ is the rate of forward reaction during the formation of ES complex, k_−1_ is the rate of backward reaction during ES dissociation into E and S and k_2_ is the rate of reaction for the dissociation of ES complex into E and P. From this equation Km is defined as (k_2_ + k_−1_)/ k_1_. When k_2_ ≪ k_−1_ under the rapid equilibrium assumption, K_m_ = k_−1_/k_1_. Thus Km is equal to the dissociation constant. There is also a relationship between dissociation constant and binding energy (deltaG (binding energy) is proportional to −lnKm). But when lnKm is plotted against binding energy was plotted a linear fit graph was not obtained (data not shown). However, the negative relationship between Km and binding energy makes sense from this equation(Nelson and Cox, 2012).

This result demonstrates that if binding energies can be compared among homologs, the homolog with the highest binding energy will give the lowest Km value. This can be used to predict enzyme sequence, which will give the lowest Km value. In this paper, the binding energies of asnBs from various Streptomyces species were calculated to obtain the one with the highest binding energy. Three of the thirteen asnBs give biding energy of −5.3 kcal/mol and −5.2 kcal/mol with asparagine. asnBs from *Streptomyces griseus*, *Streptomyces collinus,* and *Streptomyces venezuelae* gave these values. This value is higher than the binding energy of E. coli and Erwinia asnBs. Thus it can be predicted that an enzyme with better kinetics that currently commercially available asparaginase can be cloned from Streptomyces species.

For the three optimal asnBs and E. coli asnB, LigPlot diagram with of the active site along with interacting aspargine was drawn. It was demonstrated in E. coli that the catalytic pentad residues were actively involved in bonding. Four of the five active-site residues formed hydrogen bonds, whereas one stayed in the active site forming hydrophobic interaction. Although the residues interacting are the same in the active site published by pdb site, different amino acid residues form hydrogen bonds with asparagine at different locations from the one given in the LigPlot in this paper. This is in line with the idea that the exact mechanism of asparaginase catalysis is not figured out, though it is predicted that the mechanism for type I and type II asparaginases will be conserved(Schalk et al., 2016). Two different mechanisms have been proposed for asparaginase catalysis. One mechanism describes double displacement, where the ammonia in asparagine is first displaced by the enzyme before the enzyme attached to asparagine is again displaced by water. The second mechanism describes the single displacement where water directly displaces ammonia from asparagine. There are contrary experimental and theoretical predictions for the validity of the two models (Schalk et al., 2016),(Lubkowski and Wlodawer, 2019).

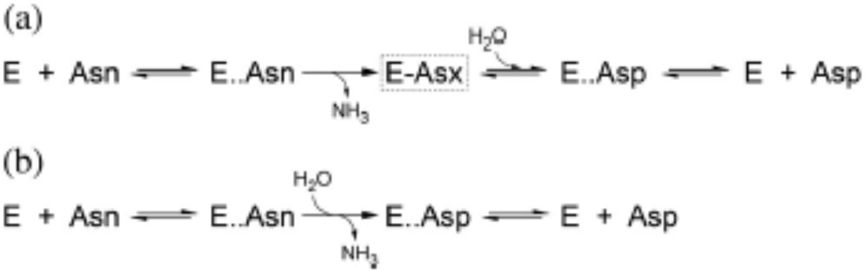

From the LigPlot of *Streptomyces griseus 1* and *Streptomyces venezuelae 2*, it can be demonstrated that three of the pentad residues are present in the active site. This shows that the active site in these distant species is conserved. It has been predicted that one of the two threonines acts as a nucleophile in the double displacement mechanism. Conservation of both threonines suggests that this could indeed be the case. A dynamic simulation modeling rather than the static docking modeling we have carried out might give a clearer answer to the active sites involved, the catalytic mechanism, and the relevant nucleophiles and electrophiles.

## 5. CONCLUSION

Thus, we have devised a method to *in silico* predict the enzyme kinetics (Km value) from a sequence of an enzyme. In this paper, we have validated the effectiveness of this method to predict Km values of asparaginase II with a high degree of accuracy. This method is not only applicable to asparaginases but also to a slew of other industrial proteins such as amylases, cellulases, and many others. In the future, it will be worthwhile to apply this technique to the prediction of Km and the selection of industrially valuable sequences of other enzymes. We have predicted three possible highly promising L-Asparaginase II enzymes produced by three Streptomyces species. The next step will be to verify using cloning if these sequences give a very low Km value.

## 6. Acknowledgements

The authors acknowledge the Department of Biotechnology, Kathmandu University, Dhulikhel, Nepal for providing all support during the study period. The authors would also like to thanks Mr. Sandesh Acharya and Mr. Prithivi Jung Thapa for their guidance on the docking procedure.

## 7. Author Contributions

H.K.B. conceived and initiated the project. H.K.B. and Ad.B. designed the experiments. Ad.B. and R.G. carried out the phylogenetic analysis. Ad.B. and Am.B. carried out homology modelling. Ad.B. and S.K. worked on the verification of the protein model. Ad.B. and R.G. carried out the docking experiment. H.K.B., Ad.B., and R.G. wrote the manuscript. All authors read and approved the final manuscript.

