## Supplemental file for "In-silico development of a method for the selection of optimal enzymes using L-asparaginase II against Acute Lymphoblastic Leukemia as an example"

**Author information.**

**Affiliation**

**Department of Biotechnology, Kathmandu University, Dhulikhel, Nepal.**

Adesh Baral, Ritesh Gorkhali , Amit Basnet, Shubham Koirala, Hitesh K. Bhattarai

**Corresponding author**.

**Validation of other structures used in the experiment.**

*Streptomyces albidoflavus*

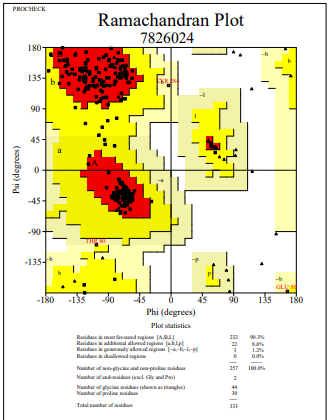

**Figure 1**: Ramachandran Plot of model protein asnB of *Streptomyces albidoflavus.* The Ramachandran plot shows the phi-psi torsion angles for all residues (black cubic box) in the structure (except those at the chain termini). Glycine residues are separately identified by triangles as these are not restricted to the regions of the plot appropriate to the other sidechain types. The darkest red area indicates the "core" regions representing the most favourable combinations of phi-psi values. The regions are labelled as follows: **A** -Core alpha , **L**	- Core left-handed alpha , **a**	- Allowed alpha , **l**	- Allowed left-handed alpha , **~a** - Generous alpha , **~l**	- Generous left-handed alpha, **B**	- Core beta ,**p**	- Allowed epsilon , **b** - Allowed beta ,**~p** - Generous epsilon , **~b** - Generous beta.

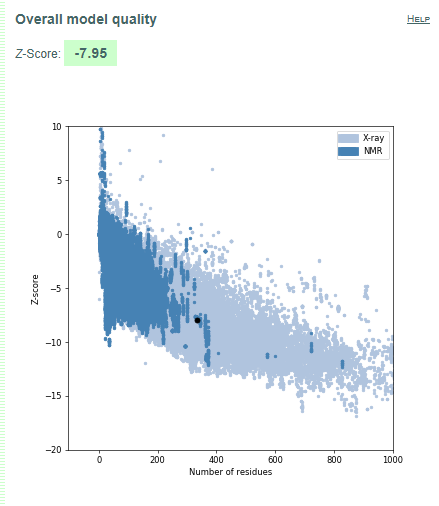

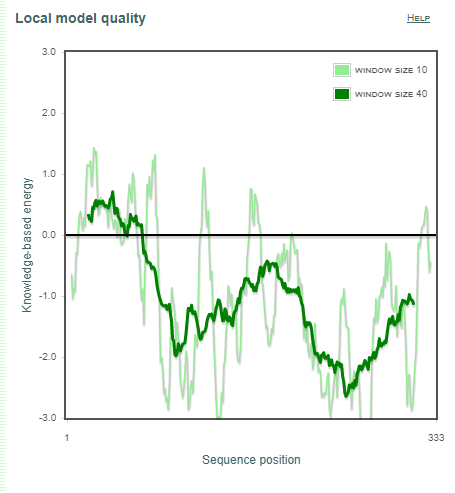

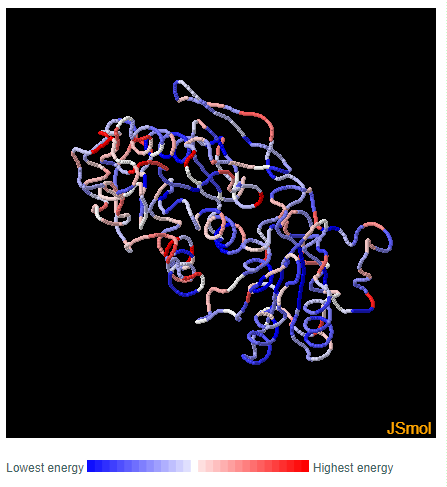

**a	b	c**

**Figure 2.** Validation of model protein structure of asnB of *Streptomyces albidoflavus* **(a)** ProSA-web z-scores of all protein chains in PDB determined by X-ray crystallography (light blue) or NMR spectroscopy (dark blue) with respect to their length. Black dot in the plot indicates that the model protein structure falls inside the range of plot that contains the z-score of all the experimentally determined protein in the PDB. The plot shows only chains with less than 1000 residues and a z-score 10. The z-scores of model proteins are highlighted as large dots. **(b)** Energy plot of model protein which indicates the local model quality by plotting energy as the function of amino acid sequence. Generally, the portion in the positive region of the plot indicates the erroneous part of the structure **(c)** Residues are colored from blue to red in the order of increasing residue energy.

**RESULT:**
*Streptomyces albidoflavus* (Fig.1) plot shows that 90.3% of residues in most favoured regions, 8.6% in additional allowed regions, 1.2% residues in generously allowed regions and 0.0% residues in disallowed regions. More than 99% residues are in allowed region given by Ramachandran plot indicates a very good model. Furthermore, the Ramachandran Z-score calculated by WHATCHECK -0.799 falls on the accepted region(Sousa et al., 2006) and allowed by the WHATCHECK. The structures were finally validated using ProSA-web server. This server gives the z-score which indicates the overall model quality and measures the deviation of the total the total energy of the structure with respect to an energy distribution derived from random conformations(Hooft et al., 1997).The z-score given by the server -7.95 falls inside the range of plot (black dot) that contains the z-score of all the experimentally determined protein in the PDB (X-ray, NMR) (Fig. 2a). In the energy plot (Figs. 2b) which indicates the local model quality by plotting energy as the function of amino acid sequence. Generally, the portion in the positive region of the plot indicates the erroneous part of the structure. We can conclude from the plot that the structure in feasible or accepted as overall residues energies fall under the negative part of the plot. The colored 3D structure of the proteins (Figs. 2c,) shows that the portion in red color is of high energy and portion with the blue color are of low energy(Wiederstein & Sippl, 2007).

*Streptomyces albidoflavus 2*

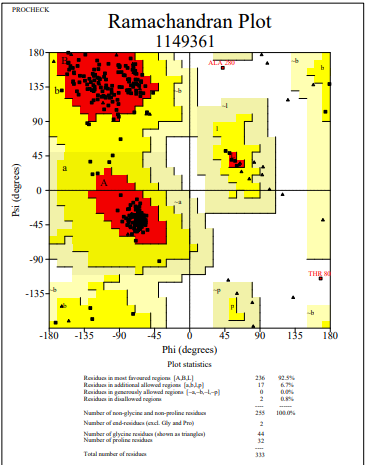

**Figure 3**: Ramachandran Plot of model protein asnB of *Streptomyces albidoflavus 2.* The Ramachandran plot shows the phi-psi torsion angles for all residues (black cubic box) in the structure (except those at the chain termini). Glycine residues are separately identified by triangles as these are not restricted to the regions of the plot appropriate to the other sidechain types. The darkest red area indicates the "core" regions representing the most favourable combinations of phi-psi values. The regions are labelled as follows: **A** -Core alpha , **L**	- Core left-handed alpha , **a**	- Allowed alpha , **l**	- Allowed left-handed alpha , **~a** - Generous alpha , **~l**	- Generous left-handed alpha, **B**	- Core beta ,**p**	- Allowed epsilon , **b** - Allowed beta ,**~p** - Generous epsilon , **~b** - Generous beta.

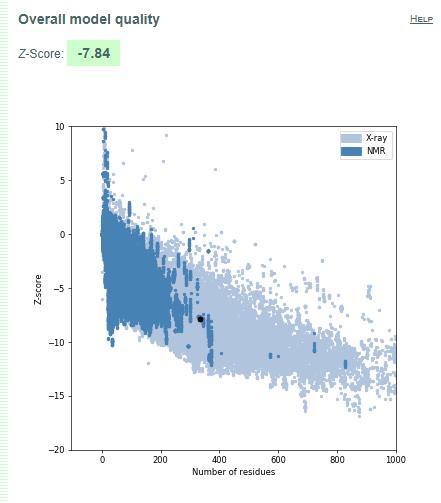

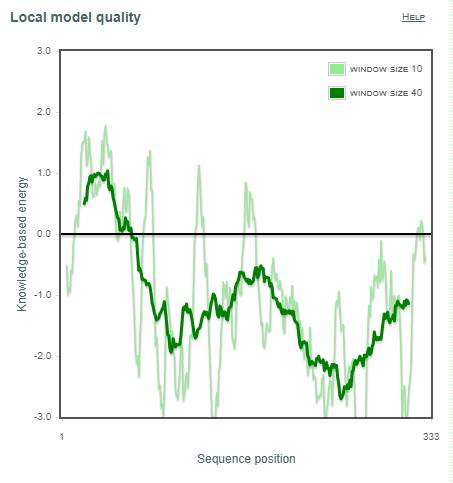

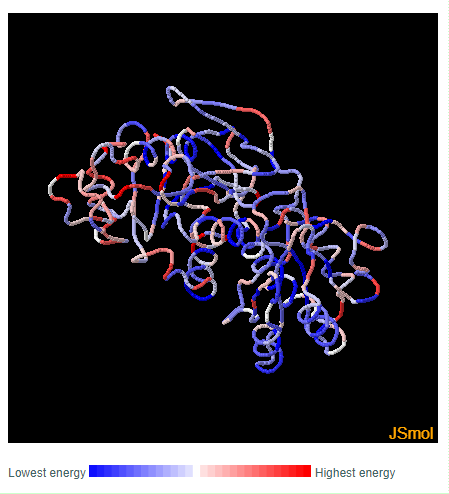

**a	b	c**

**Figure 4.** Validation of model protein structure of asnB of *Streptomyces albidoflavus* 2 **(a)** ProSA-web z-scores of all protein chains in PDB determined by X-ray crystallography (light blue) or NMR spectroscopy (dark blue) with respect to their length. Black dot in the plot indicates that the model protein structure falls inside the range of plot that contains the z-score of all the experimentally determined protein in the PDB. The plot shows only chains with less than 1000 residues and a z-score 10. The z-scores of model proteins are highlighted as large dots. **(b)** Energy plot of model protein which indicates the local model quality by plotting energy as the function of amino acid sequence. Generally, the portion in the positive region of the plot indicates the erroneous part of the structure **(c)** Residues are colored from blue to red in the order of increasing residue energy.

**RESULT:**
*Streptomyces albidoflavus 2* (Fig.3) plot shows that 92.5% of residues in most favoured regions, 6.7% in additional allowed regions, 0.0% residues in generously allowed regions and 0.8% residues in disallowed regions. More than 99% residues are in allowed region given by Ramachandran plot indicates a very good model. Furthermore, the Ramachandran Z-score calculated by WHATCHECK -0.690 falls on the accepted region(Sousa et al., 2006) and allowed by the WHATCHECK. The structures were finally validated using ProSA-web server. This server gives the z-score which indicates the overall model quality and measures the deviation of the total the total energy of the structure with respect to an energy distribution derived from random conformations(Hooft et al., 1997).The z-score given by the server -7.84 falls inside the range of plot (black dot) that contains the z-score of all the experimentally determined protein in the PDB (X-ray, NMR) (Fig. 4a). In the energy plot (Figs. 4b) which indicates the local model quality by plotting energy as the function of amino acid sequence. Generally, the portion in the positive region of the plot indicates the erroneous part of the structure. We can conclude from the plot that the structure in feasible or accepted as overall residues energies fall under the negative part of the plot. The colored 3D structure of the proteins (Figs. 4c,) shows that the portion in red color is of high energy and portion with the blue color are of low energy(Wiederstein & Sippl, 2007).

*Streptomyces albidoflavus 3*

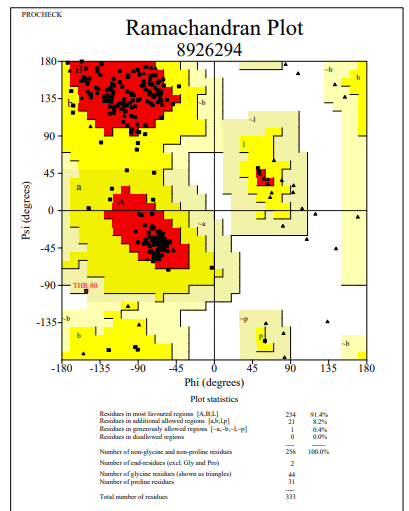

**Figure 5**: Ramachandran Plot of model protein asnB of *Streptomyces albidoflavus 3.* The Ramachandran plot shows the phi-psi torsion angles for all residues (black cubic box) in the structure (except those at the chain termini). Glycine residues are separately identified by triangles as these are not restricted to the regions of the plot appropriate to the other sidechain types. The darkest red area indicates the "core" regions representing the most favourable combinations of phi-psi values. The regions are labelled as follows: **A** -Core alpha , **L**	- Core left-handed alpha , **a**	- Allowed alpha , **l**	- Allowed left-handed alpha , **~a** - Generous alpha , **~l**	- Generous left-handed alpha, **B**	- Core beta ,**p**	- Allowed epsilon , **b** - Allowed beta ,**~p** - Generous epsilon , **~b** - Generous beta.

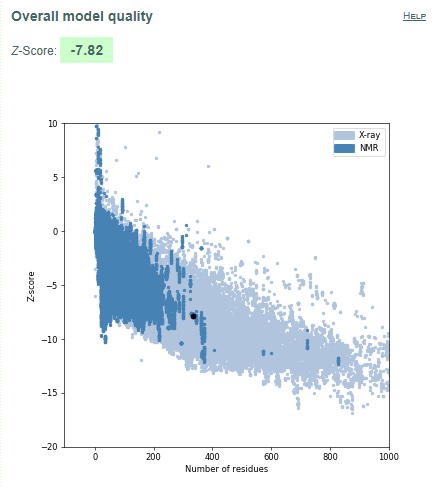

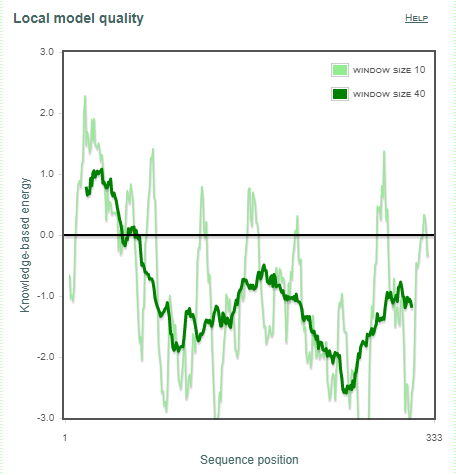

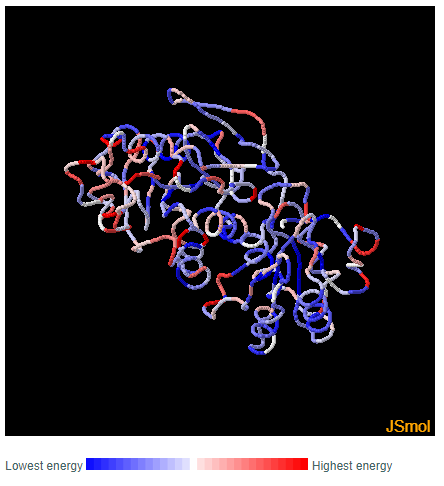

**a	b	c**

**Figure 6.** Validation of model protein structure of asnB of *Streptomyces albidoflavus* *3* **(a)** ProSA-web z-scores of all protein chains in PDB determined by X-ray crystallography (light blue) or NMR spectroscopy (dark blue) with respect to their length. Black dot in the plot indicates that the model protein structure falls inside the range of plot that contains the z-score of all the experimentally determined protein in the PDB. The plot shows only chains with less than 1000 residues and a z-score 10. The z-scores of model proteins are highlighted as large dots. **(b)** Energy plot of model protein which indicates the local model quality by plotting energy as the function of amino acid sequence. Generally, the portion in the positive region of the plot indicates the erroneous part of the structure **(c)** Residues are colored from blue to red in the order of increasing residue energy.

**RESULT:**
*Streptomyces albidoflavus 3* (Fig.5) plot shows that 91.4% of residues in most favoured regions, 8.2% in additional allowed regions, 0.4% residues in generously allowed regions and 0.0% residues in disallowed regions. More than 99% residues are in allowed region given by Ramachandran plot indicates a very good model. Furthermore, the Ramachandran Z-score calculated by WHATCHECK -0.938 falls on the accepted region(Sousa et al., 2006) and allowed by the WHATCHECK. The structures were finally validated using ProSA-web server. This server gives the z-score which indicates the overall model quality and measures the deviation of the total the total energy of the structure with respect to an energy distribution derived from random conformations[16].The z-score given by the server -7.82 falls inside the range of plot (black dot) that contains the z-score of all the experimentally determined protein in the PDB (X-ray, NMR) (Fig. 6a). In the energy plot (Figs. 6b) which indicates the local model quality by plotting energy as the function of amino acid sequence. Generally, the portion in the positive region of the plot indicates the erroneous part of the structure. We can conclude from the plot that the structure in feasible or accepted as overall residues energies fall under the negative part of the plot. The colored 3D structure of the proteins (Figs. 6c,) shows that the portion in red color is of high energy and portion with the blue color are of low energy[48].

*Streptomyces aurantiacus*

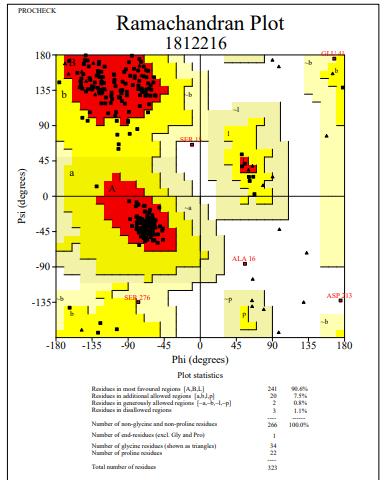

**Figure 7**: Ramachandran Plot of model protein asnB of *Streptomyces aurantiacus.* The Ramachandran plot shows the phi-psi torsion angles for all residues (black cubic box) in the structure (except those at the chain termini). Glycine residues are separately identified by triangles as these are not restricted to the regions of the plot appropriate to the other sidechain types. The darkest red area indicates the "core" regions representing the most favourable combinations of phi-psi values. The regions are labelled as follows: **A** -Core alpha , **L**	- Core left-handed alpha , **a**	- Allowed alpha , **l**	- Allowed left-handed alpha , **~a** - Generous alpha , **~l**	- Generous left-handed alpha, **B**	- Core beta ,**p**	- Allowed epsilon , **b** - Allowed beta ,**~p** - Generous epsilon , **~b** - Generous beta.

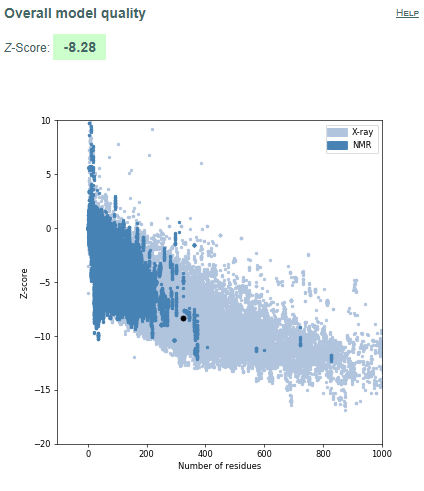

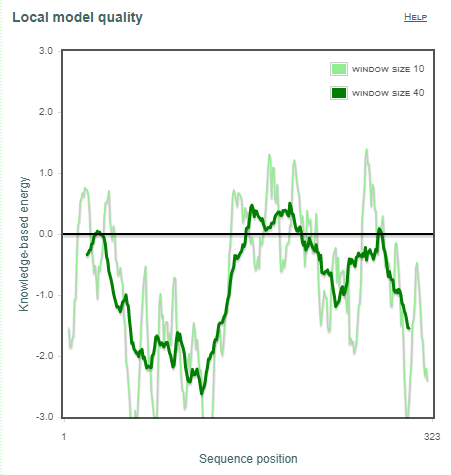

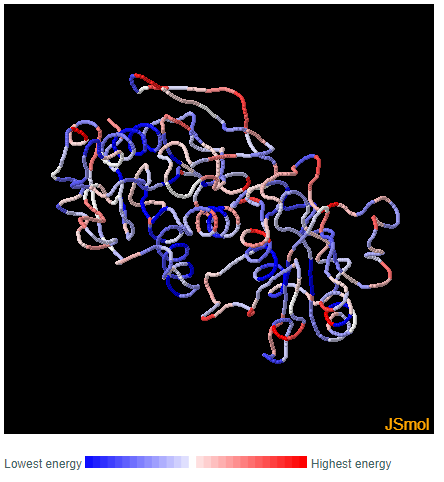

**a	b	c**

**Figure 8.** Validation of model protein structure of asnB of *Streptomyces aurantiacus* **(a)** ProSA-web z-scores of all protein chains in PDB determined by X-ray crystallography (light blue) or NMR spectroscopy (dark blue) with respect to their length. Black dot in the plot indicates that the model protein structure falls inside the range of plot that contains the z-score of all the experimentally determined protein in the PDB. The plot shows only chains with less than 1000 residues and a z-score 10. The z-scores of model proteins are highlighted as large dots. **(b)** Energy plot of model protein which indicates the local model quality by plotting energy as the function of amino acid sequence. Generally, the portion in the positive region of the plot indicates the erroneous part of the structure **(c)** Residues are colored from blue to red in the order of increasing residue energy.

**RESULT:**
*Streptomyces aurantiacus* (Fig.7) plot shows that 90.6% of residues in most favoured regions, 7.5% in additional allowed regions, 0.8% residues in generously allowed regions and 1.1% residues in disallowed regions. More than 99% residues are in allowed region given by Ramachandran plot indicates a very good model. Furthermore, the Ramachandran Z-score calculated by WHATCHECK -1.049 falls on the accepted region(Sousa et al., 2006) and allowed by the WHATCHECK. The structures were finally validated using ProSA-web server. This server gives the z-score which indicates the overall model quality and measures the deviation of the total the total energy of the structure with respect to an energy distribution derived from random conformations[16].The z-score given by the server -8.28 falls inside the range of plot (black dot) that contains the z-score of all the experimentally determined protein in the PDB (X-ray, NMR) (Fig. 8a). In the energy plot (Figs. 8b) which indicates the local model quality by plotting energy as the function of amino acid sequence. Generally, the portion in the positive region of the plot indicates the erroneous part of the structure. We can conclude from the plot that the structure in feasible or accepted as overall residues energies fall under the negative part of the plot. The colored 3D structure of the proteins (Figs. 8c,) shows that the portion in red color is of high energy and portion with the blue color are of low energy[48].

*Streptomyces fradiae*

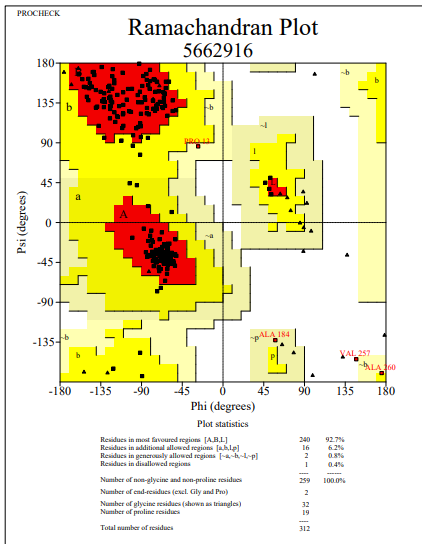

**Figure 9**: Ramachandran Plot of model protein asnB of *Streptomyces fradiae.* The Ramachandran plot shows the phi-psi torsion angles for all residues (black cubic box) in the structure (except those at the chain termini). Glycine residues are separately identified by triangles as these are not restricted to the regions of the plot appropriate to the other sidechain types. The darkest red area indicates the "core" regions representing the most favourable combinations of phi-psi values. The regions are labelled as follows: **A** -Core alpha , **L**	- Core left-handed alpha , **a**	- Allowed alpha , **l**	- Allowed left-handed alpha , **~a** - Generous alpha , **~l**	- Generous left-handed alpha, **B**	- Core beta ,**p**	- Allowed epsilon , **b** - Allowed beta ,**~p** - Generous epsilon , **~b** - Generous beta.

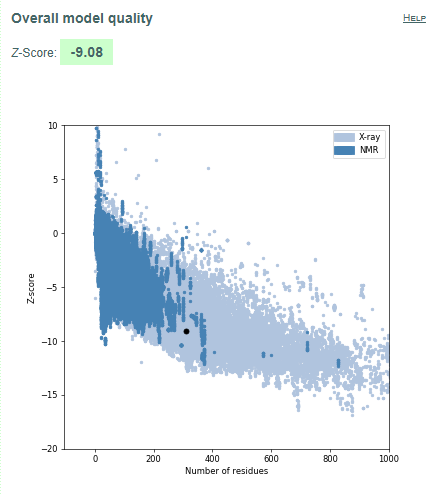

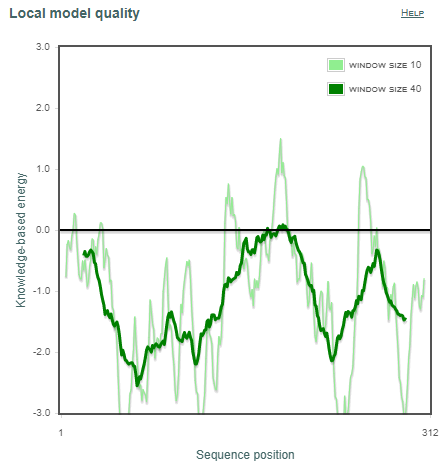

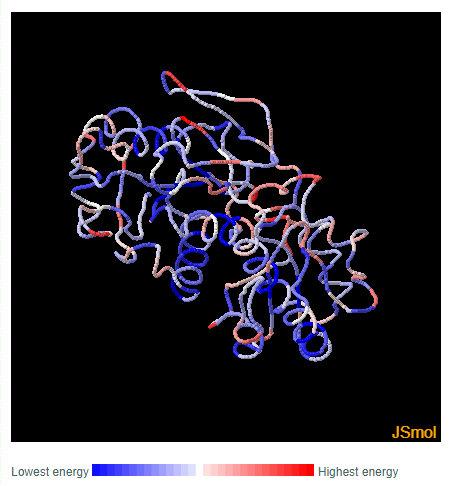

**a	b	c**

**Figure 10.** Validation of model protein structure of asnB of *Streptomyces fradiae* **(a)** ProSA-web z-scores of all protein chains in PDB determined by X-ray crystallography (light blue) or NMR spectroscopy (dark blue) with respect to their length. Black dot in the plot indicates that the model protein structure falls inside the range of plot that contains the z-score of all the experimentally determined protein in the PDB. The plot shows only chains with less than 1000 residues and a z-score 10. The z-scores of model proteins are highlighted as large dots. **(b)** Energy plot of model protein which indicates the local model quality by plotting energy as the function of amino acid sequence. Generally, the portion in the positive region of the plot indicates the erroneous part of the structure **(c)** Residues are colored from blue to red in the order of increasing residue energy.

**RESULT:**
*Streptomyces fradiae* (Fig.9) plot shows that 92.7% of residues in most favoured regions, 6.2% in additional allowed regions, 0.8% residues in generously allowed regions and 0.4% residues in disallowed regions. More than 99% residues are in allowed region given by Ramachandran plot indicates a very good model. Furthermore, the Ramachandran Z-score calculated by WHATCHECK -0.331 falls on the accepted region(Sousa et al., 2006) and allowed by the WHATCHECK. The structures were finally validated using ProSA-web server. This server gives the z-score which indicates the overall model quality and measures the deviation of the total the total energy of the structure with respect to an energy distribution derived from random conformations[16].The z-score given by the server -9.08 falls inside the range of plot (black dot) that contains the z-score of all the experimentally determined protein in the PDB (X-ray, NMR) (Fig. 10a). In the energy plot (Figs. 10b) which indicates the local model quality by plotting energy as the function of amino acid sequence. Generally, the portion in the positive region of the plot indicates the erroneous part of the structure. We can conclude from the plot that the structure in feasible or accepted as overall residues energies fall under the negative part of the plot. The colored 3D structure of the proteins (Figs. 10c,) shows that the portion in red color is of high energy and portion with the blue color are of low energy[48].

*Streptomyces fradiae 2*

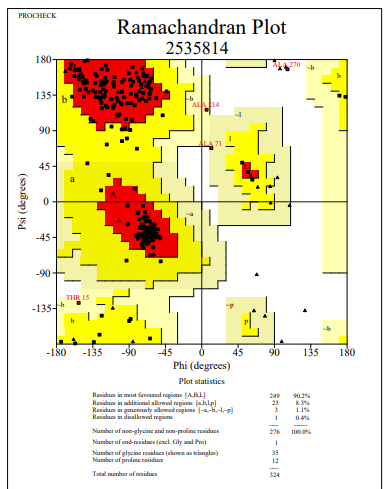

**Figure 11**: Ramachandran Plot of model protein asnB of *Streptomyces fradiae 2.* The Ramachandran plot shows the phi-psi torsion angles for all residues (black cubic box) in the structure (except those at the chain termini). Glycine residues are separately identified by triangles as these are not restricted to the regions of the plot appropriate to the other sidechain types. The darkest red area indicates the "core" regions representing the most favourable combinations of phi-psi values. The regions are labelled as follows: **A** -Core alpha , **L**	- Core left-handed alpha , **a**	- Allowed alpha , **l**	- Allowed left-handed alpha , **~a** - Generous alpha , **~l**	- Generous left-handed alpha, **B**	- Core beta ,**p**	- Allowed epsilon , **b** - Allowed beta ,**~p** - Generous epsilon , **~b** - Generous beta.

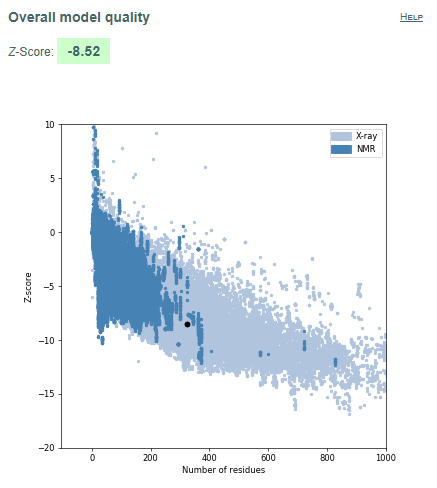

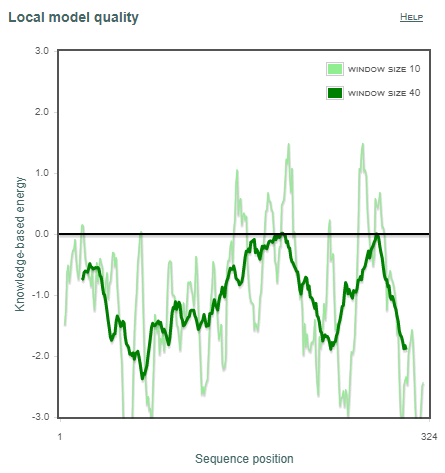

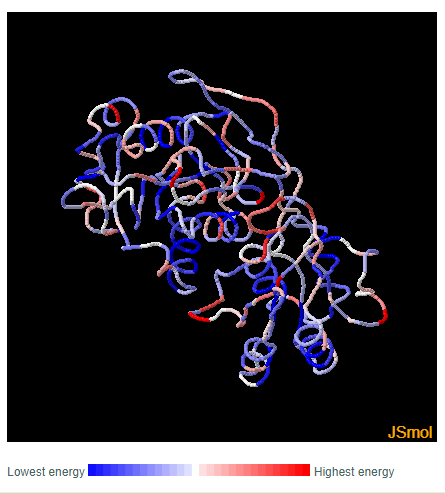

**a	b	c**

**Figure 12.** Validation of model protein structure of asnB of *Streptomyces fradiae 2* **(a)** ProSA-web z-scores of all protein chains in PDB determined by X-ray crystallography (light blue) or NMR spectroscopy (dark blue) with respect to their length. Black dot in the plot indicates that the model protein structure falls inside the range of plot that contains the z-score of all the experimentally determined protein in the PDB. The plot shows only chains with less than 1000 residues and a z-score 10. The z-scores of model proteins are highlighted as large dots. **(b)** Energy plot of model protein which indicates the local model quality by plotting energy as the function of amino acid sequence. Generally, the portion in the positive region of the plot indicates the erroneous part of the structure **(c)** Residues are colored from blue to red in the order of increasing residue energy.

**RESULT:**
*Streptomyces fradiae 2* (Fig.11) plot shows that 90.2% of residues in most favoured regions, 8.3% in additional allowed regions, 1.1% residues in generously allowed regions and 0.4% residues in disallowed regions. More than 99% residues are in allowed region given by Ramachandran plot indicates a very good model. Furthermore, the Ramachandran Z-score calculated by WHATCHECK -0.368 falls on the accepted region(Sousa et al., 2006) and allowed by the WHATCHECK. The structures were finally validated using ProSA-web server. This server gives the z-score which indicates the overall model quality and measures the deviation of the total the total energy of the structure with respect to an energy distribution derived from random conformations[16].The z-score given by the server -8.52 falls inside the range of plot (black dot) that contains the z-score of all the experimentally determined protein in the PDB (X-ray, NMR) (Fig. 12a). In the energy plot (Figs. 12b) which indicates the local model quality by plotting energy as the function of amino acid sequence. Generally, the portion in the positive region of the plot indicates the erroneous part of the structure. We can conclude from the plot that the structure in feasible or accepted as overall residues energies fall under the negative part of the plot. The colored 3D structure of the proteins (Figs. 12c,) shows that the portion in red color is of high energy and portion with the blue color are of low energy[48].

*Streptomyces globisporus*

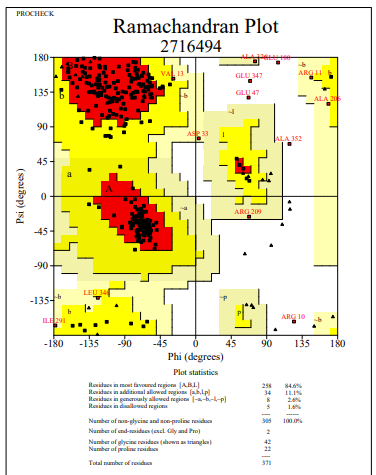

**Figure 13**: Ramachandran Plot of model protein asnB of *Streptomyces globisporus.* The Ramachandran plot shows the phi-psi torsion angles for all residues (black cubic box) in the structure (except those at the chain termini). Glycine residues are separately identified by triangles as these are not restricted to the regions of the plot appropriate to the other sidechain types. The darkest red area indicates the "core" regions representing the most favourable combinations of phi-psi values. The regions are labelled as follows: **A** -Core alpha , **L**	- Core left-handed alpha , **a**	- Allowed alpha , **l**	- Allowed left-handed alpha , **~a** - Generous alpha , **~l**	- Generous left-handed alpha, **B**	- Core beta ,**p**	- Allowed epsilon , **b** - Allowed beta ,**~p** - Generous epsilon , **~b** - Generous beta.

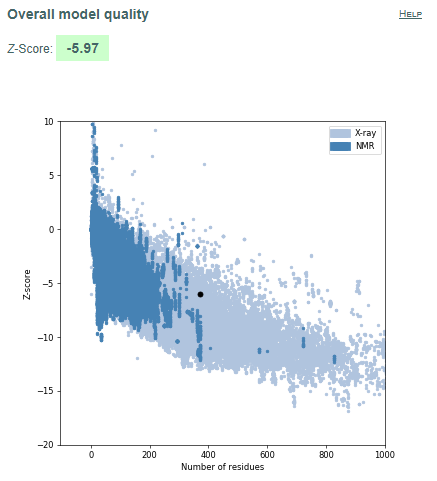

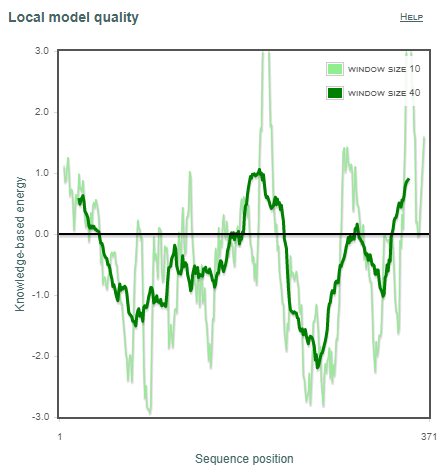

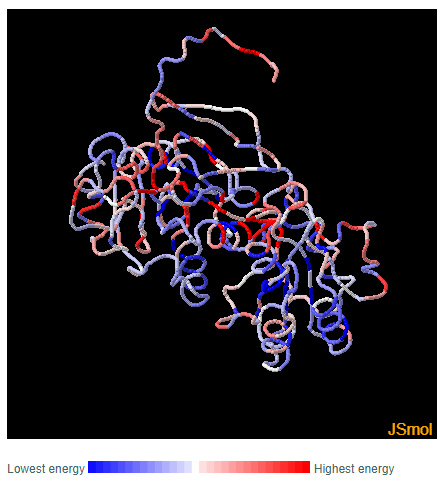

**a	b	c**

**Figure 14.** Validation of model protein structure of asnB of *Streptomyces globisporus* **(a)** ProSA-web z-scores of all protein chains in PDB determined by X-ray crystallography (light blue) or NMR spectroscopy (dark blue) with respect to their length. Black dot in the plot indicates that the model protein structure falls inside the range of plot that contains the z-score of all the experimentally determined protein in the PDB. The plot shows only chains with less than 1000 residues and a z-score 10. The z-scores of model proteins are highlighted as large dots. **(b)** Energy plot of model protein which indicates the local model quality by plotting energy as the function of amino acid sequence. Generally, the portion in the positive region of the plot indicates the erroneous part of the structure **(c)** Residues are colored from blue to red in the order of increasing residue energy.

**RESULT:**
*Streptomyces globisporus* (Fig.13) plot shows that 84.6% of residues in most favoured regions, 11.1% in additional allowed regions, 2.6% residues in generously allowed regions and 1.6% residues in disallowed regions. More than 99% residues are in allowed region given by Ramachandran plot indicates a very good model. Furthermore, the Ramachandran Z-score calculated by WHATCHECK -2.202 falls on the accepted region(Sousa et al., 2006) and allowed by the WHATCHECK. The structures were finally validated using ProSA-web server. This server gives the z-score which indicates the overall model quality and measures the deviation of the total the total energy of the structure with respect to an energy distribution derived from random conformations[16].The z-score given by the server -5.97 falls inside the range of plot (black dot) that contains the z-score of all the experimentally determined protein in the PDB (X-ray, NMR) (Fig. 14a). In the energy plot (Figs. 14b) which indicates the local model quality by plotting energy as the function of amino acid sequence. Generally, the portion in the positive region of the plot indicates the erroneous part of the structure. We can conclude from the plot that the structure in feasible or accepted as overall residues energies fall under the negative part of the plot. The colored 3D structure of the proteins (Figs. 14c,) shows that the portion in red color is of high energy and portion with the blue color are of low energy[48]

*Streptomyces griseus 2*

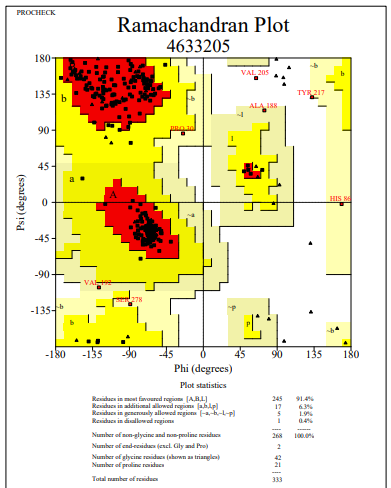

**Figure 15**: Ramachandran Plot of model protein asnB of *Streptomyces griseus 2.* The Ramachandran plot shows the phi-psi torsion angles for all residues (black cubic box) in the structure (except those at the chain termini). Glycine residues are separately identified by triangles as these are not restricted to the regions of the plot appropriate to the other sidechain types. The darkest red area indicates the "core" regions representing the most favourable combinations of phi-psi values. The regions are labelled as follows: **A** -Core alpha , **L**	- Core left-handed alpha , **a**	- Allowed alpha , **l**	- Allowed left-handed alpha , **~a** - Generous alpha , **~l**	- Generous left-handed alpha, **B**	- Core beta ,**p**	- Allowed epsilon , **b** - Allowed beta ,**~p** - Generous epsilon , **~b** - Generous beta.

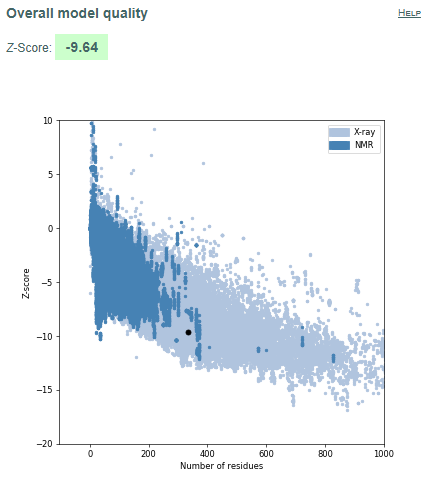

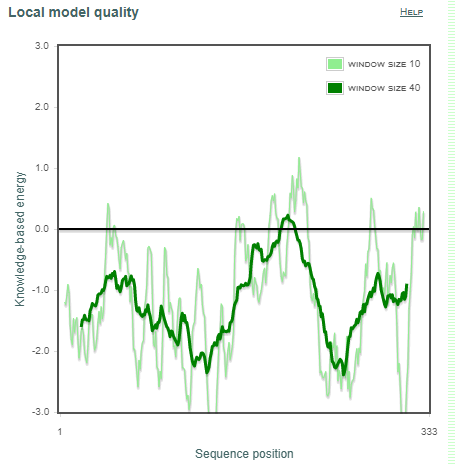

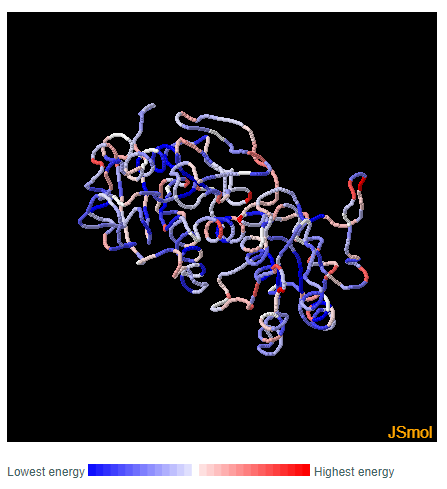

**a	b	c**

**Figure 16.** Validation of model protein structure of asnB of *Streptomyces griseus 2* **(a)** ProSA-web z-scores of all protein chains in PDB determined by X-ray crystallography (light blue) or NMR spectroscopy (dark blue) with respect to their length. Black dot in the plot indicates that the model protein structure falls inside the range of plot that contains the z-score of all the experimentally determined protein in the PDB. The plot shows only chains with less than 1000 residues and a z-score 10. The z-scores of model proteins are highlighted as large dots. **(b)** Energy plot of model protein which indicates the local model quality by plotting energy as the function of amino acid sequence. Generally, the portion in the positive region of the plot indicates the erroneous part of the structure **(c)** Residues are colored from blue to red in the order of increasing residue energy

**RESULT:**
*Streptomyces griseus 2* (Fig.15) plot shows that 91.4% of residues in most favoured regions, 6.3% in additional allowed regions, 1.9% residues in generously allowed regions and 0.4% residues in disallowed regions. More than 99% residues are in allowed region given by Ramachandran plot indicates a very good model. Furthermore, the Ramachandran Z-score calculated by WHATCHECK -0.461 falls on the accepted region(Sousa et al., 2006) and allowed by the WHATCHECK. The structures were finally validated using ProSA-web server. This server gives the z-score which indicates the overall model quality and measures the deviation of the total the total energy of the structure with respect to an energy distribution derived from random conformations[16].The z-score given by the server -9.64 falls inside the range of plot (black dot) that contains the z-score of all the experimentally determined protein in the PDB (X-ray, NMR) (Fig. 16a). In the energy plot (Figs. 16b) which indicates the local model quality by plotting energy as the function of amino acid sequence. Generally, the portion in the positive region of the plot indicates the erroneous part of the structure. We can conclude from the plot that the structure in feasible or accepted as overall residues energies fall under the negative part of the plot. The colored 3D structure of the proteins (Figs. 16c,) shows that the portion in red color is of high energy and portion with the blue color are of low energy[48].

*Streptomyces katrae*

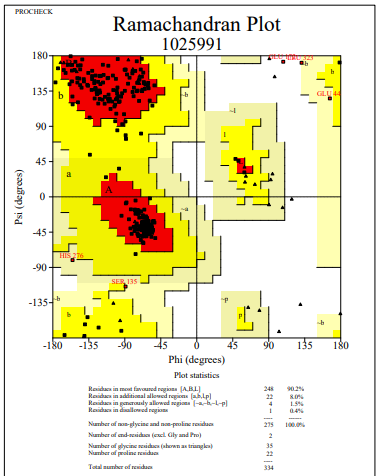

**Figure 17**: Ramachandran Plot of model protein asnB of *Streptomyces katrae.* The Ramachandran plot shows the phi-psi torsion angles for all residues (black cubic box) in the structure (except those at the chain termini). Glycine residues are separately identified by triangles as these are not restricted to the regions of the plot appropriate to the other sidechain types. The darkest red area indicates the "core" regions representing the most favourable combinations of phi-psi values. The regions are labelled as follows: **A** -Core alpha , **L**	- Core left-handed alpha , **a**	- Allowed alpha , **l**	- Allowed left-handed alpha , **~a** - Generous alpha , **~l**	- Generous left-handed alpha, **B**	- Core beta ,**p**	- Allowed epsilon , **b** - Allowed beta ,**~p** - Generous epsilon , **~b** - Generous beta.

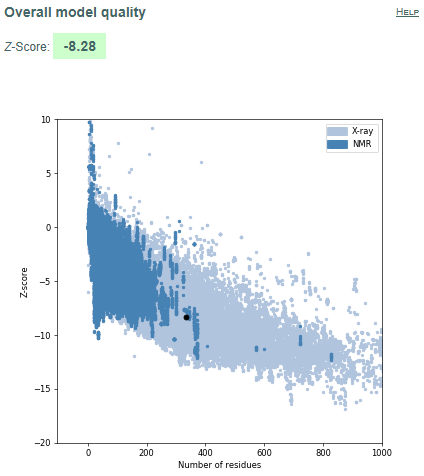

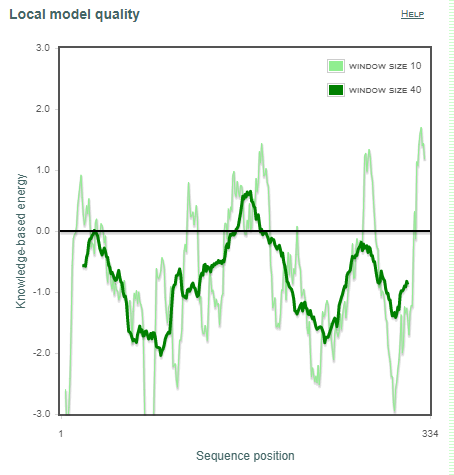

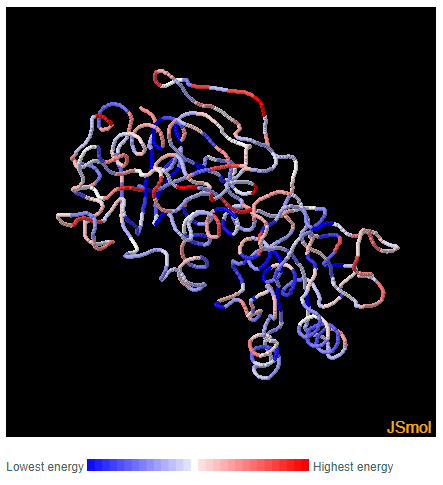

**a	b	c**

**Figure 18.** Validation of model protein structure of asnB of *Streptomyces katrae* **(a)** ProSA-web z-scores of all protein chains in PDB determined by X-ray crystallography (light blue) or NMR spectroscopy (dark blue) with respect to their length. Black dot in the plot indicates that the model protein structure falls inside the range of plot that contains the z-score of all the experimentally determined protein in the PDB. The plot shows only chains with less than 1000 residues and a z-score 10. The z-scores of model proteins are highlighted as large dots. **(b)** Energy plot of model protein which indicates the local model quality by plotting energy as the function of amino acid sequence. Generally, the portion in the positive region of the plot indicates the erroneous part of the structure **(c)** Residues are colored from blue to red in the order of increasing residue energy.

**RESULT:**
*Streptomyces katrae* (Fig.17) plot shows that 90.2% of residues in most favoured regions, 8.0% in additional allowed regions, 1.5% residues in generously allowed regions and 0.4% residues in disallowed regions. More than 99% residues are in allowed region given by Ramachandran plot indicates a very good model. Furthermore, the Ramachandran Z-score calculated by WHATCHECK -0.972 falls on the accepted region(Sousa et al., 2006) and allowed by the WHATCHECK. The structures were finally validated using ProSA-web server. This server gives the z-score which indicates the overall model quality and measures the deviation of the total the total energy of the structure with respect to an energy distribution derived from random conformations[16].The z-score given by the server -8.28 falls inside the range of plot (black dot) that contains the z-score of all the experimentally determined protein in the PDB (X-ray, NMR) (Fig. 18a). In the energy plot (Figs. 18b) which indicates the local model quality by plotting energy as the function of amino acid sequence. Generally, the portion in the positive region of the plot indicates the erroneous part of the structure. We can conclude from the plot that the structure in feasible or accepted as overall residues energies fall under the negative part of the plot. The colored 3D structure of the proteins (Figs. 18c,) shows that the portion in red color is of high energy and portion with the blue color are of low energy[48].

*Streptomyces venezuelae* 1

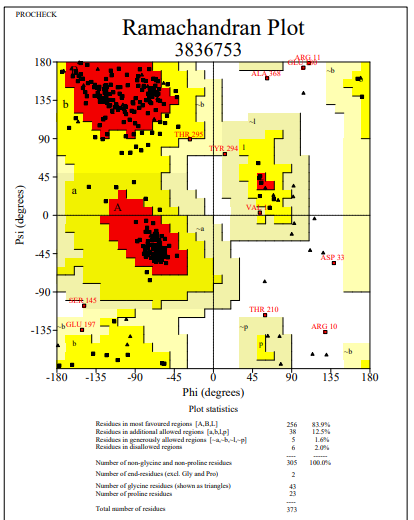

**Figure 19**: Ramachandran Plot of model protein asnB of *Streptomyces venezuelae1.* The Ramachandran plot shows the phi-psi torsion angles for all residues (black cubic box) in the structure (except those at the chain termini). Glycine residues are separately identified by triangles as these are not restricted to the regions of the plot appropriate to the other sidechain types. The darkest red area indicates the "core" regions representing the most favourable combinations of phi-psi values. The regions are labelled as follows: **A** -Core alpha , **L**	- Core left-handed alpha , **a**	- Allowed alpha , **l**	- Allowed left-handed alpha , **~a** - Generous alpha , **~l**	- Generous left-handed alpha, **B**	- Core beta ,**p**	- Allowed epsilon , **b** - Allowed beta ,**~p** - Generous epsilon , **~b** - Generous beta.

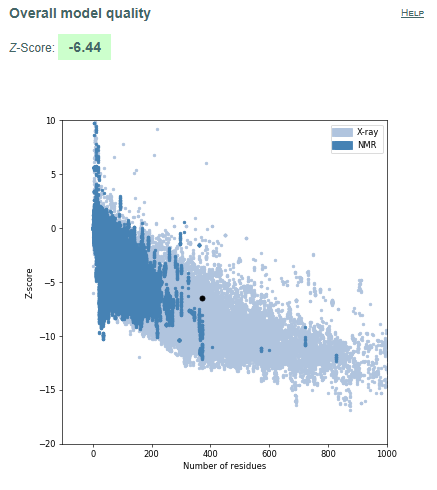

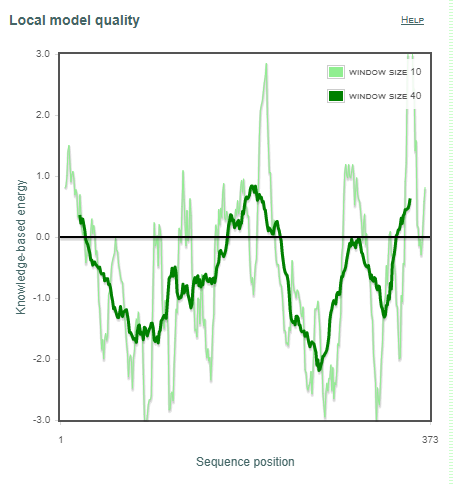

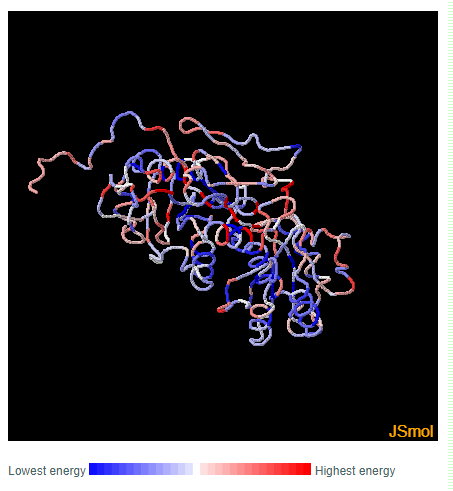

**a	b	c**

**Figure 20.** Validation of model protein structure of asnB of *Streptomyces venezuelae1* **(a)** ProSA-web z-scores of all protein chains in PDB determined by X-ray crystallography (light blue) or NMR spectroscopy (dark blue) with respect to their length. Black dot in the plot indicates that the model protein structure falls inside the range of plot that contains the z-score of all the experimentally determined protein in the PDB. The plot shows only chains with less than 1000 residues and a z-score 10. The z-scores of model proteins are highlighted as large dots. **(b)** Energy plot of model protein which indicates the local model quality by plotting energy as the function of amino acid sequence. Generally, the portion in the positive region of the plot indicates the erroneous part of the structure **(c)** Residues are colored from blue to red in the order of increasing residue energy.

**RESULT:**
*Streptomyces venezuelae* (Fig.19) plot shows that 83.9% of residues in most favoured regions, 12.5% in additional allowed regions, 1.6% residues in generously allowed regions and 2.0% residues in disallowed regions. More than 99% residues are in allowed region given by Ramachandran plot indicates a very good model. Furthermore, the Ramachandran Z-score calculated by WHATCHECK -1.410 falls on the accepted region(Sousa et al., 2006) and allowed by the WHATCHECK. The structures were finally validated using ProSA-web server. This server gives the z-score which indicates the overall model quality and measures the deviation of the total the total energy of the structure with respect to an energy distribution derived from random conformations[16].The z-score given by the server -6.44 falls inside the range of plot (black dot) that contains the z-score of all the experimentally determined protein in the PDB (X-ray, NMR) (Fig. 20a). In the energy plot (Figs. 20b) which indicates the local model quality by plotting energy as the function of amino acid sequence. Generally, the portion in the positive region of the plot indicates the erroneous part of the structure. We can conclude from the plot that the structure in feasible or accepted as overall residues energies fall under the negative part of the plot. The colored 3D structure of the proteins (Figs. 20c,) shows that the portion in red color is of high energy and portion with the blue color are of low energy[48].

*Azotobacter vinelandi*

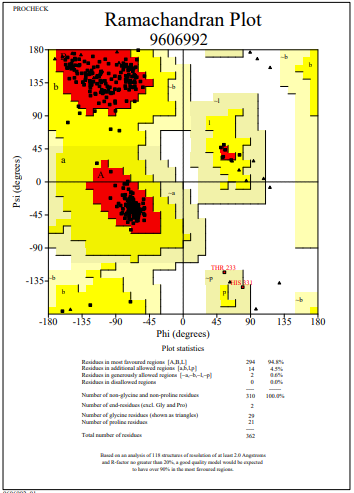

**Figure 21**: Ramachandran Plot of model protein asnB of *Azotobacter vinelandi.* The Ramachandran plot shows the phi-psi torsion angles for all residues (black cubic box) in the structure (except those at the chain termini). Glycine residues are separately identified by triangles as these are not restricted to the regions of the plot appropriate to the other side chain types. The darkest red area indicates the "core" regions representing the most favourable combinations of phi-psi values. The regions are labelled as follows: **A** -Core alpha , **L**	- Core left-handed alpha , **a**	- Allowed alpha , **l**	- Allowed left-handed alpha , **~a** - Generous alpha , **~l**	- Generous left-handed alpha, **B**	- Core beta ,**p**	- Allowed epsilon , **b** - Allowed beta ,**~p** - Generous epsilon , **~b** - Generous beta.

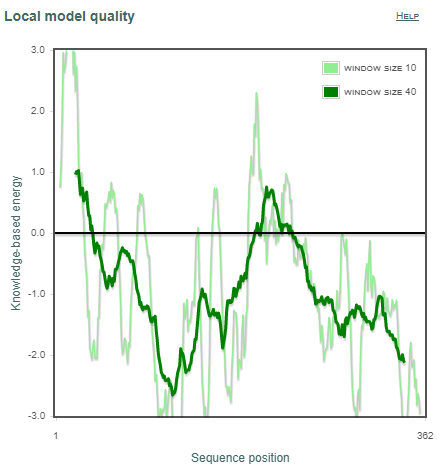

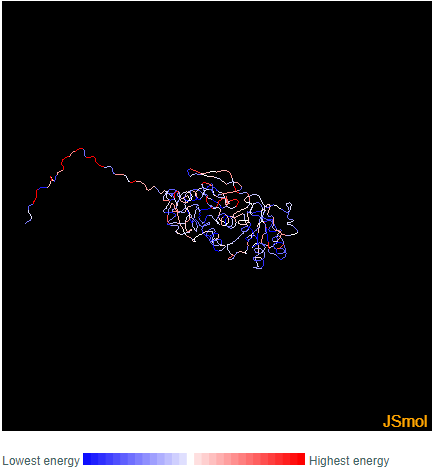

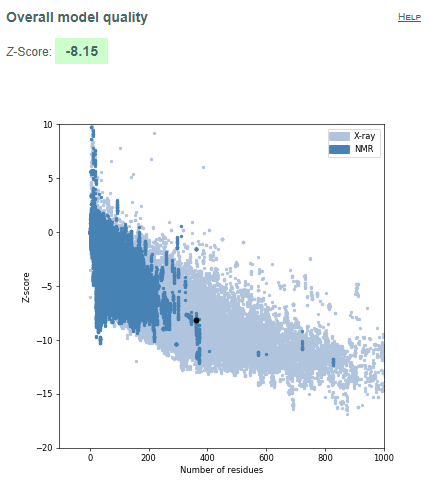

**a	b	c**

**Figure 22.** Validation of model protein structure of asnB of *Azotobacter vinelandi* **(a)** ProSA-web z-scores of all protein chains in PDB determined by X-ray crystallography (light blue) or NMR spectroscopy (dark blue) with respect to their length. Black dot in the plot indicates that the model protein structure falls inside the range of plot that contains the z-score of all the experimentally determined protein in the PDB. The plot shows only chains with less than 1000 residues and a z-score 10. The z-scores of model proteins are highlighted as large dots. **(b)** Energy plot of model protein which indicates the local model quality by plotting energy as the function of amino acid sequence. Generally, the portion in the positive region of the plot indicates the erroneous part of the structure **(c)** Residues are coloured from blue to red in the order of increasing residue energy.

**RESULT:**
*Azotobacter vinelandi* (Fig.21) plot shows that 94.8% of residues in most favoured regions, 4.5% in additional allowed regions, 0.6% residues in generously allowed regions and 0.0% residues in disallowed regions. More than 99% residues are in allowed region given by Ramachandran plot indicates a very good model. Furthermore, the Ramachandran Z-score calculated by WHATCHECK -0.313 falls on the accepted region(Sousa et al., 2006) and allowed by the WHATCHECK. The structures were finally validated using ProSA-web server. This server gives the z-score which indicates the overall model quality and measures the deviation of the total the total energy of the structure with respect to an energy distribution derived from random conformations[16].The z-score given by the server -8.15 falls inside the range of plot (black dot) that contains the z-score of all the experimentally determined protein in the PDB (X-ray, NMR) (Fig. 22a). In the energy plot (Figs. 22b) which indicates the local model quality by plotting energy as the function of amino acid sequence. Generally, the portion in the positive region of the plot indicates the erroneous part of the structure. We can conclude from the plot that the structure in feasible or accepted as overall residues energies fall under the negative part of the plot. The colored 3D structure of the proteins (Figs. 22c,) shows that the portion in red color is of high energy and portion with the blue color are of low energy[48].

*Bacillus licheniformis 1*

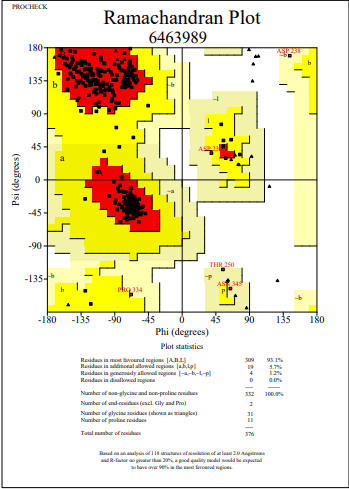

**Figure 23**: Ramachandran Plot of model protein asnB of *Bacillus licheniformis 1.* The Ramachandran plot shows the phi-psi torsion angles for all residues (black cubic box) in the structure (except those at the chain termini). Glycine residues are separately identified by triangles as these are not restricted to the regions of the plot appropriate to the other side chain types. The darkest red area indicates the "core" regions representing the most favourable combinations of phi-psi values. The regions are labelled as follows: **A** -Core alpha , **L**	- Core left-handed alpha , **a**	- Allowed alpha , **l**	- Allowed left-handed alpha , **~a** - Generous alpha , **~l**	- Generous left-handed alpha, **B**	- Core beta ,**p**	- Allowed epsilon , **b** - Allowed beta ,**~p** - Generous epsilon , **~b** - Generous beta.

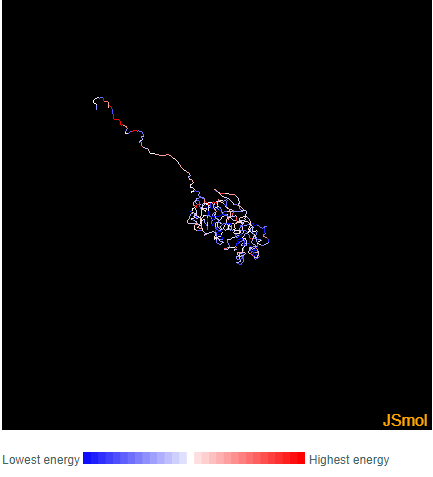

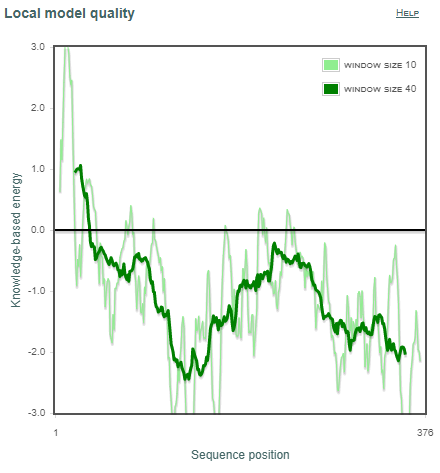

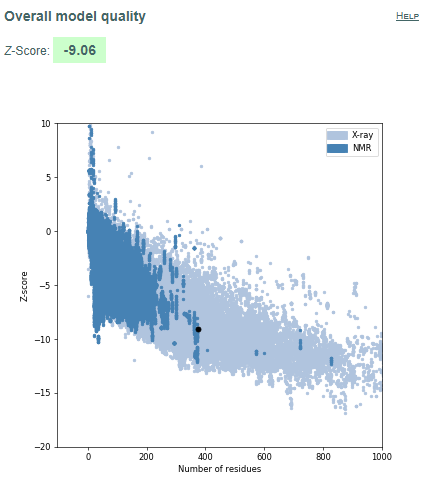

**a	b	c**

**Figure 24.** Validation of model protein structure of asnB of *Bacillus licheniformis 1* **(a)** ProSA-web z-scores of all protein chains in PDB determined by X-ray crystallography (light blue) or NMR spectroscopy (dark blue) with respect to their length. Black dot in the plot indicates that the model protein structure falls inside the range of plot that contains the z-score of all the experimentally determined protein in the PDB. The plot shows only chains with less than 1000 residues and a z-score 10. The z-scores of model proteins are highlighted as large dots. **(b)** Energy plot of model protein which indicates the local model quality by plotting energy as the function of amino acid sequence. Generally, the portion in the positive region of the plot indicates the erroneous part of the structure **(c)** Residues are coloured from blue to red in the order of increasing residue energy.

**RESULT:**
*Bacillus licheniformis 1* (Fig.23) plot shows that 93.1% of residues in most favoured regions, 5.7% in additional allowed regions, 1.2% residues in generously allowed regions and 0.0% residues in disallowed regions. More than 99% residues are in allowed region given by Ramachandran plot indicates a very good model. Furthermore, the Ramachandran Z-score calculated by WHATCHECK -0.436 falls on the accepted region(Sousa et al., 2006) and allowed by the WHATCHECK. The structures were finally validated using ProSA-web server. This server gives the z-score which indicates the overall model quality and measures the deviation of the total the total energy of the structure with respect to an energy distribution derived from random conformations[16].The z-score given by the server -9.06 falls inside the range of plot (black dot) that contains the z-score of all the experimentally determined protein in the PDB (X-ray, NMR) (Fig. 24a). In the energy plot (Figs. 24b) which indicates the local model quality by plotting energy as the function of amino acid sequence. Generally, the portion in the positive region of the plot indicates the erroneous part of the structure. We can conclude from the plot that the structure in feasible or accepted as overall residues energies fall under the negative part of the plot. The colored 3D structure of the proteins (Figs. 24c,) shows that the portion in red color is of high energy and portion with the blue color are of low energy[48].

*Bacillus licheniformis 2*

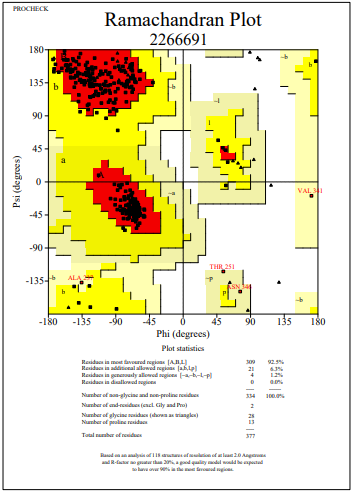

**Figure 25**: Ramachandran Plot of model protein asnB of *Bacillus licheniformis 2.* The Ramachandran plot shows the phi-psi torsion angles for all residues (black cubic box) in the structure (except those at the chain termini). Glycine residues are separately identified by triangles as these are not restricted to the regions of the plot appropriate to the other side chain types. The darkest red area indicates the "core" regions representing the most favourable combinations of phi-psi values. The regions are labelled as follows: **A** -Core alpha , **L**	- Core left-handed alpha , **a**	- Allowed alpha , **l**	- Allowed left-handed alpha , **~a** - Generous alpha , **~l**	- Generous left-handed alpha, **B**	- Core beta ,**p**	- Allowed epsilon , **b** - Allowed beta ,**~p** - Generous epsilon , **~b** - Generous beta.

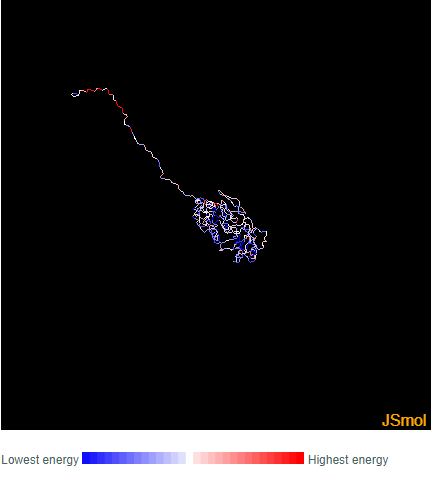

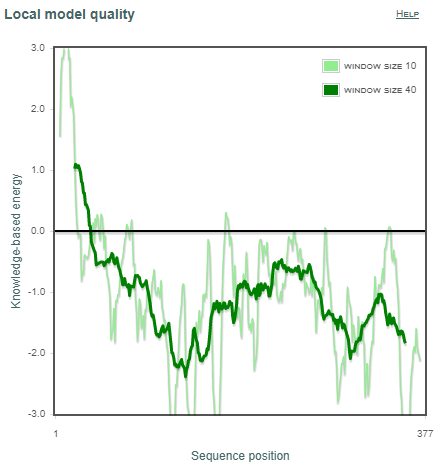

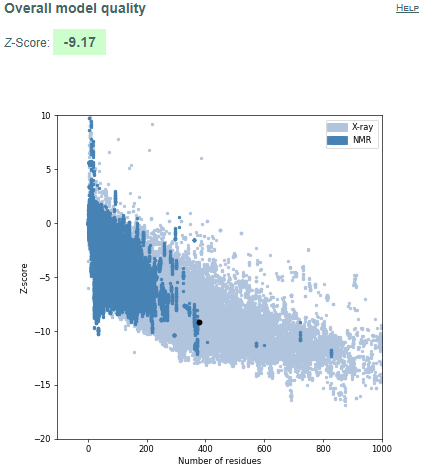

**a	b	c**

**Figure 26.** Validation of model protein structure of asnB of *Bacillus licheniformis 2* **(a)** ProSA-web z-scores of all protein chains in PDB determined by X-ray crystallography (light blue) or NMR spectroscopy (dark blue) with respect to their length. Black dot in the plot indicates that the model protein structure falls inside the range of plot that contains the z-score of all the experimentally determined protein in the PDB. The plot shows only chains with less than 1000 residues and a z-score 10. The z-scores of model proteins are highlighted as large dots. **(b)** Energy plot of model protein which indicates the local model quality by plotting energy as the function of amino acid sequence. Generally, the portion in the positive region of the plot indicates the erroneous part of the structure **(c)** Residues are coloured from blue to red in the order of increasing residue energy.

**RESULT:**
*Bacillus licheniformis 2* (Fig.25) plot shows that 92.5% of residues in most favoured regions, 6.3% in additional allowed regions, 1.2% residues in generously allowed regions and 0.0% residues in disallowed regions. More than 99% residues are in allowed region given by Ramachandran plot indicates a very good model. Furthermore, the Ramachandran Z-score calculated by WHATCHECK -0.947 falls on the accepted region(Sousa et al., 2006) and allowed by the WHATCHECK. The structures were finally validated using ProSA-web server. This server gives the z-score which indicates the overall model quality and measures the deviation of the total the total energy of the structure with respect to an energy distribution derived from random conformations[16].The z-score given by the server -9.17 falls inside the range of plot (black dot) that contains the z-score of all the experimentally determined protein in the PDB (X-ray, NMR) (Fig. 26a). In the energy plot (Figs. 26b) which indicates the local model quality by plotting energy as the function of amino acid sequence. Generally, the portion in the positive region of the plot indicates the erroneous part of the structure. We can conclude from the plot that the structure in feasible or accepted as overall residues energies fall under the negative part of the plot. The colored 3D structure of the proteins (Figs. 26c,) shows that the portion in red color is of high energy and portion with the blue color are of low energy[48].

*Bacillus subtilis 1*

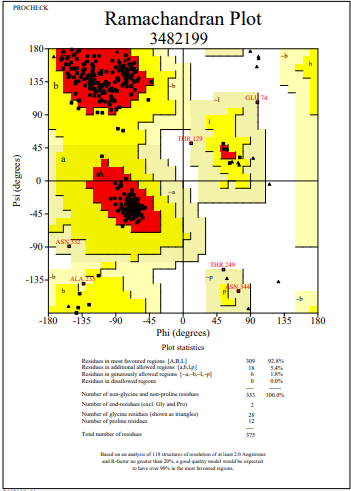

**Figure 27**: Ramachandran Plot of model protein asnB of *Bacillus subtilis 1.* The Ramachandran plot shows the phi-psi torsion angles for all residues (black cubic box) in the structure (except those at the chain termini). Glycine residues are separately identified by triangles as these are not restricted to the regions of the plot appropriate to the other side chain types. The darkest red area indicates the "core" regions representing the most favourable combinations of phi-psi values. The regions are labelled as follows: **A** -Core alpha , **L**	- Core left-handed alpha , **a**	- Allowed alpha , **l**	- Allowed left-handed alpha , **~a** - Generous alpha , **~l**	- Generous left-handed alpha, **B**	- Core beta ,**p**	- Allowed epsilon , **b** - Allowed beta ,**~p** - Generous epsilon , **~b** - Generous beta.

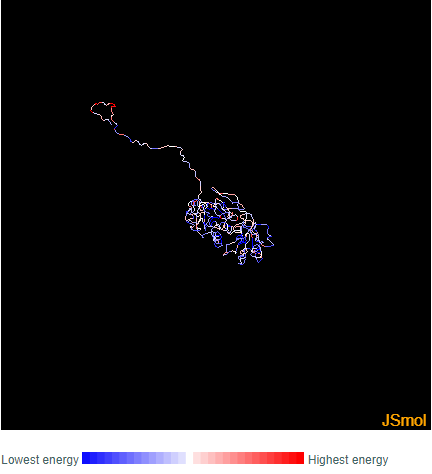

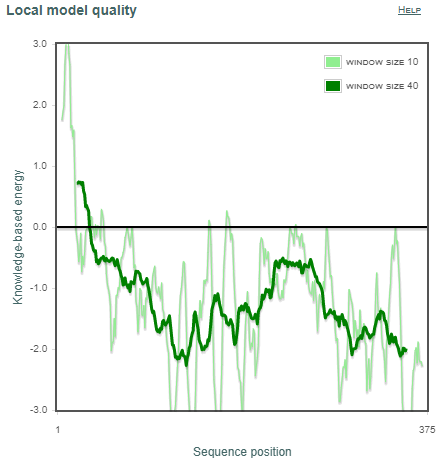

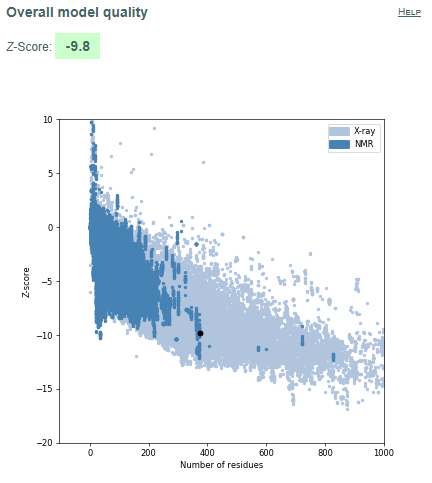

**a	b	c**

**Figure 28.** Validation of model protein structure of asnB of *Bacillus subtilis 1* **(a)** ProSA-web z-scores of all protein chains in PDB determined by X-ray crystallography (light blue) or NMR spectroscopy (dark blue) with respect to their length. Black dot in the plot indicates that the model protein structure falls inside the range of plot that contains the z-score of all the experimentally determined protein in the PDB. The plot shows only chains with less than 1000 residues and a z-score 10. The z-scores of model proteins are highlighted as large dots. **(b)** Energy plot of model protein which indicates the local model quality by plotting energy as the function of amino acid sequence. Generally, the portion in the positive region of the plot indicates the erroneous part of the structure **(c)** Residues are coloured from blue to red in the order of increasing residue energy.

**RESULT:**
*Bacillus subtilis 1* (Fig.27) plot shows that 92.8% of residues in most favoured regions, 5.4% in additional allowed regions, 1.8% residues in generously allowed regions and 0.0% residues in disallowed regions. More than 99% residues are in allowed region given by Ramachandran plot indicates a very good model. Furthermore, the Ramachandran Z-score calculated by WHATCHECK -0.911 falls on the accepted region(Sousa et al., 2006) and allowed by the WHATCHECK. The structures were finally validated using ProSA-web server. This server gives the z-score which indicates the overall model quality and measures the deviation of the total the total energy of the structure with respect to an energy distribution derived from random conformations(Hooft et al., 1997).The z-score given by the server -9.8 falls inside the range of plot (black dot) that contains the z-score of all the experimentally determined protein in the PDB (X-ray, NMR) (Fig. 28a). In the energy plot (Figs. 28b) which indicates the local model quality by plotting energy as the function of amino acid sequence. Generally, the portion in the positive region of the plot indicates the erroneous part of the structure. We can conclude from the plot that the structure in feasible or accepted as overall residues energies fall under the negative part of the plot. The colored 3D structure of the proteins (Figs. 28c,) shows that the portion in red color is of high energy and portion with the blue color are of low energy(Wiederstein & Sippl, 2007).

*Bacillus subtilis 2*

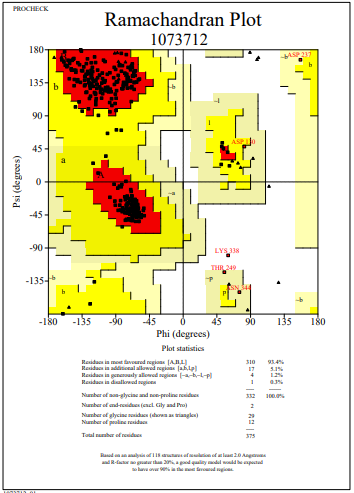

**Figure 29**: Ramachandran Plot of model protein asnB of *Bacillus subtilis 2.* The Ramachandran plot shows the phi-psi torsion angles for all residues (black cubic box) in the structure (except those at the chain termini). Glycine residues are separately identified by triangles as these are not restricted to the regions of the plot appropriate to the other side chain types. The darkest red area indicates the "core" regions representing the most favourable combinations of phi-psi values. The regions are labelled as follows: **A** -Core alpha , **L**	- Core left-handed alpha , **a**	- Allowed alpha , **l**	- Allowed left-handed alpha , **~a** - Generous alpha , **~l**	- Generous left-handed alpha, **B**	- Core beta ,**p**	- Allowed epsilon , **b** - Allowed beta ,**~p** - Generous epsilon , **~b** - Generous beta.

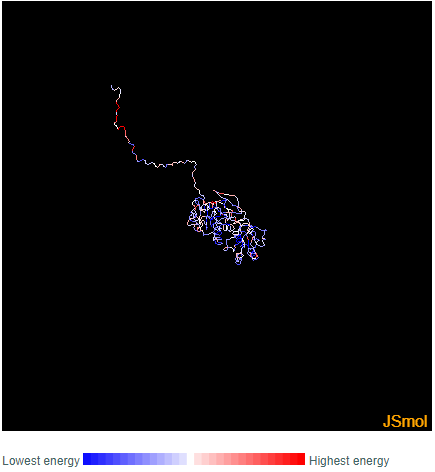

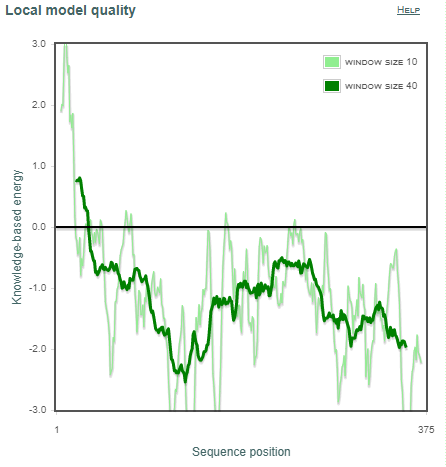

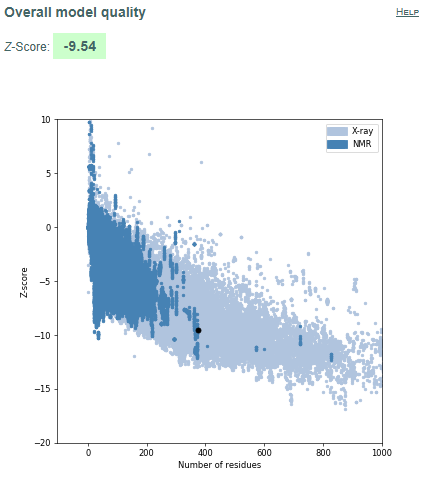

**a	b	c**

**Figure 30.** Validation of model protein structure of asnB of *Bacillus subtilis 2* **(a)** ProSA-web z-scores of all protein chains in PDB determined by X-ray crystallography (light blue) or NMR spectroscopy (dark blue) with respect to their length. Black dot in the plot indicates that the model protein structure falls inside the range of plot that contains the z-score of all the experimentally determined protein in the PDB. The plot shows only chains with less than 1000 residues and a z-score 10. The z-scores of model proteins are highlighted as large dots. **(b)** Energy plot of model protein which indicates the local model quality by plotting energy as the function of amino acid sequence. Generally, the portion in the positive region of the plot indicates the erroneous part of the structure **(c)** Residues are coloured from blue to red in the order of increasing residue energy.

**RESULT:**
*Bacillus subtilis 2* (Fig.29) plot shows that 93.4% of residues in most favoured regions, 5.1% in additional allowed regions, 1.2% residues in generously allowed regions and 0.3% residues in disallowed regions. More than 99% residues are in allowed region given by Ramachandran plot indicates a very good model. Furthermore, the Ramachandran Z-score calculated by WHATCHECK -0.652 falls on the accepted region(Sousa et al., 2006) and allowed by the WHATCHECK. The structures were finally validated using ProSA-web server. This server gives the z-score which indicates the overall model quality and measures the deviation of the total the total energy of the structure with respect to an energy distribution derived from random conformations(Hooft et al., 1997).The z-score given by the server -9.54 falls inside the range of plot (black dot) that contains the z-score of all the experimentally determined protein in the PDB (X-ray, NMR) (Fig. 30a). In the energy plot (Figs. 30b) which indicates the local model quality by plotting energy as the function of amino acid sequence. Generally, the portion in the positive region of the plot indicates the erroneous part of the structure. We can conclude from the plot that the structure in feasible or accepted as overall residues energies fall under the negative part of the plot. The colored 3D structure of the proteins (Figs. 30c,) shows that the portion in red color is of high energy and portion with the blue color are of low energy(Wiederstein & Sippl, 2007).

*Bacilus aryabhattai*

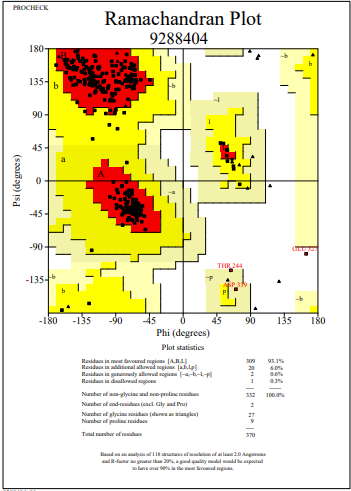

**Figure 31**: Ramachandran Plot of model protein asnB of *Bacilus aryabhattai.* The Ramachandran plot shows the phi-psi torsion angles for all residues (black cubic box) in the structure (except those at the chain termini). Glycine residues are separately identified by triangles as these are not restricted to the regions of the plot appropriate to the other side chain types. The darkest red area indicates the "core" regions representing the most favourable combinations of phi-psi values. The regions are labelled as follows: **A** -Core alpha , **L**	- Core left-handed alpha , **a**	- Allowed alpha , **l**	- Allowed left-handed alpha , **~a** - Generous alpha , **~l**	- Generous left-handed alpha, **B**	- Core beta ,**p**	- Allowed epsilon , **b** - Allowed beta ,**~p** - Generous epsilon , **~b** - Generous beta.

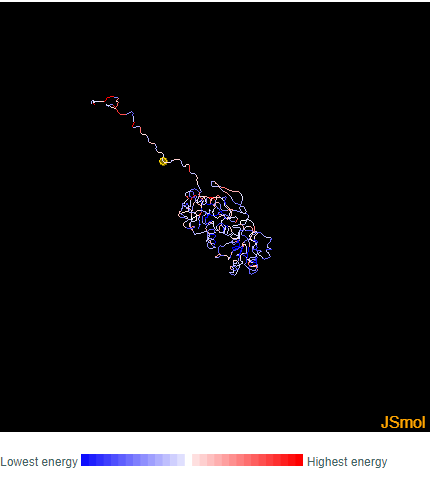

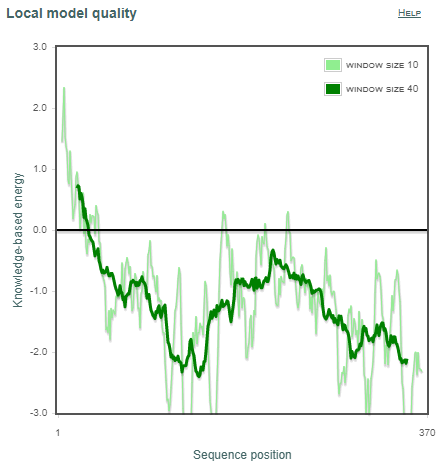

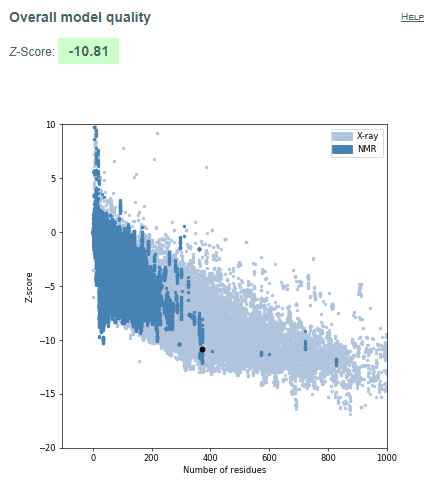

**a	b	c**

**Figure 32.** Validation of model protein structure of asnB of *Bacilus aryabhattai* **(a)** ProSA-web z-scores of all protein chains in PDB determined by X-ray crystallography (light blue) or NMR spectroscopy (dark blue) with respect to their length. Black dot in the plot indicates that the model protein structure falls inside the range of plot that contains the z-score of all the experimentally determined protein in the PDB. The plot shows only chains with less than 1000 residues and a z-score 10. The z-scores of model proteins are highlighted as large dots. **(b)** Energy plot of model protein which indicates the local model quality by plotting energy as the function of amino acid sequence. Generally, the portion in the positive region of the plot indicates the erroneous part of the structure **(c)** Residues are coloured from blue to red in the order of increasing residue energy.

**RESULT:**
*Bacilus aryabhattai* (Fig.31) plot shows that 93.1% of residues in most favoured regions, 6.0% in additional allowed regions, 0.6% residues in generously allowed regions and 0.3% residues in disallowed regions. More than 99% residues are in allowed region given by Ramachandran plot indicates a very good model. Furthermore, the Ramachandran Z-score calculated by WHATCHECK -0.453 falls on the accepted region(Sousa et al., 2006) and allowed by the WHATCHECK. The structures were finally validated using ProSA-web server. This server gives the z-score which indicates the overall model quality and measures the deviation of the total the total energy of the structure with respect to an energy distribution derived from random conformations(Hooft et al., 1997).The z-score given by the server -10.81 falls inside the range of plot (black dot) that contains the z-score of all the experimentally determined protein in the PDB (X-ray, NMR) (Fig. 32a). In the energy plot (Figs. 32b) which indicates the local model quality by plotting energy as the function of amino acid sequence. Generally, the portion in the positive region of the plot indicates the erroneous part of the structure. We can conclude from the plot that the structure in feasible or accepted as overall residues energies fall under the negative part of the plot. The colored 3D structure of the proteins (Figs. 32c,) shows that the portion in red color is of high energy and portion with the blue color are of low energy(Wiederstein & Sippl, 2007)

.

*Delftia acidovorans 1*

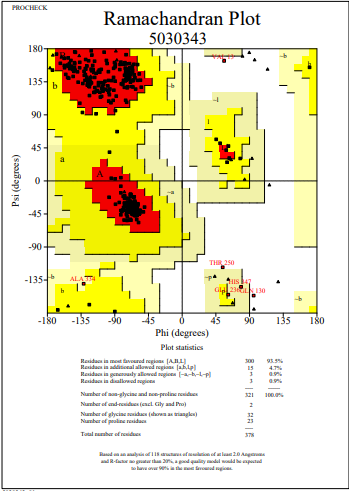

**Figure 33**: Ramachandran Plot of model protein asnB of *Delftia acidovorans 1.* The Ramachandran plot shows the phi-psi torsion angles for all residues (black cubic box) in the structure (except those at the chain termini). Glycine residues are separately identified by triangles as these are not restricted to the regions of the plot appropriate to the other side chain types. The darkest red area indicates the "core" regions representing the most favourable combinations of phi-psi values. The regions are labelled as follows: **A** -Core alpha , **L**	- Core left-handed alpha , **a**	- Allowed alpha , **l**	- Allowed left-handed alpha , **~a** - Generous alpha , **~l**	- Generous left-handed alpha, **B**	- Core beta ,**p**	- Allowed epsilon , **b** - Allowed beta ,**~p** - Generous epsilon , **~b** - Generous beta.

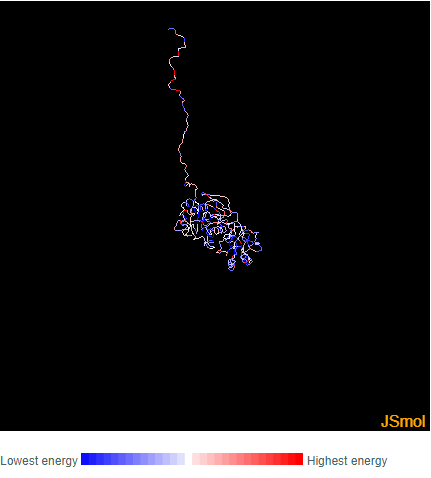

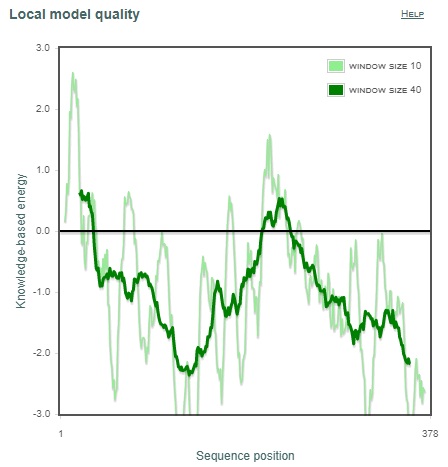

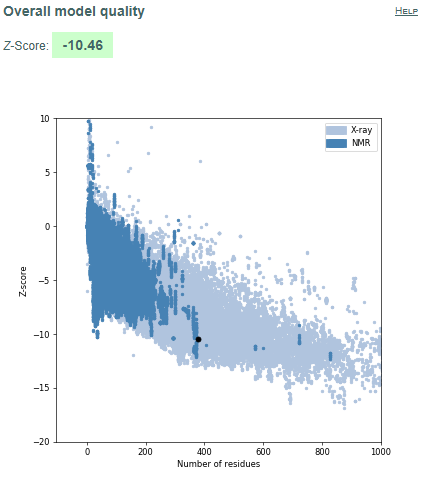

**a	b	c**

**Figure 34.** Validation of model protein structure of asnB of *Delftia acidovorans 1* **(a)** ProSA-web z-scores of all protein chains in PDB determined by X-ray crystallography (light blue) or NMR spectroscopy (dark blue) with respect to their length. Black dot in the plot indicates that the model protein structure falls inside the range of plot that contains the z-score of all the experimentally determined protein in the PDB. The plot shows only chains with less than 1000 residues and a z-score 10. The z-scores of model proteins are highlighted as large dots. **(b)** Energy plot of model protein which indicates the local model quality by plotting energy as the function of amino acid sequence. Generally, the portion in the positive region of the plot indicates the erroneous part of the structure **(c)** Residues are coloured from blue to red in the order of increasing residue energy.

**RESULT:**
*Delftia acidovorans 1* (Fig.33) plot shows that 93.5% of residues in most favoured regions, 4.7% in additional allowed regions, 0.9% residues in generously allowed regions and 0.9% residues in disallowed regions. More than 99% residues are in allowed region given by Ramachandran plot indicates a very good model. Furthermore, the Ramachandran Z-score calculated by WHATCHECK -0.385 falls on the accepted region(Sousa et al., 2006) and allowed by the WHATCHECK. The structures were finally validated using ProSA-web server. This server gives the z-score which indicates the overall model quality and measures the deviation of the total the total energy of the structure with respect to an energy distribution derived from random conformations(Hooft et al., 1997).The z-score given by the server -10.46 falls inside the range of plot (black dot) that contains the z-score of all the experimentally determined protein in the PDB (X-ray, NMR) (Fig. 34a). In the energy plot (Figs. 34b) which indicates the local model quality by plotting energy as the function of amino acid sequence. Generally, the portion in the positive region of the plot indicates the erroneous part of the structure. We can conclude from the plot that the structure in feasible or accepted as overall residues energies fall under the negative part of the plot. The colored 3D structure of the proteins (Figs. 34c,) shows that the portion in red colour is of high energy and portion with the blue colour are of low energy(Wiederstein & Sippl, 2007).

*Delftia acidovorans 2*

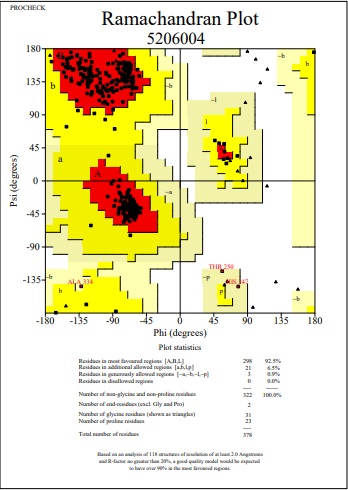

**Figure 35**: Ramachandran Plot of model protein asnB of *Delftia acidovorans 2.* The Ramachandran plot shows the phi-psi torsion angles for all residues (black cubic box) in the structure (except those at the chain termini). Glycine residues are separately identified by triangles as these are not restricted to the regions of the plot appropriate to the other side chain types. The darkest red area indicates the "core" regions representing the most favourable combinations of phi-psi values. The regions are labelled as follows: **A** -Core alpha , **L**	- Core left-handed alpha , **a**	- Allowed alpha , **l**	- Allowed left-handed alpha , **~a** - Generous alpha , **~l**	- Generous left-handed alpha, **B**	- Core beta ,**p**	- Allowed epsilon , **b** - Allowed beta ,**~p** - Generous epsilon , **~b** - Generous beta.

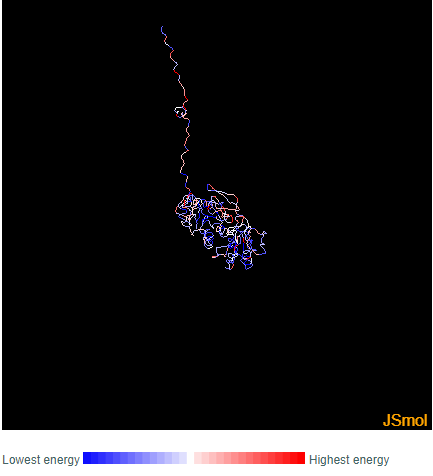

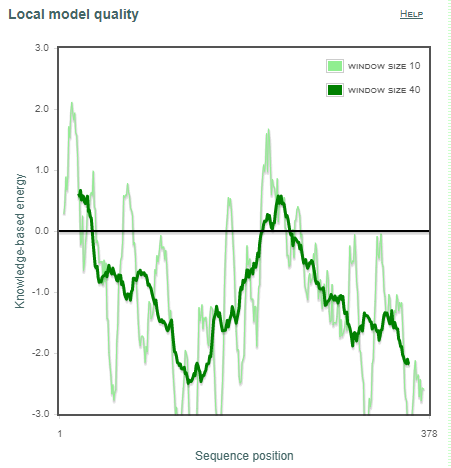

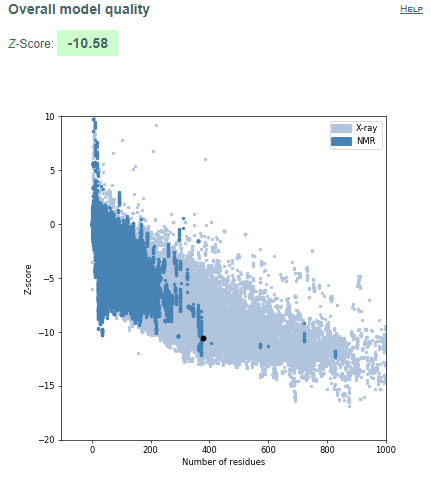

**a	b	c**

**Figure 36.** Validation of model protein structure of asnB of *Delftia acidovorans 2* **(a)** ProSA-web z-scores of all protein chains in PDB determined by X-ray crystallography (light blue) or NMR spectroscopy (dark blue) with respect to their length. Black dot in the plot indicates that the model protein structure falls inside the range of plot that contains the z-score of all the experimentally determined protein in the PDB. The plot shows only chains with less than 1000 residues and a z-score 10. The z-scores of model proteins are highlighted as large dots. **(b)** Energy plot of model protein which indicates the local model quality by plotting energy as the function of amino acid sequence. Generally, the portion in the positive region of the plot indicates the erroneous part of the structure **(c)** Residues are coloured from blue to red in the order of increasing residue energy.

**RESULT:**
*Delftia acidovorans 2* (Fig.35) plot shows that 92.5% of residues in most favoured regions, 6.5% in additional allowed regions, 0.9% residues in generously allowed regions and 0.0% residues in disallowed regions. More than 99% residues are in allowed region given by Ramachandran plot indicates a very good model. Furthermore, the Ramachandran Z-score calculated by WHATCHECK -0.391 falls on the accepted region(Sousa et al., 2006) and allowed by the WHATCHECK. The structures were finally validated using ProSA-web server. This server gives the z-score which indicates the overall model quality and measures the deviation of the total the total energy of the structure with respect to an energy distribution derived from random conformations(Hooft et al., 1997).The z-score given by the server -10.58 falls inside the range of plot (black dot) that contains the z-score of all the experimentally determined protein in the PDB (X-ray, NMR) (Fig. 36a). In the energy plot (Figs. 36b) which indicates the local model quality by plotting energy as the function of amino acid sequence. Generally, the portion in the positive region of the plot indicates the erroneous part of the structure. We can conclude from the plot that the structure in feasible or accepted as overall residues energies fall under the negative part of the plot. The colored 3D structure of the proteins (Figs. 36c,) shows that the portion in red colour is of high energy and portion with the blue colour are of low energy(Wiederstein & Sippl, 2007).

*Dickeya chrysanthami 1*

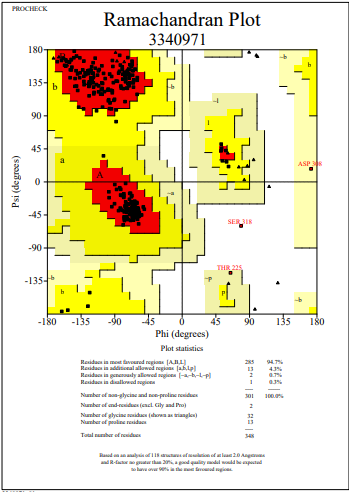

**Figure 37**: Ramachandran Plot of model protein asnB of *Dickeya chrysanthami 1.* The Ramachandran plot shows the phi-psi torsion angles for all residues (black cubic box) in the structure (except those at the chain termini). Glycine residues are separately identified by triangles as these are not restricted to the regions of the plot appropriate to the other side chain types. The darkest red area indicates the "core" regions representing the most favourable combinations of phi-psi values. The regions are labelled as follows: **A** -Core alpha , **L**	- Core left-handed alpha , **a**	- Allowed alpha , **l**	- Allowed left-handed alpha , **~a** - Generous alpha , **~l**	- Generous left-handed alpha, **B**	- Core beta ,**p**	- Allowed epsilon , **b** - Allowed beta ,**~p** - Generous epsilon , **~b** - Generous beta.

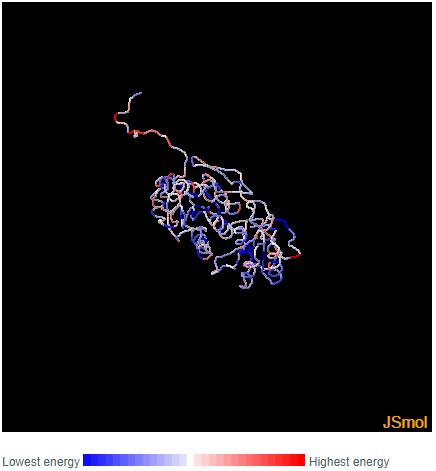

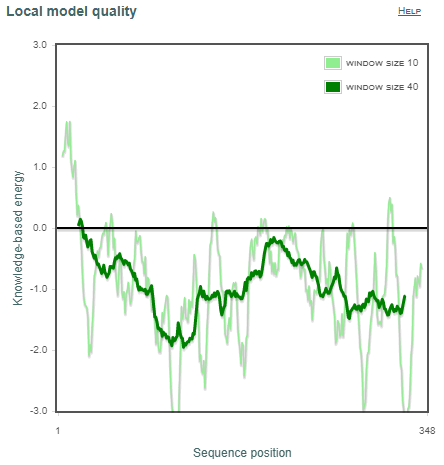

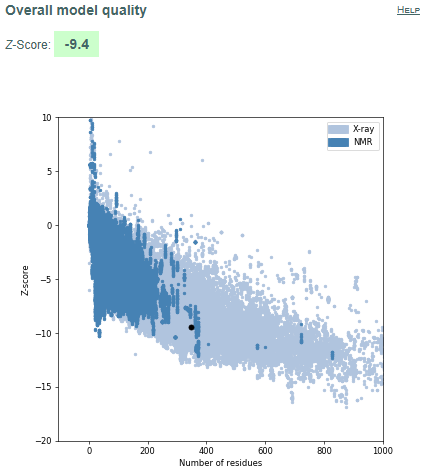

**a	b	c**

**Figure 38.** Validation of model protein structure of asnB of *Dickeya chrysanthami 1* **(a)** ProSA-web z-scores of all protein chains in PDB determined by X-ray crystallography (light blue) or NMR spectroscopy (dark blue) with respect to their length. Black dot in the plot indicates that the model protein structure falls inside the range of plot that contains the z-score of all the experimentally determined protein in the PDB. The plot shows only chains with less than 1000 residues and a z-score 10. The z-scores of model proteins are highlighted as large dots. **(b)** Energy plot of model protein which indicates the local model quality by plotting energy as the function of amino acid sequence. Generally, the portion in the positive region of the plot indicates the erroneous part of the structure **(c)** Residues are coloured from blue to red in the order of increasing residue energy.

**RESULT:**
*Dickeya chrysanthami 1* (Fig.37) plot shows that 94.7% of residues in most favoured regions, 4.3% in additional allowed regions, 0.7% residues in generously allowed regions and 0.3% residues in disallowed regions. More than 99% residues are in allowed region given by Ramachandran plot indicates a very good model. Furthermore, the Ramachandran Z-score calculated by WHATCHECK -0.125 falls on the accepted region(Sousa et al., 2006) and allowed by the WHATCHECK. The structures were finally validated using ProSA-web server. This server gives the z-score which indicates the overall model quality and measures the deviation of the total the total energy of the structure with respect to an energy distribution derived from random conformations(Hooft et al., 1997).The z-score given by the server -9.4 falls inside the range of plot (black dot) that contains the z-score of all the experimentally determined protein in the PDB (X-ray, NMR) (Fig. 38a). In the energy plot (Figs. 38b) which indicates the local model quality by plotting energy as the function of amino acid sequence. Generally, the portion in the positive region of the plot indicates the erroneous part of the structure. We can conclude from the plot that the structure in feasible or accepted as overall residues energies fall under the negative part of the plot. The colored 3D structure of the proteins (Figs. 38c,) shows that the portion in red colour is of high energy and portion with the blue colour are of low energy(Wiederstein & Sippl, 2007)

.

*Dickeya chrysanthami 2*

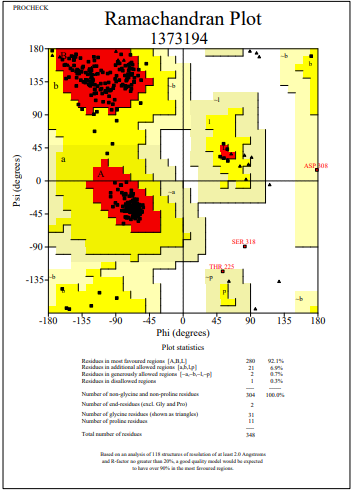

**Figure 39**: Ramachandran Plot of model protein asnB of *Dickeya chrysanthami 2.* The Ramachandran plot shows the phi-psi torsion angles for all residues (black cubic box) in the structure (except those at the chain termini). Glycine residues are separately identified by triangles as these are not restricted to the regions of the plot appropriate to the other side chain types. The darkest red area indicates the "core" regions representing the most favourable combinations of phi-psi values. The regions are labelled as follows: **A** -Core alpha , **L**	- Core left-handed alpha , **a**	- Allowed alpha , **l**	- Allowed left-handed alpha , **~a** - Generous alpha , **~l**	- Generous left-handed alpha, **B**	- Core beta ,**p**	- Allowed epsilon , **b** - Allowed beta ,**~p** - Generous epsilon , **~b** - Generous beta.

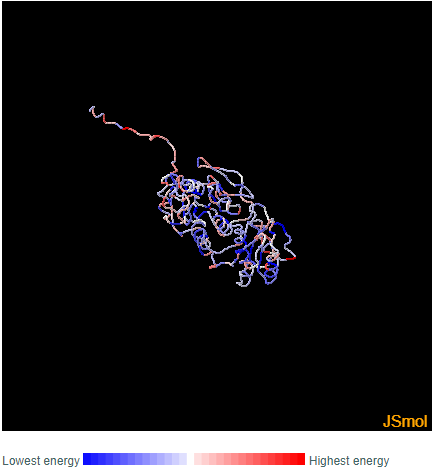

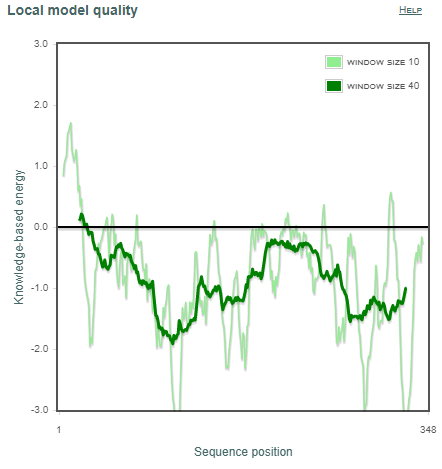

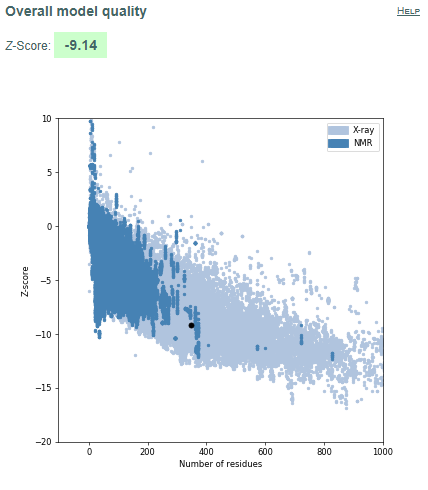

**a	b	c**

**Figure 40.** Validation of model protein structure of asnB of *Dickeya chrysanthami 2* **(a)** ProSA-web z-scores of all protein chains in PDB determined by X-ray crystallography (light blue) or NMR spectroscopy (dark blue) with respect to their length. Black dot in the plot indicates that the model protein structure falls inside the range of plot that contains the z-score of all the experimentally determined protein in the PDB. The plot shows only chains with less than 1000 residues and a z-score 10. The z-scores of model proteins are highlighted as large dots. **(b)** Energy plot of model protein which indicates the local model quality by plotting energy as the function of amino acid sequence. Generally, the portion in the positive region of the plot indicates the erroneous part of the structure **(c)** Residues are coloured from blue to red in the order of increasing residue energy.

**RESULT:**
*Dickeya chrysanthami 2* (Fig.39) plot shows that 92.1% of residues in most favoured regions, 6.9% in additional allowed regions, 0.7% residues in generously allowed regions and 0.3% residues in disallowed regions. More than 99% residues are in allowed region given by Ramachandran plot indicates a very good model. Furthermore, the Ramachandran Z-score calculated by WHATCHECK -0.609 falls on the accepted region(Sousa et al., 2006) and allowed by the WHATCHECK. The structures were finally validated using ProSA-web server. This server gives the z-score which indicates the overall model quality and measures the deviation of the total the total energy of the structure with respect to an energy distribution derived from random conformations(Hooft et al., 1997).The z-score given by the server -9.14 falls inside the range of plot (black dot) that contains the z-score of all the experimentally determined protein in the PDB (X-ray, NMR) (Fig. 40a). In the energy plot (Figs. 40b) which indicates the local model quality by plotting energy as the function of amino acid sequence. Generally, the portion in the positive region of the plot indicates the erroneous part of the structure. We can conclude from the plot that the structure in feasible or accepted as overall residues energies fall under the negative part of the plot. The colored 3D structure of the proteins (Figs. 40c,) shows that the portion in red colour is of high energy and portion with the blue colour are of low energy(Wiederstein & Sippl, 2007).

*Helicobacter pylori 1*

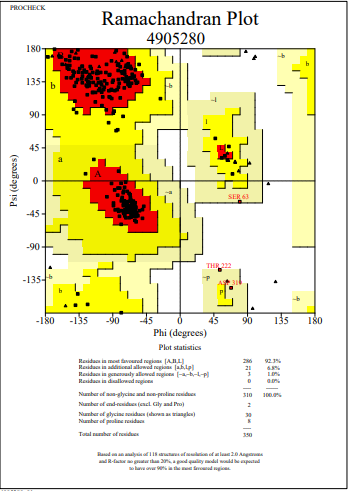

**Figure 41**: Ramachandran Plot of model protein asnB of *Helicobacter pylori 1.* The Ramachandran plot shows the phi-psi torsion angles for all residues (black cubic box) in the structure (except those at the chain termini). Glycine residues are separately identified by triangles as these are not restricted to the regions of the plot appropriate to the other side chain types. The darkest red area indicates the "core" regions representing the most favourable combinations of phi-psi values. The regions are labelled as follows: **A** -Core alpha , **L**	- Core left-handed alpha , **a**	- Allowed alpha , **l**	- Allowed left-handed alpha , **~a** - Generous alpha , **~l**	- Generous left-handed alpha, **B**	- Core beta ,**p**	- Allowed epsilon , **b** - Allowed beta ,**~p** - Generous epsilon , **~b** - Generous beta.

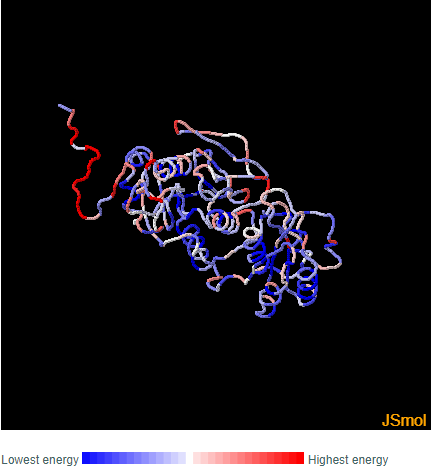

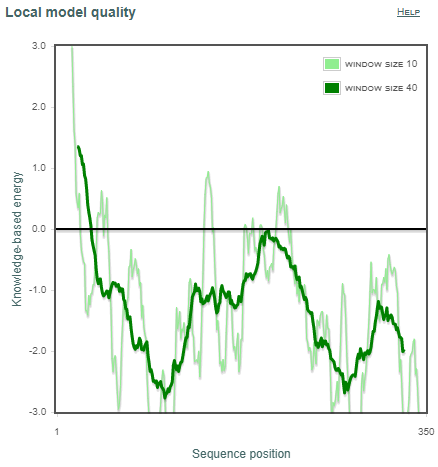

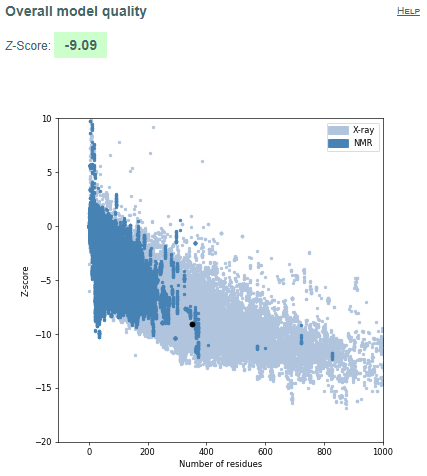

**a	b	c**

**Figure 42.** Validation of model protein structure of asnB of *Helicobacter pylori 1* **(a)** ProSA-web z-scores of all protein chains in PDB determined by X-ray crystallography (light blue) or NMR spectroscopy (dark blue) with respect to their length. Black dot in the plot indicates that the model protein structure falls inside the range of plot that contains the z-score of all the experimentally determined protein in the PDB. The plot shows only chains with less than 1000 residues and a z-score 10. The z-scores of model proteins are highlighted as large dots. **(b)** Energy plot of model protein which indicates the local model quality by plotting energy as the function of amino acid sequence. Generally, the portion in the positive region of the plot indicates the erroneous part of the structure **(c)** Residues are coloured from blue to red in the order of increasing residue energy.

**RESULT:**
*Helicobacter pylori 1* (Fig.41) plot shows that 92.3% of residues in most favoured regions, 6.8% in additional allowed regions, 1.0% residues in generously allowed regions and 0.0% residues in disallowed regions. More than 99% residues are in allowed region given by Ramachandran plot indicates a very good model. Furthermore, the Ramachandran Z-score calculated by WHATCHECK -0.360 falls on the accepted region(Sousa et al., 2006) and allowed by the WHATCHECK. The structures were finally validated using ProSA-web server. This server gives the z-score which indicates the overall model quality and measures the deviation of the total the total energy of the structure with respect to an energy distribution derived from random conformations(Hooft et al., 1997).The z-score given by the server -9.09 falls inside the range of plot (black dot) that contains the z-score of all the experimentally determined protein in the PDB (X-ray, NMR) (Fig. 42a). In the energy plot (Figs. 42b) which indicates the local model quality by plotting energy as the function of amino acid sequence. Generally, the portion in the positive region of the plot indicates the erroneous part of the structure. We can conclude from the plot that the structure in feasible or accepted as overall residues energies fall under the negative part of the plot. The colored 3D structure of the proteins (Figs. 42c,) shows that the portion in red colour is of high energy and portion with the blue colour are of low energy(Wiederstein & Sippl, 2007).

*Helicobacter pylori 2*

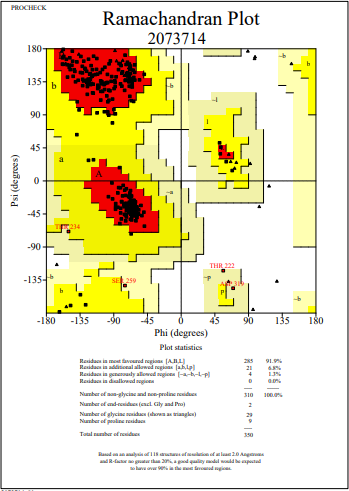

**Figure 43**: Ramachandran Plot of model protein asnB of *Helicobacter pylori 2.* The Ramachandran plot shows the phi-psi torsion angles for all residues (black cubic box) in the structure (except those at the chain termini). Glycine residues are separately identified by triangles as these are not restricted to the regions of the plot appropriate to the other side chain types. The darkest red area indicates the "core" regions representing the most favourable combinations of phi-psi values. The regions are labelled as follows: **A** -Core alpha , **L**	- Core left-handed alpha , **a**	- Allowed alpha , **l**	- Allowed left-handed alpha , **~a** - Generous alpha , **~l**	- Generous left-handed alpha, **B**	- Core beta ,**p**	- Allowed epsilon , **b** - Allowed beta ,**~p** - Generous epsilon , **~b** - Generous beta.

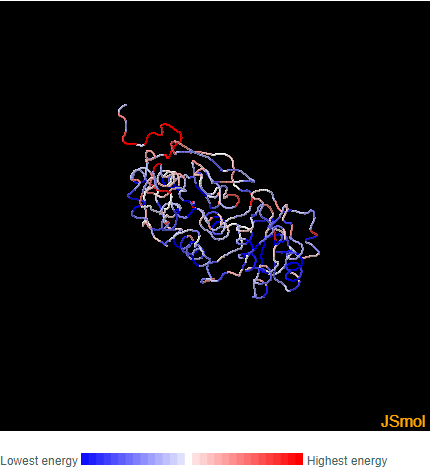

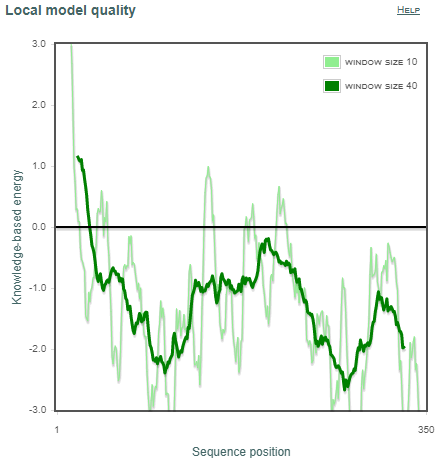

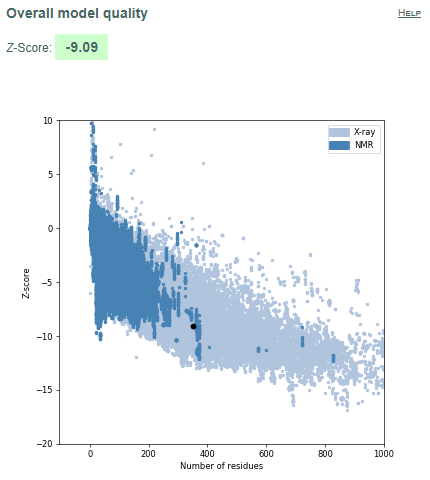

**a	b	c**

**Figure 44.** Validation of model protein structure of asnB of *Helicobacter pylori 2* **(a)** ProSA-web z-scores of all protein chains in PDB determined by X-ray crystallography (light blue) or NMR spectroscopy (dark blue) with respect to their length. Black dot in the plot indicates that the model protein structure falls inside the range of plot that contains the z-score of all the experimentally determined protein in the PDB. The plot shows only chains with less than 1000 residues and a z-score 10. The z-scores of model proteins are highlighted as large dots. **(b)** Energy plot of model protein which indicates the local model quality by plotting energy as the function of amino acid sequence. Generally, the portion in the positive region of the plot indicates the erroneous part of the structure **(c)** Residues are coloured from blue to red in the order of increasing residue energy.

**RESULT:**
*Helicobacter pylori 2* (Fig.43) plot shows that 91.9% of residues in most favoured regions, 6.8% in additional allowed regions, 1.3% residues in generously allowed regions and 0.0% residues in disallowed regions. More than 99% residues are in allowed region given by Ramachandran plot indicates a very good model. Furthermore, the Ramachandran Z-score calculated by WHATCHECK -0.309 falls on the accepted region(Sousa et al., 2006) and allowed by the WHATCHECK. The structures were finally validated using ProSA-web server. This server gives the z-score which indicates the overall model quality and measures the deviation of the total the total energy of the structure with respect to an energy distribution derived from random conformations(Hooft et al., 1997).The z-score given by the server -9.09 falls inside the range of plot (black dot) that contains the z-score of all the experimentally determined protein in the PDB (X-ray, NMR) (Fig. 44a). In the energy plot (Figs. 44b) which indicates the local model quality by plotting energy as the function of amino acid sequence. Generally, the portion in the positive region of the plot indicates the erroneous part of the structure. We can conclude from the plot that the structure in feasible or accepted as overall residues energies fall under the negative part of the plot. The colored 3D structure of the proteins (Figs. 44c,) shows that the portion in red colour is of high energy and portion with the blue colour are of low energy(Wiederstein & Sippl, 2007).

*Pectobacterium carotovorum 1*

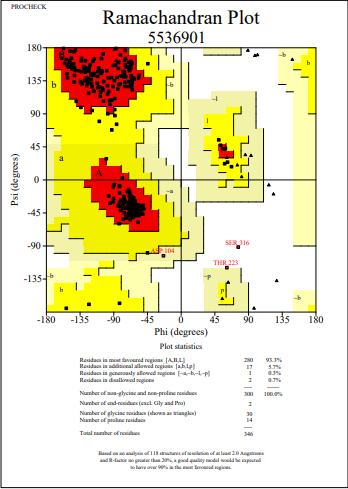

**Figure 45**: Ramachandran Plot of model protein asnB of *Pectobacterium carotovorum 1.* The Ramachandran plot shows the phi-psi torsion angles for all residues (black cubic box) in the structure (except those at the chain termini). Glycine residues are separately identified by triangles as these are not restricted to the regions of the plot appropriate to the other side chain types. The darkest red area indicates the "core" regions representing the most favourable combinations of phi-psi values. The regions are labelled as follows: **A** -Core alpha , **L**	- Core left-handed alpha , **a**	- Allowed alpha , **l**	- Allowed left-handed alpha , **~a** - Generous alpha , **~l**	- Generous left-handed alpha, **B**	- Core beta ,**p**	- Allowed epsilon , **b** - Allowed beta ,**~p** - Generous epsilon , **~b** - Generous beta.

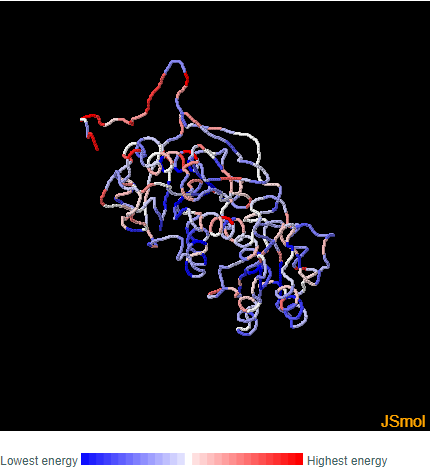

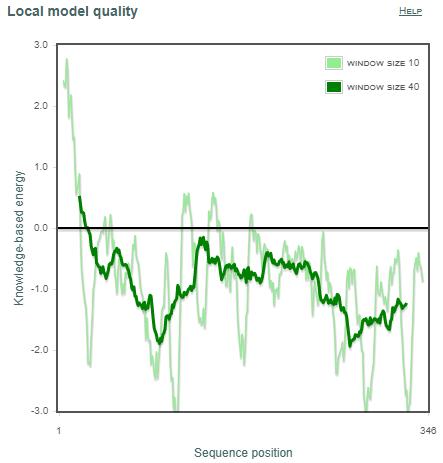

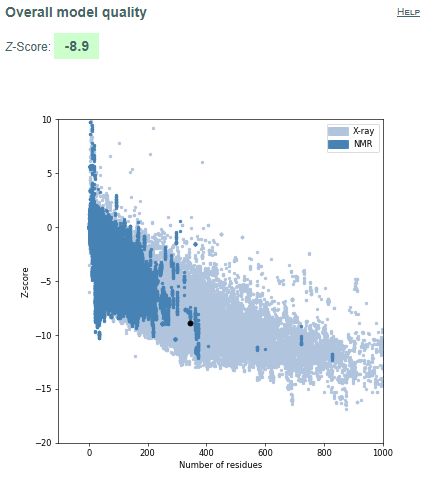

**a	b	c**

**Figure 46.** Validation of model protein structure of asnB of *Pectobacterium carotovorum 1* **(a)** ProSA-web z-scores of all protein chains in PDB determined by X-ray crystallography (light blue) or NMR spectroscopy (dark blue) with respect to their length. Black dot in the plot indicates that the model protein structure falls inside the range of plot that contains the z-score of all the experimentally determined protein in the PDB. The plot shows only chains with less than 1000 residues and a z-score 10. The z-scores of model proteins are highlighted as large dots. **(b)** Energy plot of model protein which indicates the local model quality by plotting energy as the function of amino acid sequence. Generally, the portion in the positive region of the plot indicates the erroneous part of the structure **(c)** Residues are coloured from blue to red in the order of increasing residue energy.

**RESULT:**
*Pectobacterium carotovorum 1* (Fig.45) plot shows that 93.3% of residues in most favoured regions, 5.7% in additional allowed regions, 0.3% residues in generously allowed regions and 0.7% residues in disallowed regions. More than 99% residues are in allowed region given by Ramachandran plot indicates a very good model. Furthermore, the Ramachandran Z-score calculated by WHATCHECK -0.358 falls on the accepted region(Sousa et al., 2006) and allowed by the WHATCHECK. The structures were finally validated using ProSA-web server. This server gives the z-score which indicates the overall model quality and measures the deviation of the total the total energy of the structure with respect to an energy distribution derived from random conformations(Hooft et al., 1997).The z-score given by the server -8.9 falls inside the range of plot (black dot) that contains the z-score of all the experimentally determined protein in the PDB (X-ray, NMR) (Fig. 46a). In the energy plot (Figs. 46b) which indicates the local model quality by plotting energy as the function of amino acid sequence. Generally, the portion in the positive region of the plot indicates the erroneous part of the structure. We can conclude from the plot that the structure in feasible or accepted as overall residues energies fall under the negative part of the plot. The colored 3D structure of the proteins (Figs. 46c,) shows that the portion in red colour is of high energy and portion with the blue colour are of low energy(Wiederstein & Sippl, 2007).

*Pectobacterium carotovorum 2*

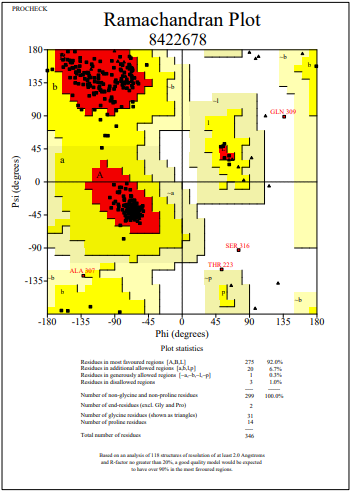

**Figure 47**: Ramachandran Plot of model protein asnB of *Pectobacterium carotovorum 2.* The Ramachandran plot shows the phi-psi torsion angles for all residues (black cubic box) in the structure (except those at the chain termini). Glycine residues are separately identified by triangles as these are not restricted to the regions of the plot appropriate to the other side chain types. The darkest red area indicates the "core" regions representing the most favourable combinations of phi-psi values. The regions are labelled as follows: **A** -Core alpha , **L**	- Core left-handed alpha , **a**	- Allowed alpha , **l**	- Allowed left-handed alpha , **~a** - Generous alpha , **~l**	- Generous left-handed alpha, **B**	- Core beta ,**p**	- Allowed epsilon , **b** - Allowed beta ,**~p** - Generous epsilon , **~b** - Generous beta.

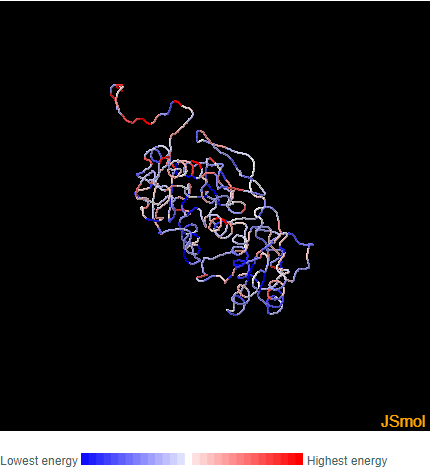

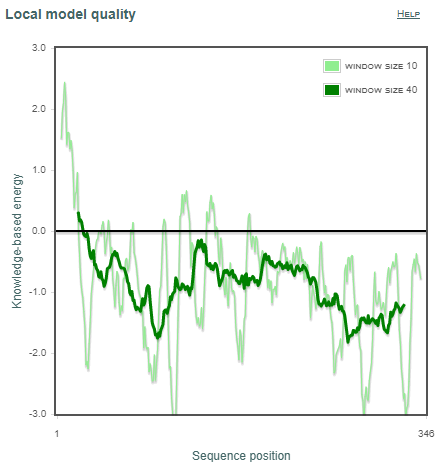

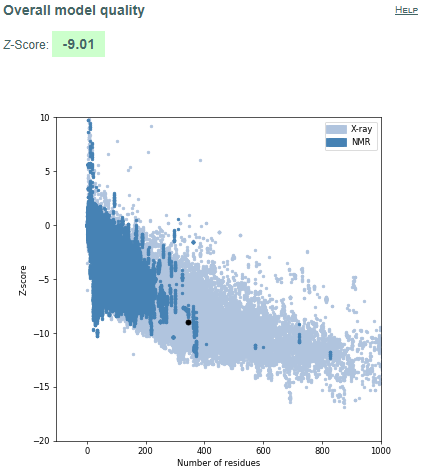

**a	b	c**

**Figure 48.** Validation of model protein structure of asnB of *Pectobacterium carotovorum 2* **(a)** ProSA-web z-scores of all protein chains in PDB determined by X-ray crystallography (light blue) or NMR spectroscopy (dark blue) with respect to their length. Black dot in the plot indicates that the model protein structure falls inside the range of plot that contains the z-score of all the experimentally determined protein in the PDB. The plot shows only chains with less than 1000 residues and a z-score 10. The z-scores of model proteins are highlighted as large dots. **(b)** Energy plot of model protein which indicates the local model quality by plotting energy as the function of amino acid sequence. Generally, the portion in the positive region of the plot indicates the erroneous part of the structure **(c)** Residues are coloured from blue to red in the order of increasing residue energy.

**RESULT:**
*Pectobacterium carotovorum 2* (Fig.47) plot shows that 92.0% of residues in most favoured regions, 6.7% in additional allowed regions, 0.3% residues in generously allowed regions and 1.0% residues in disallowed regions. More than 99% residues are in allowed region given by Ramachandran plot indicates a very good model. Furthermore, the Ramachandran Z-score calculated by WHATCHECK -0.642 falls on the accepted region(Sousa et al., 2006) and allowed by the WHATCHECK. The structures were finally validated using ProSA-web server. This server gives the z-score which indicates the overall model quality and measures the deviation of the total the total energy of the structure with respect to an energy distribution derived from random conformations(Hooft et al., 1997).The z-score given by the server -9.01 falls inside the range of plot (black dot) that contains the z-score of all the experimentally determined protein in the PDB (X-ray, NMR) (Fig. 48a). In the energy plot (Figs. 48b) which indicates the local model quality by plotting energy as the function of amino acid sequence. Generally, the portion in the positive region of the plot indicates the erroneous part of the structure. We can conclude from the plot that the structure in feasible or accepted as overall residues energies fall under the negative part of the plot. The colored 3D structure of the proteins (Figs. 48c,) shows that the portion in red colour is of high energy and portion with the blue colour are of low energy(Wiederstein & Sippl, 2007).

*Pseudomonas fluorescens*

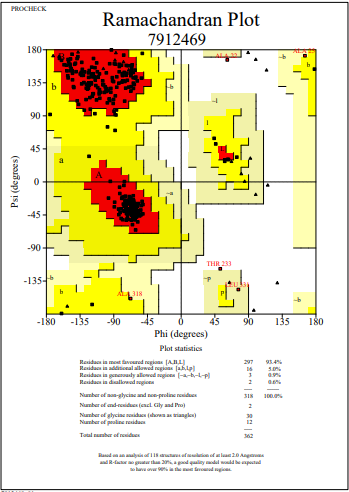

**Figure 49**: Ramachandran Plot of model protein asnB of *Pseudomonas fluorescens.* The Ramachandran plot shows the phi-psi torsion angles for all residues (black cubic box) in the structure (except those at the chain termini). Glycine residues are separately identified by triangles as these are not restricted to the regions of the plot appropriate to the other side chain types. The darkest red area indicates the "core" regions representing the most favourable combinations of phi-psi values. The regions are labelled as follows: **A** -Core alpha , **L**	- Core left-handed alpha , **a**	- Allowed alpha , **l**	- Allowed left-handed alpha , **~a** - Generous alpha , **~l**	- Generous left-handed alpha, **B**	- Core beta ,**p**	- Allowed epsilon , **b** - Allowed beta ,**~p** - Generous epsilon , **~b** - Generous beta.

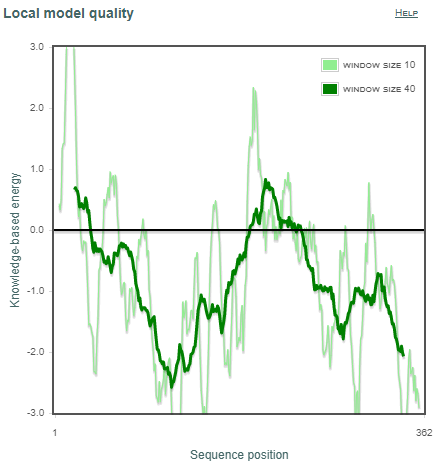

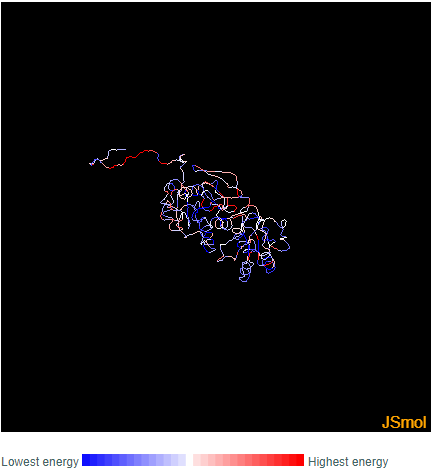

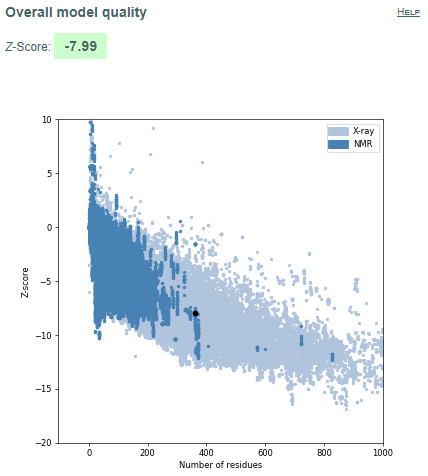

**a	b	c**

**Figure 50.** Validation of model protein structure of asnB of *Pseudomonas fluorescens* **(a)** ProSA-web z-scores of all protein chains in PDB determined by X-ray crystallography (light blue) or NMR spectroscopy (dark blue) with respect to their length. Black dot in the plot indicates that the model protein structure falls inside the range of plot that contains the z-score of all the experimentally determined protein in the PDB. The plot shows only chains with less than 1000 residues and a z-score 10. The z-scores of model proteins are highlighted as large dots. **(b)** Energy plot of model protein which indicates the local model quality by plotting energy as the function of amino acid sequence. Generally, the portion in the positive region of the plot indicates the erroneous part of the structure **(c)** Residues are coloured from blue to red in the order of increasing residue energy.

**RESULT:**
*Pseudomonas fluorescens* (Fig.49) plot shows that 93.4% of residues in most favoured regions, 5.0% in additional allowed regions, 0.9% residues in generously allowed regions and 0.6% residues in disallowed regions. More than 99% residues are in allowed region given by Ramachandran plot indicates a very good model. Furthermore, the Ramachandran Z-score calculated by WHATCHECK -0.470 falls on the accepted region(Sousa et al., 2006) and allowed by the WHATCHECK. The structures were finally validated using ProSA-web server. This server gives the z-score which indicates the overall model quality and measures the deviation of the total the total energy of the structure with respect to an energy distribution derived from random conformations(Hooft et al., 1997).The z-score given by the server -7.99 falls inside the range of plot (black dot) that contains the z-score of all the experimentally determined protein in the PDB (X-ray, NMR) (Fig. 50a). In the energy plot (Figs. 50b) which indicates the local model quality by plotting energy as the function of amino acid sequence. Generally, the portion in the positive region of the plot indicates the erroneous part of the structure. We can conclude from the plot that the structure in feasible or accepted as overall residues energies fall under the negative part of the plot. The colored 3D structure of the proteins (Figs. 50c,) shows that the portion in red colour is of high energy and portion with the blue colour are of low energy(Wiederstein & Sippl, 2007).

*Pseudomonas strutzeri 1*

**Figure 51**: Ramachandran Plot of model protein asnB of *Pseudomonas strutzeri 1.* The Ramachandran plot shows the phi-psi torsion angles for all residues (black cubic box) in the structure (except those at the chain termini). Glycine residues are separately identified by triangles as these are not restricted to the regions of the plot appropriate to the other side chain types. The darkest red area indicates the "core" regions representing the most favourable combinations of phi-psi values. The regions are labelled as follows: **A** -Core alpha , **L**	- Core left-handed alpha , **a**	- Allowed alpha , **l**	- Allowed left-handed alpha , **~a** - Generous alpha , **~l**	- Generous left-handed alpha, **B**	- Core beta ,**p**	- Allowed epsilon , **b** - Allowed beta ,**~p** - Generous epsilon , **~b** - Generous beta.

**a	b	c**

**Figure 52.** Validation of model protein structure of asnB of *Pseudomonas strutzeri 1* **(a)** ProSA-web z-scores of all protein chains in PDB determined by X-ray crystallography (light blue) or NMR spectroscopy (dark blue) with respect to their length. Black dot in the plot indicates that the model protein structure falls inside the range of plot that contains the z-score of all the experimentally determined protein in the PDB. The plot shows only chains with less than 1000 residues and a z-score 10. The z-scores of model proteins are highlighted as large dots. **(b)** Energy plot of model protein which indicates the local model quality by plotting energy as the function of amino acid sequence. Generally, the portion in the positive region of the plot indicates the erroneous part of the structure **(c)** Residues are coloured from blue to red in the order of increasing residue energy.

**RESULT:**
*Pseudomonas strutzeri 1* (Fig.51) plot shows that 89.6% of residues in most favoured regions, 8.4% in additional allowed regions, 1.3% residues in generously allowed regions and 0.7% residues in disallowed regions. More than 99% residues are in allowed region given by Ramachandran plot indicates a very good model. Furthermore, the Ramachandran Z-score calculated by WHATCHECK -1.433 falls on the accepted region(Sousa et al., 2006) and allowed by the WHATCHECK. The structures were finally validated using ProSA-web server. This server gives the z-score which indicates the overall model quality and measures the deviation of the total the total energy of the structure with respect to an energy distribution derived from random conformations(Hooft et al., 1997).The z-score given by the server -10.00 falls inside the range of plot (black dot) that contains the z-score of all the experimentally determined protein in the PDB (X-ray, NMR) (Fig. 52a). In the energy plot (Figs. 52b) which indicates the local model quality by plotting energy as the function of amino acid sequence. Generally, the portion in the positive region of the plot indicates the erroneous part of the structure. We can conclude from the plot that the structure in feasible or accepted as overall residues energies fall under the negative part of the plot. The colored 3D structure of the proteins (Figs. 52c,) shows that the portion in red colour is of high energy and portion with the blue colour are of low energy(Wiederstein & Sippl, 2007).

*Pseudomonas strutzeri 2*

**Figure 53**: Ramachandran Plot of model protein asnB of *Pseudomonas strutzeri 2.* The Ramachandran plot shows the phi-psi torsion angles for all residues (black cubic box) in the structure (except those at the chain termini). Glycine residues are separately identified by triangles as these are not restricted to the regions of the plot appropriate to the other side chain types. The darkest red area indicates the "core" regions representing the most favourable combinations of phi-psi values. The regions are labelled as follows: **A** -Core alpha , **L**	- Core left-handed alpha , **a**	- Allowed alpha , **l**	- Allowed left-handed alpha , **~a** - Generous alpha , **~l**	- Generous left-handed alpha, **B**	- Core beta ,**p**	- Allowed epsilon , **b** - Allowed beta ,**~p** - Generous epsilon , **~b** - Generous beta.

**a	b	c**

**Figure 54.** Validation of model protein structure of asnB of *Pseudomonas strutzeri 2* **(a)** ProSA-web z-scores of all protein chains in PDB determined by X-ray crystallography (light blue) or NMR spectroscopy (dark blue) with respect to their length. Black dot in the plot indicates that the model protein structure falls inside the range of plot that contains the z-score of all the experimentally determined protein in the PDB. The plot shows only chains with less than 1000 residues and a z-score 10. The z-scores of model proteins are highlighted as large dots. **(b)** Energy plot of model protein which indicates the local model quality by plotting energy as the function of amino acid sequence. Generally, the portion in the positive region of the plot indicates the erroneous part of the structure **(c)** Residues are coloured from blue to red in the order of increasing residue energy.

**RESULT:**
*Pseudomonas strutzeri 2* (Fig.53) plot shows that 89.0% of residues in most favoured regions, 9.7% in additional allowed regions, 1.0% residues in generously allowed regions and 0.3% residues in disallowed regions. More than 99% residues are in allowed region given by Ramachandran plot indicates a very good model. Furthermore, the Ramachandran Z-score calculated by WHATCHECK -1.182 falls on the accepted region(Sousa et al., 2006) and allowed by the WHATCHECK. The structures were finally validated using ProSA-web server. This server gives the z-score which indicates the overall model quality and measures the deviation of the total the total energy of the structure with respect to an energy distribution derived from random conformations(Hooft et al., 1997).The z-score given by the server -9.85 falls inside the range of plot (black dot) that contains the z-score of all the experimentally determined protein in the PDB (X-ray, NMR) (Fig. 54a). In the energy plot (Figs. 54b) which indicates the local model quality by plotting energy as the function of amino acid sequence. Generally, the portion in the positive region of the plot indicates the erroneous part of the structure. We can conclude from the plot that the structure in feasible or accepted as overall residues energies fall under the negative part of the plot. The colored 3D structure of the proteins (Figs. 54c,) shows that the portion in red colour is of high energy and portion with the blue colour are of low energy(Wiederstein & Sippl, 2007).

*Escherichia coli*

**Figure 55**: Ramachandran Plot of model protein asnB of *Pseudomonas strutzeri 2.* The Ramachandran plot shows the phi-psi torsion angles for all residues (black cubic box) in the structure (except those at the chain termini). Glycine residues are separately identified by triangles as these are not restricted to the regions of the plot appropriate to the other side chain types. The darkest red area indicates the "core" regions representing the most favourable combinations of phi-psi values. The regions are labelled as follows: **A** -Core alpha , **L**	- Core left-handed alpha , **a**	- Allowed alpha , **l**	- Allowed left-handed alpha , **~a** - Generous alpha , **~l**	- Generous left-handed alpha, **B**	- Core beta ,**p**	- Allowed epsilon , **b** - Allowed beta ,**~p** - Generous epsilon , **~b** - Generous beta

**a	b	c**

**Figure 56.** Validation of model protein structure of asnB of *Pseudomonas strutzeri 2* **(a)** ProSA-web z-scores of all protein chains in PDB determined by X-ray crystallography (light blue) or NMR spectroscopy (dark blue) with respect to their length. Black dot in the plot indicates that the model protein structure falls inside the range of plot that contains the z-score of all the experimentally determined protein in the PDB. The plot shows only chains with less than 1000 residues and a z-score 10. The z-scores of model proteins are highlighted as large dots. **(b)** Energy plot of model protein which indicates the local model quality by plotting energy as the function of amino acid sequence. Generally, the portion in the positive region of the plot indicates the erroneous part of the structure **(c)** Residues are coloured from blue to red in the order of increasing residue energy.

**RESULT:**

*Escherichia coli* (Fig.55) plot shows that 93.4% of residues in most favoured regions, 5.3% in additional allowed regions, 1.3% residues in generously allowed regions and 0.0% residues in disallowed regions. More than 99% residues are in allowed region given by Ramachandran plot indicates a very good model. Furthermore, the Ramachandran Z-score calculated by WHATCHECK -0.124 falls on the accepted region(Sousa et al., 2006) and allowed by the WHATCHECK. The structures were finally validated using ProSA-web server. This server gives the z-score which indicates the overall model quality and measures the deviation of the total the total energy of the structure with respect to an energy distribution derived from random conformations(Hooft et al., 1997).The z-score given by the server -11.31 falls inside the range of plot (black dot) that contains the z-score of all the experimentally determined protein in the PDB (X-ray, NMR) (Fig. 56a). In the energy plot (Figs. 56b) which indicates the local model quality by plotting energy as the function of amino acid sequence. Generally, the portion in the positive region of the plot indicates the erroneous part of the structure. We can conclude from the plot that the structure in feasible or accepted as overall residues energies fall under the negative part of the plot. The colored 3D structure of the proteins (Figs. 56c,) shows that the portion in red colour is of high energy and portion with the blue colour are of low energy(Wiederstein & Sippl, 2007).
